# A reference induced pluripotent stem cell line for large-scale collaborative studies

**DOI:** 10.1101/2021.12.15.472643

**Authors:** Caroline B. Pantazis, Andrian Yang, Erika Lara, Justin A. McDonough, Cornelis Blauwendraat, Lirong Peng, Hideyuki Oguro, Jitendra Kanaujiya, Jizhong Zou, David Sebesta, Gretchen Pratt, Erin Cross, Jeffrey Blockwick, Philip Buxton, Lauren Kinner-Bibeau, Constance Medura, Christopher Tompkins, Stephen Hughes, Marianita Santiana, Faraz Faghri, Mike A. Nalls, Daniel Vitale, Shannon Ballard, Yue A. Qi, Daniel M. Ramos, Kailyn M. Anderson, Julia Stadler, Priyanka Narayan, Jason Papademetriou, Luke Reilly, Matthew P. Nelson, Sanya Aggarwal, Leah U. Rosen, Peter Kirwan, Venkat Pisupati, Steven L. Coon, Sonja W. Scholz, Theresa Priebe, Miriam Öttl, Jian Dong, Marieke Meijer, Lara J.M. Janssen, Vanessa S. Lourenco, Rik van der Kant, Dennis Crusius, Dominik Paquet, Ana-Caroline Raulin, Guojun Bu, Aaron Held, Brian J. Wainger, Rebecca M.C. Gabriele, Jackie M Casey, Selina Wray, Dad Abu-Bonsrah, Clare L. Parish, Melinda S. Beccari, Don W. Cleveland, Emmy Li, Indigo V.L. Rose, Martin Kampmann, Carles Calatayud Aristoy, Patrik Verstreken, Laurin Heinrich, Max Y. Chen, Birgitt Schüle, Dan Dou, Erika L.F. Holzbaur, Maria Clara Zanellati, Richa Basundra, Mohanish Deshmukh, Sarah Cohen, Richa Khanna, Malavika Raman, Zachary S. Nevin, Madeline Matia, Jonas Van Lent, Vincent Timmerman, Bruce R. Conklin, Katherine Johnson Chase, Ke Zhang, Salome Funes, Daryl A. Bosco, Lena Erlebach, Marc Welzer, Deborah Kronenberg-Versteeg, Guochang Lyu, Ernest Arenas, Elena Coccia, Lily Sarrafha, Tim Ahfeldt, John C. Marioni, William C. Skarnes, Mark R. Cookson, Michael E. Ward, Florian T. Merkle

**Affiliations:** Center for Alzheimer’s and Related Dementias (CARD), National Institute on Aging and National Institute of Neurological Disorders and Stroke, National Institutes of Health, Bethesda, MD, USA; European Molecular Biology Laboratory, European Bioinformatics Institute, Wellcome Genome Campus, Hinxton, Cambridge CB10 1SD, UK; Cancer Research UK Cambridge Institute, University of Cambridge, Cambridge, UK; Metabolic Research Laboratories and Medical Research Council Metabolic Diseases Unit, Wellcome Trust - Medical Research Council Institute of Metabolic Science, University of Cambridge, Cambridge CB2 0QQ, UK; Wellcome Trust - Medical Research Council Cambridge Stem Cell Institute, University of Cambridge, Cambridge CB2 0AW, UK; The Jackson Laboratory for Genomic Medicine, Farmington, CT, USA; Laboratory of Neurogenetics, National Institute on Aging, National Institutes of Health, Bethesda, MD, USA; Data Tecnica International LLC, Washington, DC, USA; Integrated Research Facility, National Institute of Allergy and Infectious Diseases, National Institute of Health, Frederick, MD, USA; Department of Cell Biology, University of Connecticut Health Center, Farmington, CT, USA; iPS Cell Core Facility, National Heart, Lung, and Blood Institute, National Institutes of Health, Bethesda, MD, USA; KromaTiD Inc., Longmont, CO, USA; Genetics and Biochemistry Branch, NIDDK, NINDS, National Institutes of Health, Bethesda, MD 20814, USA; John van Geest Centre for Brain Repair, University of Cambridge, Cambridge CB2 0PY, UK; Molecular Genomics Core, Eunice Kennedy Shriver National Institute of Child Health and Human Development, National Institutes of Health, Bethesda, MD, USA; Neurodegenerative Diseases Research Unit, National Institute of Neurological Disorders and Stroke, Bethesda, MD, USA; Department of Neurology, Johns Hopkins University, Baltimore, MD 21287, USA; Department of Functional Genomics, Center for Neurogenomics and Cognitive Research, Amsterdam Neuroscience, VU University Amsterdam de Boelelaan 1087, 1081 HV Amsterdam, the Netherlands; Alzheimer Center Amsterdam, Department of Neurology, Amsterdam Neuroscience, Amsterdam UMC, Amsterdam, Netherlands; Institute for Stroke and Dementia Research, University Hospital, LMU Munich, 81377 Munich, Germany; Munich Cluster for Systems Neurology (SyNergy), 81377 Munich, Germany; Department of Neuroscience, Mayo Clinic, Jacksonville, FL, USA; Department of Neurology, Sean M. Healey & AMG Center for ALS, Massachusetts General Hospital, Harvard Medical School, Boston, MA, USA; Department of Neurology, Sean M. Healey & AMG Center for ALS, Massachusetts General Hospital, Harvard Medical School, Boston, MA, USA; Department of Anesthesiology, Critical Care and Pain Medicine, Massachusetts General Hospital, Boston, MA, USA; Harvard Stem Cell Institute, Cambridge, MA, USA; Broad Institute of Harvard University and MIT, Cambridge, MA, USA; Department of Neurodegenerative Disease, UCL Queen Square Institute of Neurology, London, WC1N 3BG, UK; Florey Institute of Neuroscience and Mental Health, Parkville, VIC 3052, Australia; Department of Cellular and Molecular Medicine, University of California at San Diego, La Jolla, CA, USA; Ludwig Institute for Cancer Research, University of California at San Diego, La Jolla, CA, USA; Institute for Neurodegenerative Diseases, University of California, San Francisco, San Francisco, CA, USA; Chan Zuckerberg Biohub, San Francisco, CA, USA; Department of Biochemistry and Biophysics, University of California, San Francisco, San Francisco, CA, USA; VIB-KU Leuven Center for Brain & Disease Research, 3000 Leuven, Belgium; KU Leuven, Department of Neurosciences, Leuven Brain Institute, Mission Lucidity, Leuven, Belgium; Department of Pathology, Stanford University School of Medicine, Stanford, California, USA; Department of Physiology, Perelman School of Medicine, University of Pennsylvania, Philadelphia, PA, USA; Department of Cell Biology and Physiology, University of North Carolina at Chapel Hill, Chapel Hill, NC, USA; Department of Developmental Molecular and Chemical Biology, Tufts University School of Medicine, Boston, MA, USA; Gladstone Institutes, San Francisco, CA, USA; Peripheral Neuropathy Research Group, Department of Biomedical Sciences, University of Antwerp, Antwerp, 2610, Belgium; Department of Neurology, UMass Chan Medical School, Worcester, MA, USA; Department of Cellular Neurology, Hertie Institute for Clinical Brain Research, University of Tübingen, Tübingen, Germany; German Center for Neurodegenerative Diseases (DZNE), Tübingen, Germany; Division of Molecular Neurobiology, Department of Medical Biochemistry and Biophysics, Karolinska Institutet, Stockholm, Sweden; Nash Family Department of Neuroscience at Mount Sinai, New York, NY, USA; Departments of Neurology and Cell, Developmental and Regenerative Biology at Mount Sinai, New York, NY, USA; Ronald M. Loeb Center for Alzheimer’s Disease at Mount Sinai, New York, NY, USA; Friedman Brain Institute at Mount Sinai, New York, NY, USA; Black Family Stem Cell Institute at Mount Sinai, New York, NY, USA; Wellcome Sanger Institute, Wellcome Genome Campus, Hinxton, UK; National Institute on Neurological Disorders and Stroke, National Institutes of Health, Bethesda, MD, USA; Department of Pediatrics, University of Melbourne, Parkville, VIC 3052, Australia

**Keywords:** iPSC, pluripotent, reference, stem cell line, p53, karyotype, whole-genome, single-cell, CRISPR

## Abstract

Human induced pluripotent stem cell (iPSC) lines are a powerful tool for studying development and disease, but the considerable phenotypic variation between lines makes it challenging to replicate key findings and integrate data across research groups. To address this issue, we sub-cloned candidate iPSC lines and deeply characterised their genetic properties using whole genome sequencing, their genomic stability upon CRISPR/Cas9-based gene editing, and their phenotypic properties including differentiation to commonly-used cell types. These studies identified KOLF2.1J as an all-around well-performing iPSC line. We then shared KOLF2.1J with groups around the world who tested its performance in head-to-head comparisons with their own preferred iPSC lines across a diverse range of differentiation protocols and functional assays. On the strength of these findings, we have made KOLF2.1J and hundreds of its gene-edited derivative clones readily accessible to promote the standardization required for large-scale collaborative science in the stem cell field.

**Summary:** The authors of this collaborative study deeply characterized human induced pluripotent stem cell (iPSC) lines to rationally select a clonally-derived cell line that performs well across multiple modalities. KOLF2.1J was identified as a candidate reference cell line based on single-cell analysis of its gene expression in the pluripotent state, whole genome sequencing, genomic stability after highly efficient CRISPR-mediated gene editing, integrity of the p53 pathway, and the efficiency with which it differentiated into multiple target cell populations. Since it is deeply characterized and can be readily acquired, KOLF2.1J is an attractive reference cell line for groups working with iPSCs.

**Graphical abstract:** 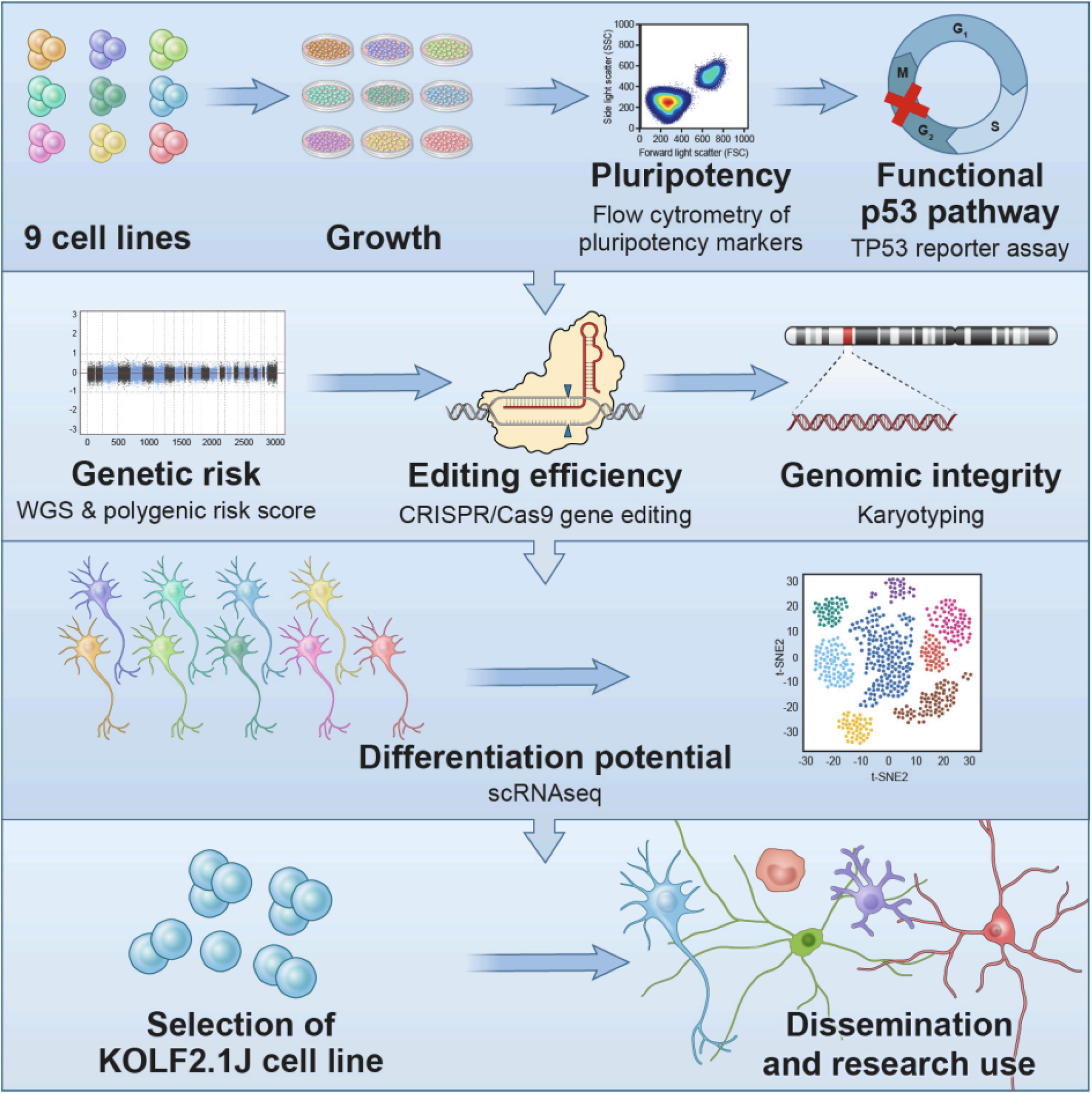

## Introduction

Human iPSCs are increasingly used to model diseases since they capture genetic contributors to disease risk and can be differentiated into relevant cell populations. Additionally, the genomes of iPSCs can be edited to introduce or correct disease-associated variants (Kwart et al. 2019; Konttinen et al. 2019; Guttikonda et al. 2021). For example, comparing control hiPSCs with isogenic knock-in of Mendelian variants associated with Alzheimer’s disease (AD) identified convergent transcriptomic events after differentiation into neurons (Kwart et al. 2019), or non-neuronal cells (Konttinen et al. 2019). In theory, the results of isogenic experiments could be compared across genetic variants, cell types, and analysis modalities by different groups. However, the use of different cell lines by different groups creates an obstacle to data integration, since genetic background influences cellular phenotypes (Doetschman 2009; Bonyadi et al. 1997; Threadgill et al. 1995). This issue has been recognized by communities working with model organisms, who appreciated that the benefits of a common reference outweigh the idiosyncrasies of a particular line or strain, since key results obtained on one genetic background can always be tested on another (Sterken et al. 2015; Mackay and Huang 2018; Sittig et al. 2016). Consequently, there is a large, unmet need in the stem cell field for common, well-characterized cell lines.

Several recent efforts have sought to address these challenges by developing gene-edited iPSC clones from a common parental cell line and making these available to the community. For example, the Allen Cell Collection (https://www.allencell.org/genomics.html) generated a series of publicly-available, gene-edited reporters from the WTC11 iPSC line derived from a healthy donor (Roberts et al. 2017, 2019). Here, we sought to identify a common cell line to facilitate large-scale collaborative studies such as the iPSC Neurodegenerative Disease Initiative (iNDI) from the NIH’s Center for Alzheimer’s and Related Dementias (CARD). The aim of this initiative is to generate hundreds of single nucleotide variant (SNV) knock-in, revertant, gene knockout, and endogenously-tagged CRISPR/Cas9-edited iPSC derivative lines relevant to Alzheimer’s Disease and Related Dementias (ADRD) on well characterized genetic backgrounds (Ramos et al. 2021). To select candidate cell lines for this purpose, we first accounted for the freedom to modify and distribute the line and its derivatives, and then deeply characterized the genomic status, functional characteristics, and differentiation potential of multiple candidate iPSC lines, leading to the identification of KOLF2.1J as a lead reference cell line.

## Results

### Rationale and establishment of clonal candidate cell sub-lines

We set out to identify one or more deeply characterized human pluripotent stem cell lines to serve as a common reference for the field (Ramos et al. 2021; Reilly et al., n.d.). Since human embryonic stem cell (hESC) lines face usage restrictions in many countries, we chose to prioritize hiPSC lines to enable a reference line to be globally shared. Although we are firm believers in the value of generating a series of reference lines from both genetically male and female donors of diverse genetic ancestries, we initially prioritized male lines due to the possibility that random X-chromosome inactivation may contribute to variance in gene expression (Mekhoubad et al. 2012; Bar et al. 2019) and because several genetic disorders that we plan to model are X-linked and thus more frequently expressed in males. We therefore searched public repositories and curated a series of iPSC lines, many of which have already been whole genome sequenced. We then focused on a subset of lines with broad consents for data sharing and further dissemination of the line and its derivatives, and identified KOLF2_C1, KUCG3, LNGPI1, MS19-ES-H, NCRM1, NCRM5, NN0003932, NN0004297, and PGP1 (Table S1A).

After obtaining these lines, we found that the vial of MS19-ES-H we obtained was prone to spontaneous differentiation, and therefore excluded it from further study. We then single-cell cloned each of the remaining parental cell lines (Table S1A) to reduce heterogeneity from genetic and epigenetic drift in culture. We refer to these derivatives as “sub-lines” that retain the name of the parental cell line. Simultaneously, we used CRISPR/Cas9 editing (see Methods) to correct a mutation present in one copy of *ARID2* in the KOLF2-C1 line(Hildebrandt et al. 2019) and named our sub-line KOLF2.1J to indicate its derivation at Jackson Laboratories and distinguish it from a similar sub-line derived in parallel at the Wellcome Sanger Institute (KOLF2.1S; Andrew Bassett, personal communication). We selected one to four sub-lines per parental cell line for further expansion based on their typical stem cell morphology under phase contrast microscopy and normal karyotype from the analysis of >20 Giemsa-band metaphase chromosome spreads. Almost all tested sub-lines were euploid (46; XY; Table S1B), but some sub-lines harbored aberrant cells. For example, 2 of the 20 analyzed spreads from a KUCG3 sub-line showed a gain of chromosome 12. We therefore selected a single clonal sub-line from each parental line for further expansion into 192 replicate stock vials to ensure that they could be distributed and used as a similar passage to the characterizations described in this study (Table S1C). To extend the standard karyotypic analysis, we analyzed 181 to 200 high-quality metaphase spreads for selected sub-lines at chromosomes 1, 2 and 3 by directional genomic hybridization (Robinson et al. 2019). This data established that selected sub-lines (except for KUCG3) were karyotypically normal (Table S1D), as later further confirmed by whole genome sequencing (WGS) and by the analysis of gene-edited cell clones.

### Morphology and proliferation rates

Each sub-line had the morphology expected for hiPSCs, including a high nuclear to cytoplasmic ratio, prominent nucleoli, growth in colonies with well-defined borders, and an absence of differentiated cells (Figure 1A). To compare their survival and growth rates, we dissociated each sub-line to a single-cell suspension, imaged cultures at 24 and 48 hours after plating to calculate their confluence, and then dissociated and counted cells 48 hours after plating. There was a significant difference between the sub-lines in their total cell numbers after 48 hours (one-way ANOVA, F_7, 24_= 185.1, *p*<0.0001; Figure 1B). Additionally, there was a significant main effect of time (repeated measures ANOVA; F_2, 80_ = 1836, *p*<0.0001) and cell sub-line (F_7, 80_ = 97.85, *p*<0.0001), on confluency as well as a time x cell sub-line interaction (F_14,80_=45.76, *p*<0.0001; Figure 1C, Table 1E). These results show that all sub-lines had similar morphology but varied in their survival and proliferation rates.

**Figure 1.**
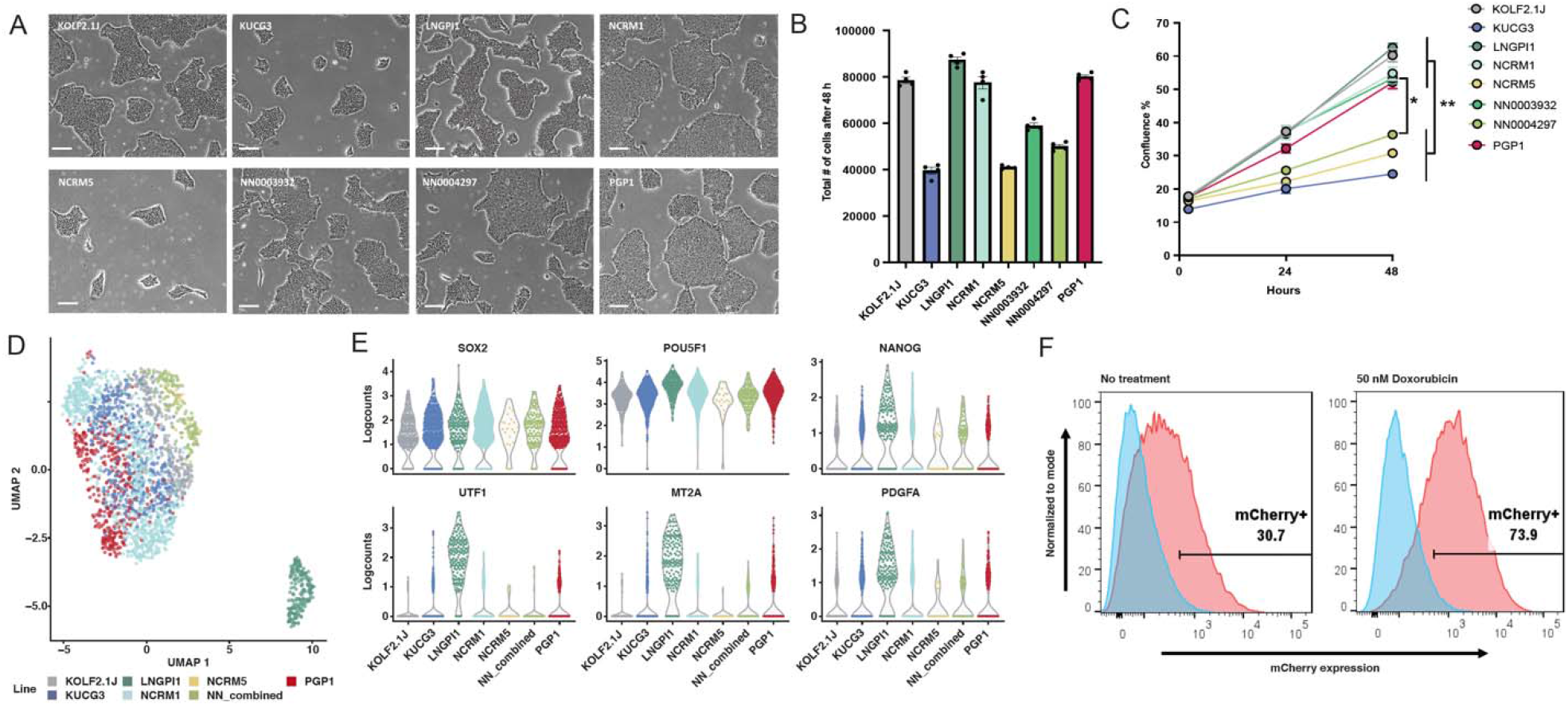
Growth, gene expression, and P53 pathway integrity of candidate cell sub-lines. (A) Representative phase contrast photomicrographs of colony morphology of the eight iPSC sub-lines on day 3 after replating. Scale bars indicate 100 µm. (B) Mean and SEM (n=4) of the total number of cells 48h after plating 30,000 cells/well. (C) Mean and SEM (n=6) of percent confluence at 0, 24 and 48 h after plating 30,000 cells/well. (D) UMAP projection of 2,270 iPSC cells color-coded by cell sub-line (E) Beeswarm plots showing expression of selected genes associated with pluripotency (top row) or poor neuronal differentiation potential (bottom row). (F) A mCherry PiggyBac reporter assay confirms the baseline activity (left plot) of the p53 pathway in KOLF2.1J (red) relative to *TP53* knockout cells (blue), which is further inducible in response to the DNA damaging agent doxorubicin (right plot).

### Pluripotency

The eight selected sub-lines were immunostained with antibodies against pluripotency markers TRA-1-60 and NANOG, and the percentage of immunopositive cells was quantified by flow cytometry analysis. We found that all analyzed sub-lines are >90% positive for both markers, except PGP1, which was 84.2% positive for NANOG (Figure S1A).

Next, we performed single-cell RNA sequencing (scRNA-seq) on all eight iPSC sub-lines (Figure 1D). We pooled them together for joint library preparation and sequencing to minimize technical sources of variation, and then used genetic diversity to assign each cell to a sub-line (Huang, McCarthy, and Stegle 2019). Since sub-lines NN0003932 and NN0004297 were derived from the same donor, these are represented in our dataset as NN_combined (Figure 1D, S1B). UMAP projection of the data from 2,270 single cells revealed two distinct groups of cells, the largest corresponding to six genetically distinct sub-lines and one small outlier group primarily from one sub-line, LNGPI1 (Figure 1D). Louvain clustering identified 5 clusters within the larger group that arise largely from cell cycle states of the proliferative cells (Figure S1C-D). Comparison of pluripotency markers across the 7 genetically distinct sub-lines showed consistent expression of core pluripotency-associated genes including *SOX2, POU5F1* (*OCT4*) and *NANOG* (Figure 1E, S1E) but LNGPI1 expressed higher levels of *UTF1* and other genes associated with inefficient neuronal differentiation (Jerber et al. 2021). Together, these findings show that all analyzed iPSC sub-lines had gene expression patterns consistent with pluripotency, and 6 of the 7 genetically distinct sub-lines showed similar transcriptional profiles.

### Integrity of the p53 response to DNA damage

Established iPSC lines can acquire genetic changes that impart a growth advantage in culture (Halliwell, Barbaric, and Andrews 2020). Of these, mutations in the *TP53* gene are recurrent(Merkle et al. 2017; Avior et al. 2021) and loss of a functional p53 pathway may be selected for during CRISPR editing and clonal expansion (Ihry et al. 2018). To measure p53 pathway function in our hiPSC sub-lines, we transfected cells with a reporter plasmid that contains 13 copies of a p53 DNA binding site linked to mCherry, and quantified mCherry expression by flow cytometry in response to treatment with vehicle or the DNA-damaging agent doxorubicin. This analysis showed doxorubicin activated reporter expression in all eight selected sub-lines compared to *TP53* knockout cells (Figure S2). To confirm these results, we developed a more stable version of the P53 reporter based on an integrating piggyBac transposon and observed similar baseline and damage-induced activity of the P53 pathway (Fig. 1F). These results confirm integrity of the p53 pathway in all lines, with particularly robust responses in KOLF2.1J, LNGPI1, NCRM1, and PGP1.

### Genomic characterization by whole genome sequencing (WGS)

To understand the genetic background of each sub-line, we sequenced their genomes at >30x coverage. The distribution of insertion-deletion (indel), loss-of-function (LOF), and missense single-nucleotide variants (SNVs) was similar across all lines (Figure 2A) and similar to human populations in the gnomAD database (Karczewski et al. 2020). When we restricted our analysis to variants with a gnomAD allele frequency of <0.001 and CADD phred score of >30 (Huang, McCarthy, and Stegle 2019), we observed a modest number of such potentially deleterious variants per cell line (Figure 2B, Table S2A). As the goal of iNDI is to model neurodegenerative diseases, we next examined WGS data for known pathogenic mutations in ADRD-associated genes. We identified heterozygous LOF variants in *DRD4* (rs587776842 in NN0003932 and NN0004297) and *MPDZ* (rs376078512 in KUCG3) but these were not the pathogenic homozygous state (Nöthen et al. 1994; Shaheen et al. 2017). Next, we tested for the presence of genetic variants that have a strong to moderate association with ADRD. Specifically, we screened for variants in the AD risk gene *APOE* (rs429358 and rs7412), the frontotemporal dementia-associated variant rs3173615 in *TMEM106B*, and the *MAPT* haplotype rs180054 associated with risk of Parkinson’s disease (PD). We found that NCRM1 and NCRM5 carry the AD risk allele *APOE* E4 and identified variants in other genes known to be risk factors for ADRD (Table S3). Finally, we calculated polygenic risk scores (PRS) for all iPSC lines based on their cumulative burden of common genetic variants associated with AD or PD (Nalls et al. 2019; Kunkle et al. 2019) and found that their PRS falls within the expected range of the population (Figure 2C). Together, these findings show that the candidate sub-lines have relatively neutral genetic risk for ADRD.

**Figure 2.**
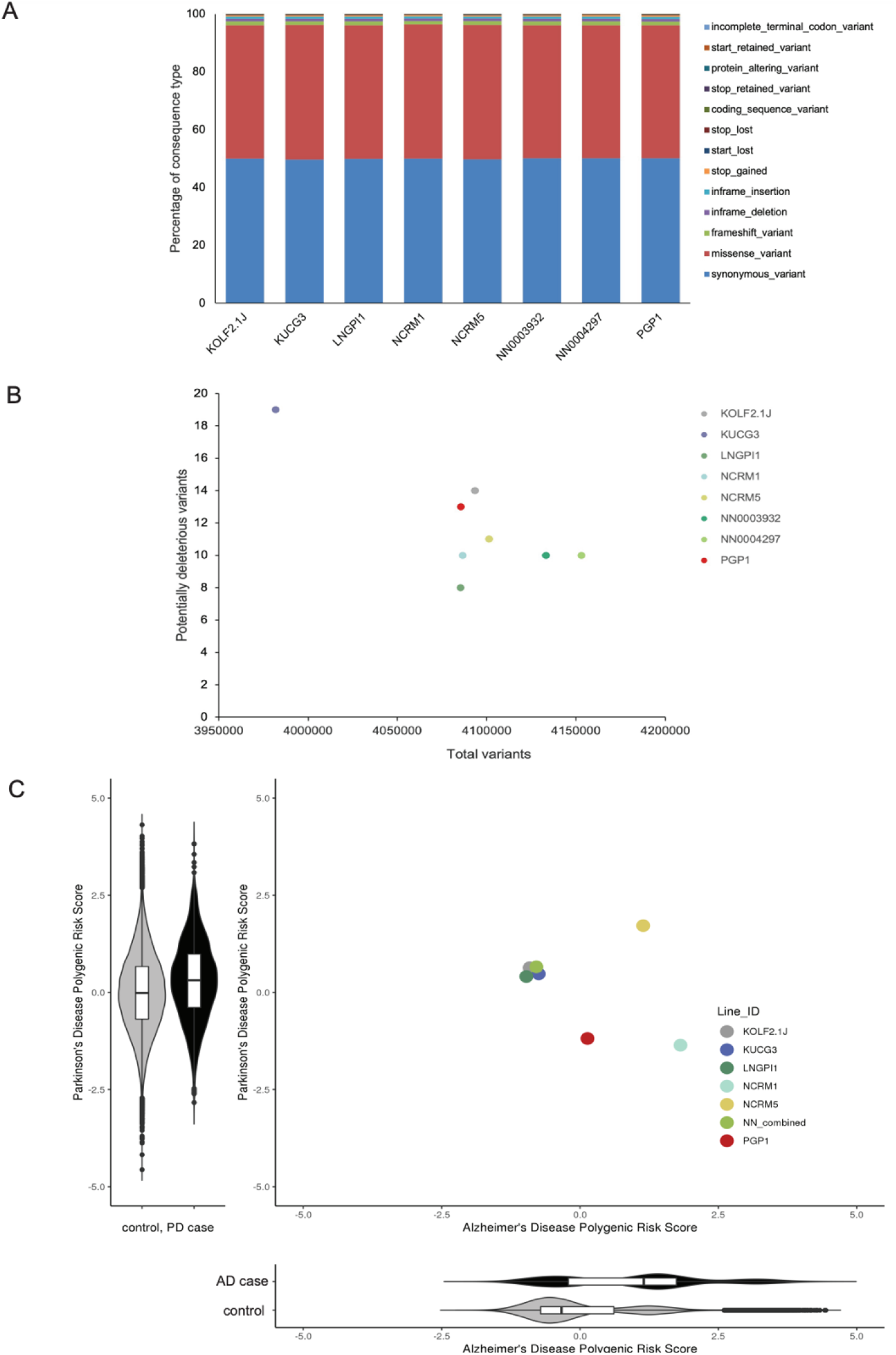
Genetic analyses of eight candidate iPSC sub-lines. (A) The percentage of genetic variant types present in the eight candidate iPSC sub-lines, grouped by their bioinformatically predicted consequences on coding sequences. (B) The number of rare (gnomAD AF<0.001) predicted deleterious (CADD phred >30) variants identified in the 8 iPSC sub-lines versus the total number of identified variants. (C) Polygenic risk scores (PRS) for Alzheimer’s disease (AD) and Parkinson’s disease (PD) are shown alongside the population-centered Z score distribution for PD PRS (y-axis) in 2995 PD cases (black) and 96,215 controls (grey) from the UK Biobank, and a similar PRS distribution for AD polygenic risk score (x-axis) in 2337 AD cases (grey) and the same controls.

### CRISPR-based gene editing potential

As many groups will want to edit the genomes of the selected hiPSC sub-line, we next characterized the efficiency with which a SNV could be introduced. Using improved conditions for homology-directed repair (Skarnes, Pellegrino, and McDonough 2019; Skarnes et al. 2021), we introduced a G to C SNV in exon 1 of the *TIMP3* gene located at Chr22q12.3 (Apte, Mattei, and Olsen 1994), resulting in an S38C missense mutation. Twenty-four edited clones from each sub-line were genotyped by Sanger sequencing of PCR amplicons spanning the targeted site of the *TIMP3* locus to quantify frequency of six possible genotypes (WT/WT, WT/SNV, WT/indel, SNV/SNV, SNV/indel, and indel/indel; Figure 3A). The overall editing efficiency of homozygous (SNV/SNV) was over 40% (Figure 3B). We also found that ratio of SNV/WT and SNV/SNV edits generated by homology directed repair (HDR) to indels generated from non-homologous end joining (NHEJ) varied across the sub-lines, with a higher frequency of WT/indel and SNV/indel clones observed in NN0003932 and NCRM1, respectively.

**Figure 3.**
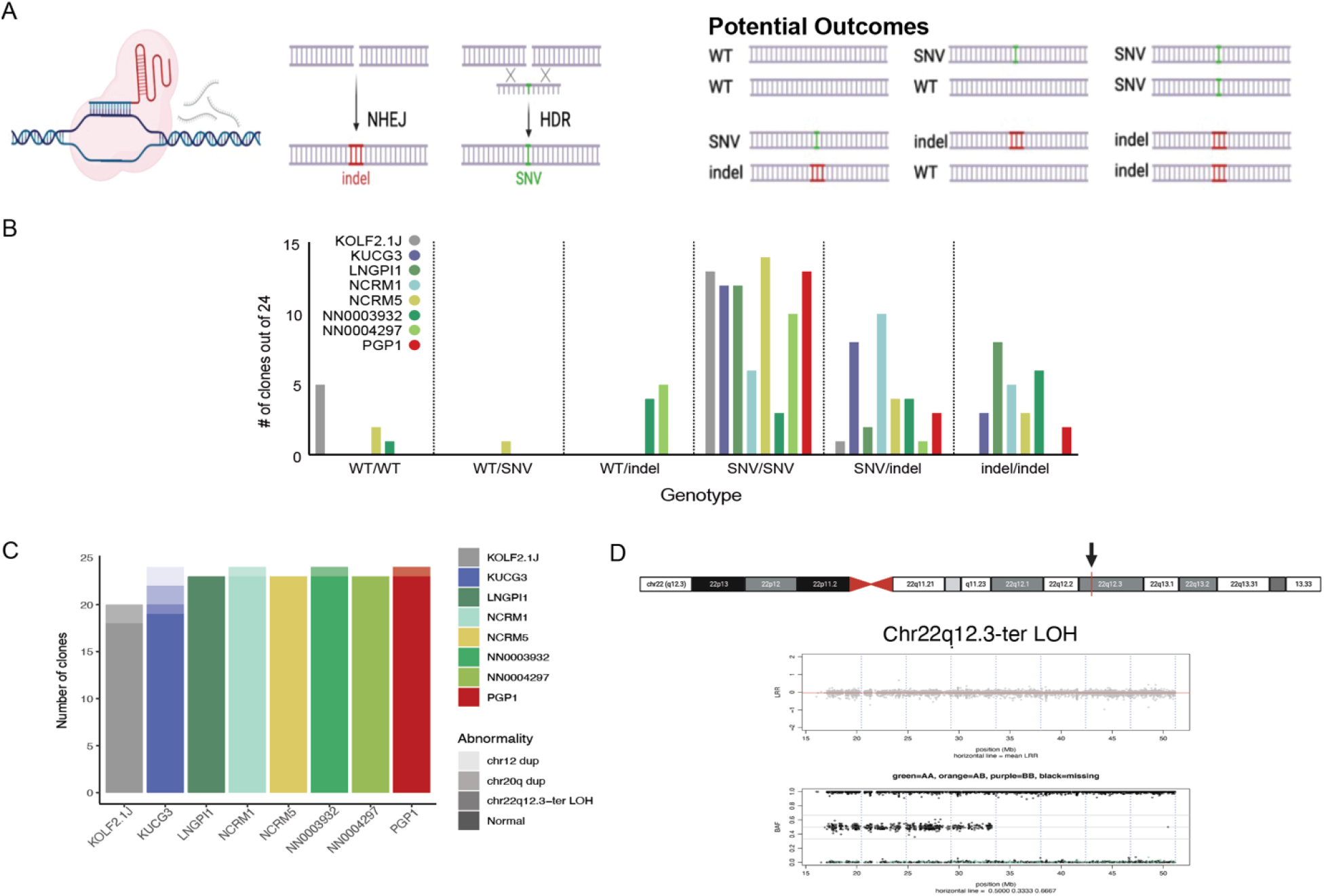
Comparative gene editing efficiency. (A) Schematic of the gene editing experiment, showing how a CRISPR/Cas9-induced double strand break can lead to the formation of insertion/deletion mutations (indels) via the non-homologous end joining (NHEJ) pathway or be repaired with a single-stranded oligodeoxynucleotide via the homology-directed repair (HDR) pathway to introduce a single nucleotide variant (SNV) of interest and resulting in 6 potential alleles. Figure created with BioRender.com (B) The number of clones out of 24 expressing each possible genotype at the targeted SNV for each analyzed sub-line. (C) Number of genomic abnormalities observed among 20-24 analyzed clones across the eight analyzed sub-lines. (D) Ideogram of chromosome 22 with the *TIMP3* gene in 22q12.3 indicated by a red bar and black arrow, using the UCSC Genome Browser. NeuroArray genotyping revealed Chr22 CN-LOH from chr22q12.3-ter in clones derived from the KOLF2.1J sub-line. Upper plots show Log R ratio (LRR) where the mean LRR (red line) is 0 for the normal clone and >1 for the abnormal clones, and middle panels show the B allele frequency (BAF) for bi-allelic probes along the arrays with evidence of duplicated alleles across the chromosome.

As CRISPR/Cas9-induced DNA double strand breaks can lead to undesired editing (Weisheit et al. 2020; Merkle, Neuhausser, et al. 2015) and because chromosomal abnormalities that confer growth advantage can be selected for in culture, we evaluated genomic fidelity of gene-edited clones using the NeuroChip DNA microarray (Blauwendraat et al. 2017). Our WGS data confirmed that parental sub-lines were clearly genetically distinct as determined by pi-hat(Purcell et al. 2007), except for clones NN0003932 and NN004297 derived from the same donor (Figure S3A). We also found very high concordance across the genome (pi-hat >0.986) when comparing gene-edited clones with their parental sub-lines using DNA microarray data, suggesting that the number of SNVs acquired during CRISPR editing was lower than the theoretical limit set by the call rates for this array (>0.968; Table S3B, Figure S3B). We then examined the microarray data to evaluate the frequency of large chromosomal abnormalities in the edited clones (Table S3B). From 185 edited clones, we identified 10 clones with chromosomal abnormalities, which involved chr12 (2 clones), chr20 (2 clones) or chr22 (6 clones). Clones derived from the KUCG3 sub-line had duplications of Chr12 (Figure S3C) previously observed by karyotyping, suggesting the sub-line harbored mosaic variants at the time of gene editing. In contrast, the absence of this variant in any of the other sub-lines (181/185, 98%) support our earlier findings that sub-lines are karyotypically normal.

We also observed recurrent chr22 abnormalities in two homozygous edited clones of KOLF2.1J and one clone each of KUCG3, NCRM1, NN0003932 and PGP1. This abnormality represents copy-neutral loss of heterozygosity (CN-LOH) from the telomere to the region of chr22q12.3 (Figure 3D) containing the target of gene editing, *TIMP3*. Intrigued by this finding, we further explored the nature and frequency of genomic variants that emerge upon CRISPR/Cas9-based gene editing by targeting additional genetic loci, as described in detail elsewhere (Skarnes et al. 2021). We found that the majority (98%, 430/440 total) of clones tested lacked detectable acquired structural variants, in marked contrast to earlier reports that CN-LOH events can be common upon gene editing(Weisheit et al. 2020). Together, these results suggest that except for KUCG3, tested iPSC sub-lines do not contain mosaic populations of large genetic variants or readily acquire them upon gene editing.

### Differentiation Potential

Since the generation of disease-relevant cell types is an important application of hiPSCs, we tested the ability of sub-lines to support cell differentiation, with an emphasis on the neural lineage. We tested four established protocols, two of which use small molecules and two of which were based on the overexpression of transcription factors (Figure 4A-B). Specifically, we directed the differentiation of iPSCs into either cortical or hypothalamic neurons and expressed either the transcription factor NGN2 to induce the formation of excitatory forebrain neurons (iNeuron, or hNGN2), or the transcription factors NGN2, ISL1, and LHX3 to induce the formation of lower motor neurons (iLowerMotorneurons, or hNIL). To assess differentiation efficiency, we profiled the differentiated cells using single-cell RNA-sequencing.

**Figure 4.**
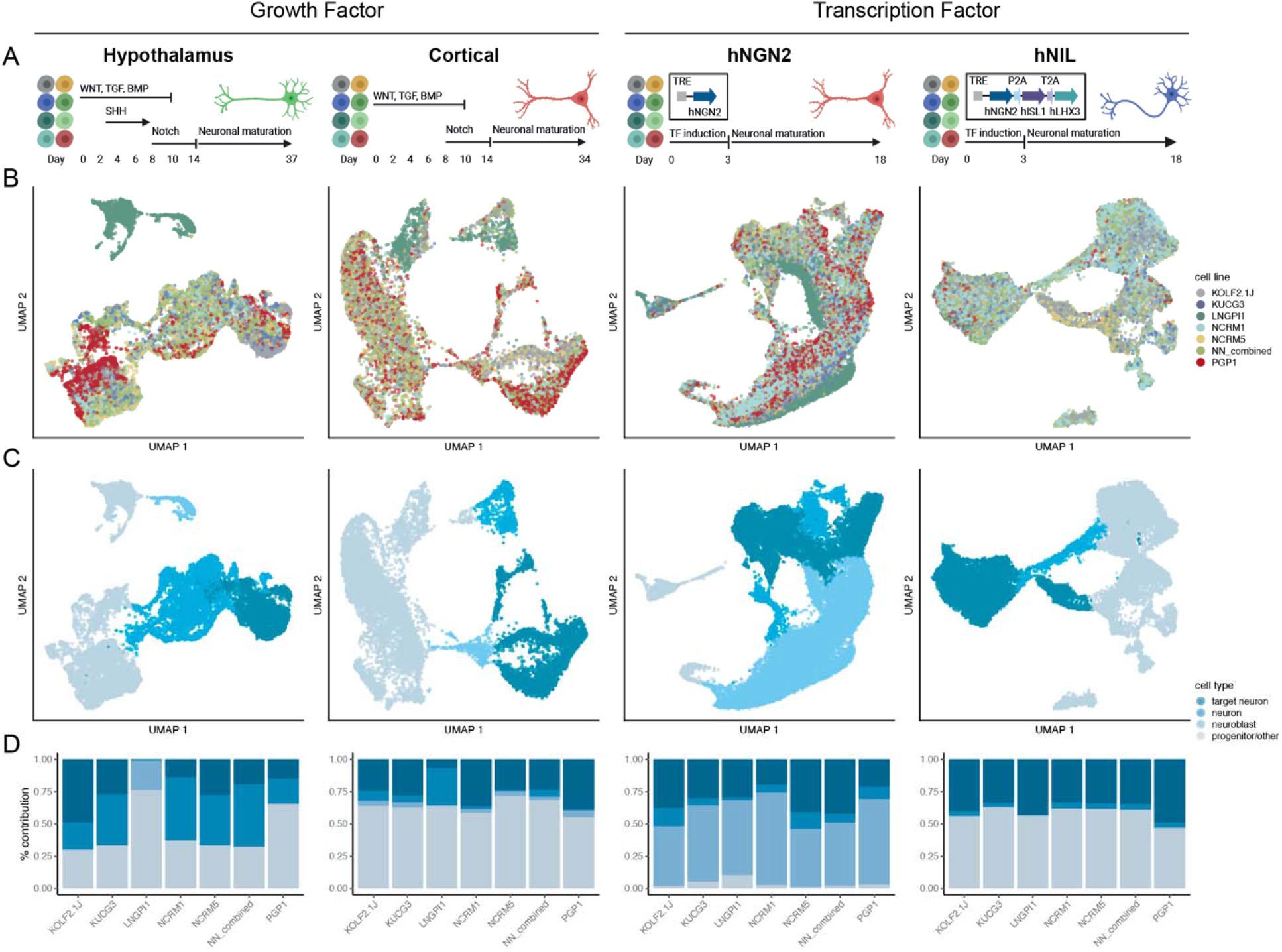
Differentiation potential of candidate cell sub-lines. (A) Schematic of experimental design for four differentiation protocols evaluated in this study: hypothalamus and cortical differentiation (growth factor-based protocols), and hNGN2 and hNIL differentiation (transcription factor-based protocols). Figure created with BioRender.com (B and C) UMAP plot for each differentiation colored by cell sub-line (B) and cell type (C). Cell type classification is derived from grouping each cluster into target neuron, neuron, neuroblast and progenitor/other categories based on the cluster annotation (Figure S4). (D) Bar plot showing proportion of cells assigned to each cell type per cell sub-line.

After quality control, demultiplexing and doublet removal, we obtained 12,818 cells from the hypothalamic protocol, 9,656 cells from the cortical protocol, 27,708 cells from the iNeuron protocol, and 18,008 cells from the iLowerMotorneurons protocol. As described above, cell sub-line identity was assigned to each cell using genotype information from the scRNA-seq reads. Louvain clustering revealed 17 clusters for hypothalamic differentiation and 13 clusters for the other 3 differentiations, which were then annotated using literature-curated genes indicative of cell identity (Figure S4). We then grouped clusters into four categories based on their expression of indicative genes: target neuron, neurons, immature neurons, or progenitor/other (Figure 4C, S4), and defined differentiation efficiency as the percentage of cells from each cell sub-line that gave rise to the target cell type (Figure 4D, Figure S5).

We found that across all four differentiation protocols, cell sub-lines consistently generated the target cell type but varied in their differentiation efficiency (Figure S6E-H). We observed that the KOLF2.1J, NCRM5, and NN_combined sub-lines consistently generated a substantial fraction of target cell types, whereas LNGPI1 did not perform well in directed differentiation protocols, as suggested by its gene expression profile in the pluripotent state (Figure 1D). Overall, the variable performance of their differentiation efficiency provides another phenotypic readout to guide the rational selection of a cell line for future development.

### Selection of KOLF2.1J as a reference cell line

Having deeply characterized the genetic and phenotypic properties of the eight lead sub-lines, we next asked if any of them showed favorable properties across all the measures we tested (Table S4). We removed LNGPI1 from consideration due to its unusual gene expression in the pluripotent state and poor differentiation properties. We also eliminated PGP1 due to possible residual expression of reprogramming factors suggested by GFP expression during FACS analysis, though we alert readers to the fact that other integration-free versions of this cell line exist (Lee et al. 2009). Though all sub-lines were amenable to CRISPR/Cas9-mediated gene editing of individual DNA bases, NN0003932 and NCRM1 showed relatively low gene editing efficiencies at the tested locus. Cell sub-lines KUCG3, NCRM5, and NN0004297 were amenable to gene editing and differentiated well but appeared to have slow growth kinetics relative to the other sub-lines, including KOLF2.1J. Consequently, since KOLF2.1J performed well across all tested assays, we selected it as a candidate reference iPSC sub-line.

### KOLF2.1J lacks genetic variants likely to cause neurological disease

We reasoned that any reference iPSC selected for large-scale studies should be extensively tested for the presence of genetic variants that might hinder the interpretation of molecular and cellular phenotypes in its differentiated progeny. First, we tested whether the process of editing *ARID2* in KOLF2.1J might have introduced unwanted mutations or culture adaptations. We therefore submitted both KOLF2.1J and the parental KOLF2-C1 line for 10x Genomics linked-read sequencing to generate phased genotyping data to complement our earlier 150 bp WGS (Figure 5A). We then called high confidence and high quality SNV and indels and tested for variants that were rare (gnomAD allele frequency < 0.001) or deleterious (CADD phred score >30). Only 25 coding variants were observed in KOLF2.1J but not KOLF2-C1, and none of them were predicted to be rare or deleterious (Table S2B; Figure 5C). These findings suggest that the process of editing *ARID2* did not select for concerning variants, consistent with the remarkable genetic stability observed upon gene editing (Figures 3 and S3).

**Figure 5.**
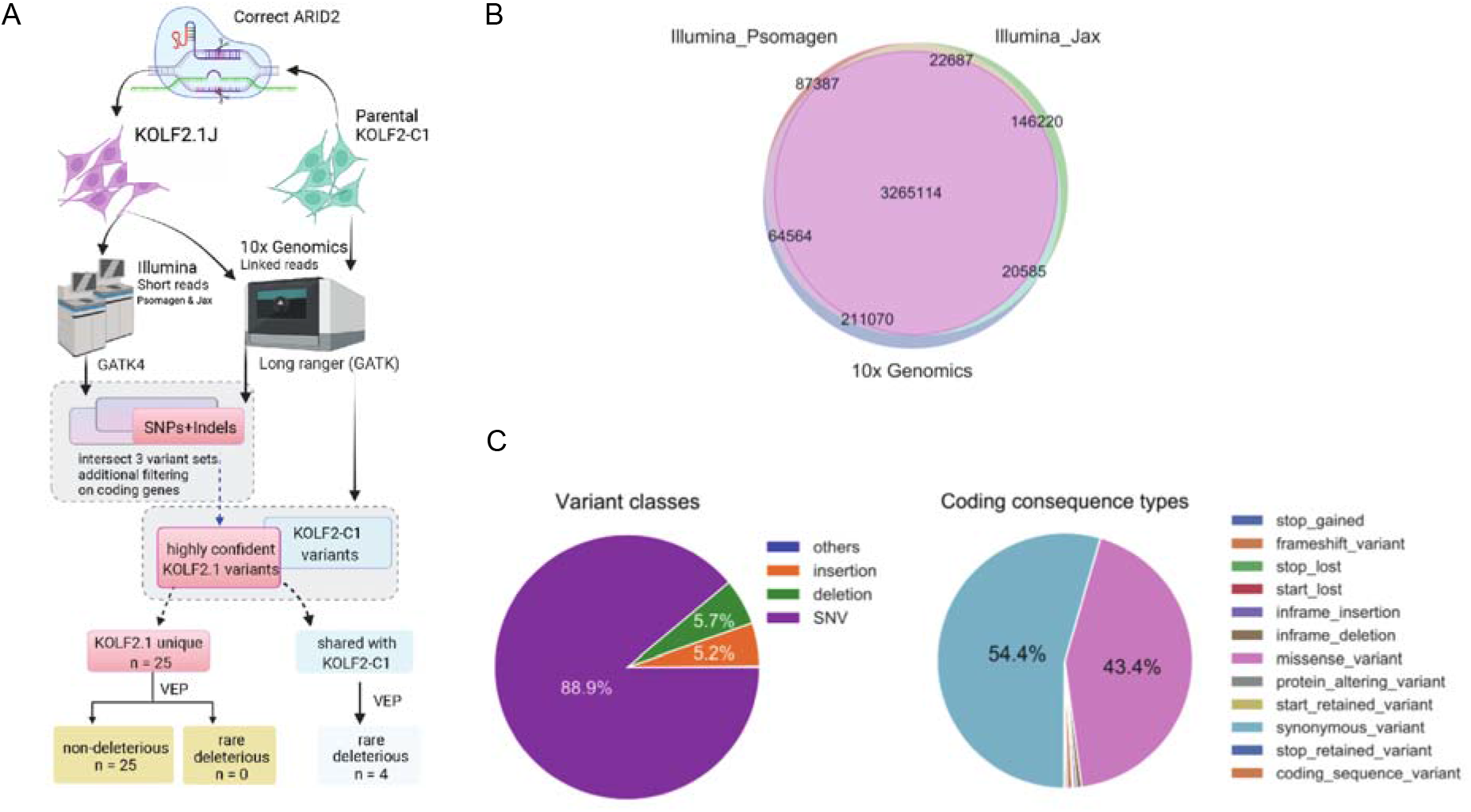
Whole genome sequencing of KOLF2.1J and the KOLF2-C1 parental line. (A) Flowchart of the discovery, filtering, annotations, and comparisons of SNP/indel variants in the *ARID2*-corrected KOLF2.1J and its parental line KOLF2-C1. Schematic created with Biorender.com (B) The consensus variant calls for KOLF2.1J from two Illumina short sequencing services (Psomagen and Jax) and one linked-read sequencing platform (10x Genomics). (C) The genetic compositions of the variant classes (left) and their effect on coding genes (right) for the 3.28 million high-confidence SNPs and indels in KOLF2.1J.

Next, we asked if KOLF2.1J might harbor other variants of concern that had been inherited from the KOLF2-C1 or the fibroblasts from which it was derived. Analysis of ∼30x WGS data showed that the overall burden of potentially deleterious SNVs in KOLF2.1J was similar to that of other iPSC lines (Figure 2B). We reasoned that combining two distinct short-read WGS datasets with the linked-read WGS dataset would enable us to filter out sequencing errors and increase our power to identify rare and potentially deleterious genetic variants. Indeed, this approach revealed 40 such variants in KOLF2.1J (Figure 5B, Table S2D). To gain insight into the origin of these variants, we manually reviewed sequencing data from the KOLF fibroblasts from which the KOLF-C1 parental line was derived and found that the vast majority of these variants (37/40, 93%) were already present in the donor fibroblasts and many of them were likely inherited form the germline. Other variants were only seen in a subset of sequencing reads, suggesting they arose as somatic mutations in a subset of the fibroblasts rather than during iPSC reprogramming, culture, or sub-cloning (Rouhani et al. 2022).

To predict which of these variants might impact cellular phenotypes of KOLF2.1J or its differentiated progeny, we prioritised genes predicted to be loss-intolerant (score < 0.03) by LoFtool (Fadista et al. 2017) or listed as haploinsufficient in ClinGen (Rehm et al. 2015). We observed variants in three such genes, including the previously described splice site disruption in *COL3A1(Hildebrandt et al. 2019)*, a gene that is associated with the vascular disease Ehlers-Danlos Syndrome (OMIM 130050). Given the role of this gene in extracellular matrix (ECM) production, we speculate that the variant will not affect most neural cell types but urge caution if using KOLF2.1J to study cell lineages or co-culture systems that interact with the ECM. We also observed a truncating variant in the homeobox gene *SHOX* that is associated with short stature (OMIM 127300) and might alter cellular phenotypes in iPSC-derived skeletal cells, but likely not cells of the neural lineage. Finally, we observed a variant in the caspase-interacting gene *DEDD* that is not associated with human disease in OMIM. Together with G-band karyotyping and dGH data showing the absence of large structural variants, these findings suggest that the genome of KOLF2.1J does not harbor genetic variants that would substantially compromise the use of this sub-line for modeling neurological disease.

### Distribution and community-based characterization of KOLF2.1J

To rigorously benchmark the performance of KOLF2.1J relative to other lines, we distributed it to groups who returned information on its growth and differentiation potential using independent approaches established in their laboratories. As of August 2022, 115 research groups successfully cultured and differentiated KOLF2.1J into numerous cell types, including three-germ layer cells (Figure 6A), cortical glutamatergic neurons (Figures 6B-C; S6A-F), cortical forebrain neurons (Figure 6D-F), skeletal myocytes (Figure S6G), motor neurons (Figures 6G; S6H-I), astrocytes (Figure 6H) microglia and macrophages (Figures 6I, S6J-K), and dopaminergic progenitors, neurons, and organoids (Figures 6J, S6L-M), using established differentiation protocols (Table S6). To rigorously test its performance, nine other research groups carried out 10 distinct differentiation protocols or assays to compare KOLF2.1J head-to-head with a total of 12 other iPSC lines.

**Figure 6.**
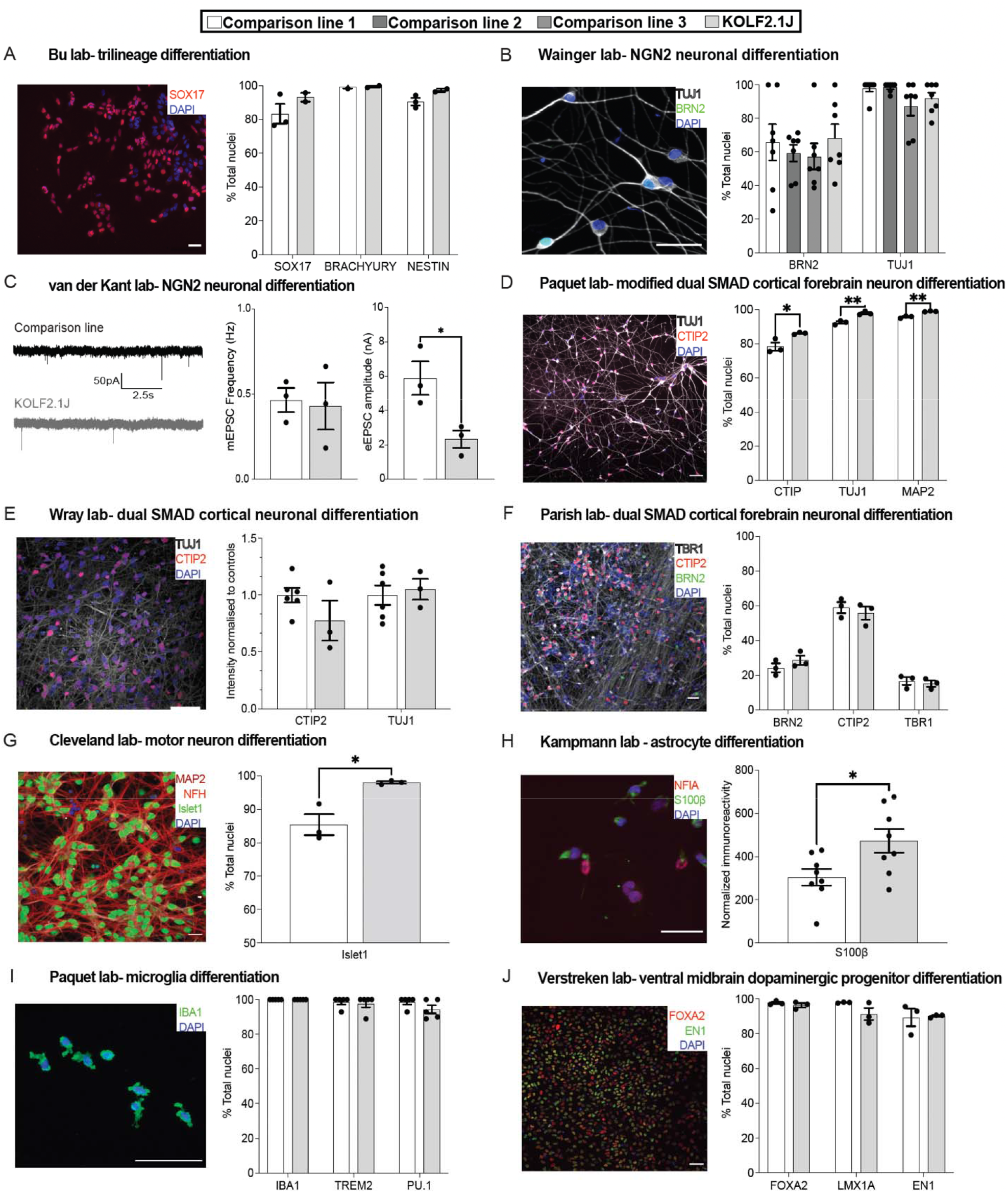
Performance of KOLF2.1J and established hPSC lines in head-to-head comparisons. In all figure panels, the performance of KOLF2.1J is shown relative to a comparison line, with the providing laboratory and differentiation method indicated. Photomicrographs of immunostaining (or recording traces) are shown to the left of each panel, where scale bars represent 50 mm and nuclear markers are in blue. Quantification of phenotypic outcomes are given on the right, where N=3 differentiations were analyzed for each cell line and unpaired t-tests were used to calculate significance unless otherwise stated. *\*p*<0.05, *\*\*p*<0.01. (A) Multi-lineage differentiation to endoderm (SOX17), mesoderm (brachyury), or ectoderm (nestin) reveals no significant differences between KOLF2.1J and APOE3 ALSTEM. (B) NGN2-induced neuronal differentiation quantified by the expression of the neuronal marker TUJ1 (grey) and the cortical layer 2/3 marker BRN2 (green) show similar expression across KOLF2.1J and comparison lines ND50003, ND50004, and 11a. n=7 wells/line, N=1 differentiation. Kruskal-Wallis test for each marker. (C) Patch clamp analysis of NGN2-induced neurons derived from BioniC13 and KOLF2.1J lines show spontaneous miniature excitatory postsynaptic currents (mEPSCs, left) in similar frequencies between the lines (center), though KOLF2.1J-derived cells appeared to have a lower excitatory EPSC amplitude (right). (D) A greater proportion of KOLF2.1J-derived cortical neurons generated by directed differentiation expressed the deep-layer cortical marker CTIP2 and TUJ1 than the ThermoFisher A18944 comparison line (D). (E) In another group, directed cortical neuron differentiation yielded similar expression of Tuj1 (grey) CTIP2 (red) as the comparison lines RBi001 and Ctrl1. Comparison lines averaged together, N=3-6 differentiations/line. (F) KOLF2.1J and H9 produced neurons expressing BRN2 (green), CTIP2 (red), and TBR1 (grey) at similar efficiencies following directed differentiation. (G) Greater efficiency of Islet1 expression was observed following differentiation of KOLF2.1J into motor neurons, compared to the CVB cell line. (H) KOLF2.1J and WTC11 cell lines were differentiated into astrocytes and immunostained for S100β (green) and NFIA (red, left), revealing a higher intensity of S100β immunoreactivity in KOLF2.1J. N=8 differentiations. (I) Microglial differentiation of KOLF2.1J and Thermo Fisher A18944 revealed no significant differences in the expression of the IBA1, TREM2, or PU.1 marker genes. Mann-Whitney test for each marker. (J) Differentiation of KOLF2.1J and SFC065 towards ventral midbrain dopaminergic neurons showed similar percentages of cells expressing the marker genes EN1 (green) and FOXA2 (red). Mann-Whitney test for EN1 expression.

Specifically, we did not observe significant differences in the potential of KOLF2.1J and a comparison line to differentiate into all three germ layers as determined by quantifying the percentage of cells immunopositive for the endoderm marker SOX17 (unpaired t-test; t_3_=1.25, *p*=0.30), the mesoderm marker brachyury (unpaired t-test; t_2_=0.13, *p*=0.91), or the ectoderm marker Nestin (unpaired t-test; t_3_=2.39, *p*=0.10; Figure 6A). Upon differentiation to glutamatergic cortical neurons through over-expression of NGN2 (Figure 6B), we did not observe significant differences between KOLF2.1J and three comparison lines in the expression of cortical markers BRN2 (Shapiro-Wilk normality test, *p*<0.05; Kruskal-Wallis test, H_3_=1.24, *p*=0.74); or TUJ1 (Shapiro-Wilk normality test, *p*<0.05; Kruskal-Wallis test, H_3_=6.48, *p*=0.09). To test the functionality of these NGN2-induced cells by patch-clamp electrophysiology, we grew them on glial islands to encourage them to make synaptic connections with themselves (Fenske et al. 2019; Rhee et al. 2019; Meijer et al. 2019) and recorded spontaneous miniature excitatory postsynaptic currents (mEPSCs) in the absence of stimulation (Figure 6C), We did not observe significant differences in their mEPSC frequency (unpaired t-test, t_4_=0.23, *p*=0.83) or amplitude (unpaired t-test, t_4_=0.40, *p*=0.71; data not shown). After inducing action potentials to record evoked excitatory postsynaptic currents (eEPSCs), we found that the KOLF2.1J-derived neurons had lower absolute amplitudes relative to the comparison line (unpaired t-test, t_4_=3.24, *p*<0.05) although there was no difference in the paired-pulse ratio (unpaired t-test, t_4_= 0.85, *p*=0.44, Figure S6E), suggesting that these differences cannot be attributed to a change in synaptic release probability.

Next, we generated cortical forebrain neurons using a modified dual SMAD protocol and found that KOLF2.1J produced a slightly greater percentage of cells expressing the neuronal markers TUJ1 (unpaired t-test, t_4_=6.08, *p*<0.01) and MAP2 (unpaired t-test; t_4_=7.80, *p*<0.01), and the neuronal subtype marker CTIP2 (unpaired t-test; t_4_=3.25, *p*<0.05, Figure 6D). These differences were not observed in other groups a slightly different protocol and examining the intensity of TUJ1 (unpaired t-test; t_7_=0.37, *p*=0.72) and CTIP2 (unpaired t-test; t_7_=1.54, *p*=0.17, Figure 6E), or the percentage of cells expressing CTIP2 (unpaired t-test, t_4_=0.65, *p*=0.55), BRN2 (unpaired t-test, t_4_=1.21, *p*=0.29), or TBR1 expression (unpaired t-test, t_4_=0.52, *p*=0.63; Figure 6F). Upon differentiation to motor neurons, we found that KOLF2.1J generated a greater percentage of cells expressing Islet 1 relative to a comparison line (unpaired t-test, t_4_=3.99, *p*<0.05, Figure 6G), and upon astrocyte differentiation, KOLF2.1J-derived cells also had greater normalized immunoreactivity for S100[(unpaired t-test, t_14_=2.51, *p*<0.05, Figure 6H). KOLF2.1J and a comparison line were similarly efficient at producing cells expressing IBA1 (identical values, t-test was not performed), TREM2 (Shapiro-Wilk normality test, *p*<0.001; Mann-Whitney test, U=12, *p*>0.99), or PU.1 (Shapiro-Wilk normality test, *p*<0.001; Mann-Whitney test, U=6.50, *p*=0.29) following differentiation into microglia (Figure 6I), and there were no differences in total cell numbers (unpaired t-test, t_4_=1.50, *p*=0.21; data not shown). Finally, when differentiated into dopaminergic neurons, we did not observe KOLF2.1J and a comparison line generated cells positive for FOXA2 (unpaired t-test, t_4_=1.01, *p*=0.37), LMX1A (unpaired t-test, t_4_=1.91, *p*=0.13), and EN1 (Shapiro-Wilk normality test, *p*<0.05; Mann-Whitney test, U=3, *p*=0.70) at similar efficiencies Figure 6J. Together, these data show that KOLF2.1J can be robustly and reproducibly differentiated into diverse functional cell types.

Having established its robust performance, we coordinated the distribution of KOLF2.1J and its gene-edited derivatives from The Jackson Laboratories via a public-facing website (https://www.jax.org/jax-mice-and-services/ipsc). Since we generated master banks of these cells prior to characterization in this study, users can be confident that distributed cell lines will have similar genetic and phenotypic properties. In addition to the cell line being readily obtainable, KOLF2.1J can be broadly used by academic groups since it was derived from material obtained under broad consent, and the accompanying material transfer agreement (MTA) is not burdensome. Our vision is that the deep genotypic and phenotypic characterization of this cell line, its proven performance in many laboratories, and its relative ease of distribution will lead to its widespread adoption by groups seeking to work with a trusted iPSC line to further the larger aim of improving reproducibility and collaboration in the field.

## Discussion

Groups working with model organisms often found that results obtained on one genetic background did not replicate on another, and that the advantages of working on a common genetic background outweigh the inherent limitations of a particular strain. For example, despite its idiosyncrasies, the inbred C57Bl/6J sub-strain has become the mouse model of choice in laboratories around the world and is distributed by organizations that perform extensive quality control to prevent genetic drift. In comparison, human iPSCs were discovered recently({Takahashi 2007) and thousands of iPSC lines have been generated and used to model disease(Bennett et al. 2018), provide HLA-matched donors for cell transplantation (Umekage, Sato, and Takasu 2019), and probe the functional effects of common genetic variants on gene expression (Mitchell et al. 2020; Jerber et al. 2021). However, the use of iPSC lines has not yet been accompanied by the widespread adoption of a common reference line, likely due to a lack of knowledge about the lines that would enable their rational selection. While some studies have performed genetic analysis of large-scale collections of hESC (Merkle et al. 2022) and hiPSC lines (Kilpinen et al. 2017), there are few studies that combine deep genetic and phenotypic characterization to support rational cell line selection. To address this issue, we sub-cloned cell lines to ensure they represented homogenous cell populations, and then expanded and banked them prior to deep characterization to ensure their genomic and phenotypic properties upon distribution would be similar to those reported in here.

We believe that KOLF2.1J is an excellent choice to become a commonly used iPSC line for several reasons. First, the parental KOLF2-C1 line was reprogrammed using non-integrating methods under feeder-free conditions in chemically defined media and substrates, and its provenance and ethical derivation are well-documented (Kilpinen et al. 2017). Second, both the parental line (Bruntraeger et al. 2019; Skarnes et al. 2021; Skarnes, Pellegrino, and McDonough 2019) and KOLF2.1J retain genomic integrity during multiple rounds of efficient CRISPR/Cas9-based gene editing (Skarnes et al. 2021).

Third, the characterization of both the KOLF2-C1 parental line (Hildebrandt et al. 2019) and the current study indicates that KOLF2.1J is free of genetic variants predicted to perturb cellular phenotypes in cells of the neural lineage (Table S2C). Fourth, KOLF2.1J performed well across all tested assays we used. Fifth, to complement the international collaborative effort that went into the initial selection of KOLF2.1J, we used a “team science” approach to confirmed its strong performance in head-to-head competitions with iPSC lines established in nine other research groups (Figure 6). Finally, we established a pipeline to distribute KOLF2.1J and its derivatives (https://www.jax.org/jax-mice-and-services/ipsc) to lower the regulatory and logistical hurdles that can frustrate the sharing of other cell lines.

Despite its good overall performance relative to other cell lines, we make no claims that KOLF2.1J is an ideal line for all purposes, just as no given individual (or strain of model organism) is free of defects. Each individual carries multiple loss-of-function mutations (Karczewski et al. 2020) and the fibroblasts from which KOLF2-C1 was derived had also acquired somatic mutations in genes including in *ARID2*, which was then corrected to produce KOLF2.1J. While the other variants we identified are of unclear potential significance, we anticipate that some groups may wish to introduce additional gene corrections to further improve the utility of KOLF2.1J. While we hope that many groups will see the value of a common reference line, which in this case is genetically male and of European ancestry, we simultaneously advocate for the establishment of other reference lines that are genetically female and/or derived from individuals of more diverse genetic ancestry. One such line might be the WTC11 due to its East Asian genetic background, established performance in gene editing (Roberts et al. 2017) and CRISPR screens (Tian et al. 2021, 2019), and its presence of a potentially neuroprotective variant (rs1990621) in *TMEM106B(Li et al. 2020)* that could be relevant to groups studying genetic modifiers of cellular vulnerability. Indeed, we hope that the workflow described in this study can serve as a blueprint to enable other groups to identify and promote additional reference iSPC lines. To facilitate such efforts, we have made the code and sequencing data from this study freely available at https://github.com/NIH-CARD/INDI-README (doi: 10.5281/zenodo.7086734) and the Alzheimer’s Disease Workbench (https://fair.addi.ad-datainitiative.org/#/data/datasets/a_reference_induced_pluripotent_stem_cell_line_for_large_scale_collaborative_studies), respectively.

## Limitations of study

While we provided a thorough characterization of candidate iPSC lines in this study, it was not comprehensive. We examined eight hiPSC sub-lines and it is possible that other lines might be more suitable for other purposes. We did not assess structural variants smaller than the detection limit of G-band karyotyping, directional genomic hybridization, and DNA microarray analysis (approximately 5 Mbp) or larger than approximately 50 bp as assessed by short-read whole genome sequencing. For KOLF2.1J, we have not found recurrent abnormalities that support cell survival such as chr20 duplication(Assou et al. 2020), but encourage users to routinely survey lines after additional editing and passages in culture. The p53 reporter assay indicated that this pathway was robustly inducible in all tested cell sub-lines, suggesting that any advantages in growth rate are not due to acquisition of oncogenic potential, but we cannot formally exclude this possibility. Since our primary motivation for selecting a well-performing iPSC line was to generate transformative foundational data and resources for ADRD through the iNDI initiative, our differentiation analysis was biased toward the neural lineage.

## Acknowledgements

This research was supported in part by the Intramural Research Program of the NIH, National Institute on Aging (NIA), National Institutes of Health, Department of Health and Human Services; project number [ZO1 AG000535], as well as the National Institute of Neurological Disorders and Stroke. A.Y. is supported by an EMBL-EBI/Cambridge Computational Biomedical Postdoctoral Fellowship (EBPOD).V.P. is supported by a UK Regenerative Medicine Platform grant from the Medical Research Council [MR/R015724/1]. The costs of directed differentiation and single-cell sequencing were supported by the Chan Zuckerberg Initiative [191942]. F.T.M. is a New York Stem Cell Foundation - Robertson Investigator [NYSCF-R-156] and is supported by the Wellcome Trust and Royal Society [211221/Z/18/Z]. J.C.M. acknowledges core funding from the European Molecular Biology Laboratory and from Cancer Research UK [C9545/A29580]. We thank Greg Strachnan and the MRC Metabolic Diseases Unit Imaging Core Facility for assistance with imaging, and Katarzyna Kania and the staff at the Genomics Core Facility of the Cancer Research UK Cambridge Institute for assistance with single-cell library preparation and sequencing. This study utilized the high-performance computational capabilities of the Biowulf Linux cluster at the National Institutes of Health, Bethesda, MD, USA. (http://biowulf.nih.gov). This research has been conducted using the UK Biobank Resource under Application Number 33601. We gratefully acknowledge the contribution of the Scientific Services at The Jackson Laboratory for work described in this publication, including the Cellular Engineering service, the Genome Technologies service for expert assistance with whole genome sequencing and DNA and RNA preparation, the Flow Cytometry service for expertise in the p53 assay, and Scientific Instrument Services. We further thank the Cytogenetics laboratory at JAX for G-band karyotyping and analysis of several iPSC sub-lines.

L.S. is supported by the Training Program in Stem Cell Biology fellowship from the New York State Department of Health (NYSTEM-C32561GG). Funding from the Dan and Diane Riccio Fund for Neuroscience was provided to D.A.B. M.S.B. is supported by T32AG066596-01 training grant. D.W.C. is funded by R01-NS27036. M.D. and S.C. are supported by a CZI Neurodegeneration Challenge Network Collaborative Pairs Award. J.V.L. is supported by a DOC-PRO4 PhD fellowship from the University of Antwerp and an FWO-Flanders travel grant to perform the research at Gladstone Institute. E.L is funded by the National Defense Science and Engineering Graduate Fellowship. D.D. was supported by T32-AG-000255. E.L.F.H. is funded by NINDS R01 NS060698. I.V.L.R. is funded by the California Institute for Regenerative Medicine Scholars Research Training Program (EDUC4-12812) and the Ruth L. Kirschstein National Research Service Award (T32 NS115706). M.K. is funded by a Chan Zuckerberg Initiative Ben Barres Early Career Acceleration award. R.v.d.K is supported by a Chan Zuckerberg Initiative Neurodegeneration Challenge Network Pilot Grant (2020-222244(5022)). L.E. is supported by the Studienstiftung des deutschen Volkes. M.R is supported by R01GM127557 and the Tufts Springboard Award. L.H is funded by the Deutsche Forschungsgemeinschaft (DFG, German Research Foundation; project number: 471227244). D.K-V. is supported by grants from the Chan Zuckerberg Initiative (2020-221779(5022) & 2021-235147) and Alzheimer Forschung Initiative e.V. (20019p). B.S. and P.V are supported by the Chan-Zuckerberg (CZI) Neurodegeneration Challenge Network and Amici Lovanienses. C.C. is funded by the Marie Skłodowska-Curie Actions - Seal of Excellence FWO and postdoctoral fellowship. P.V. is supported by Mission Lucidity, an ERC consolidator grant, and Research Foundation Flanders (FWO). B.J.W is a New York Stem Cell Foundation - Robertson Investigator and is supported by The New York Stem Cell Foundation [NYSCF-I-R44]. A.H. is supported by NRSA postdoctoral fellowship F32NS114319. G.L. and E.A. are supported by: EU H2020-MSCA-ITN-2018 (813851), CZI Foundation Neurodegeneration Challenge Network (2018-191929), Wallenberg Scholar (KAW2018.0232), ERC Advanced (PreciseCellPD, 884608), SSF (SB16-0065), VR (2020-01426), Hjarnfonden (FO2019-0068), Cancerfonden (CAN2016/572), Karolinska Institutet (StratRegen 2018). D.P. is supported by grants from the Deutsche Forschungsgemeinschaft (DFG, German Research Foundation) under Germany’s Excellence Strategy within the framework of the Munich Cluster for Systems Neurology (EXC 2145 SyNergy – ID 390857198). S.W and R.G are supported by Alzheimer’s Research UK (ARUK-SRF2016B-2), the UCL Neurogenetic Therapies Programme, generously funded by The Sigrid Rausing Trust and the National Institute for Health Research University College London Hospitals Biomedical Research Centre. K.J.C. and K.Z is supported by grants from NIH-NINDS/NIA (R01NS117461), DoD (W81XWH-21-1-0082), Target ALS, and the Frick Foundation for ALS Research.

## Author contributions

All co-first authors (C.B.P., A.Y., E.L., J.A.M., C.B., and L.P.) contributed equally to this manuscript and all authors agree that they can both indicate their equal contribution and re-order the list of co-first authors in their own publication records.

Conceptualization: C.B.P., A.Y., E.L., J.A.M., C.B., L.P., M.A.N., S.L.C., S.W.S., R. v.d.K., D.P., G.B., B.J.W., S.W., C.L.P., D.W.C., M.K., P.V., B.S., E.L.F.H., S.C., B.R.C., K.Z., D.A.B., D.K-V., E.A., T.A., J.C.M., W.C.S., M.R.C., M.E.W., F.T.M

Methodology: C.B.P., A.Y., E.L., J.A.M., C.B., L.P., M.A.N., S.L.C., S.W.S., R. v.d.K., D.P., G.B., B.J.W., S.W., C.L.P., D.W.C., M.K., P.V., B.S., E.L.F.H., S.C., B.R.C., K.Z., D.A.B., D.K-V., E.A., T.A., J.C.M., W.C.S., M.R.C., M.E.W., F.T.M

Validation: A.Y., E.L., J.A.M., C.B., L.P., H.O., J.K., T.P., M.O., J.D., M.M., L.J.M.J., V.S.L., R.v.d.K., D.C., D.P., A-C.R., G.B., A.H., B.J.W., R.M.C.G., J.M.C., S.W. D.A-B., C.L.P., M.S.B., D.W.C., E.L., I.V.L.R., M.K., C.C.A., P.V., L.H., M.Y.C., B.S., D.D., E.L.F.H., M.C.Z., R.B., M.D., S.C., R.K., M.R., Z.S.N., M.M., J.V.L., B.R.C., K.J.C., K.E., S.F., D.A.B., L.E., M.W., D.K-V., G.L., E.A., E.C., L.S., T.A., J.C.M., W.C.S., M.R.C., M.E.W., F.T.M.

Formal Analysis: C.B.P., A.Y., E.L., J.A.M., C.B., L.P., D.S., G.P., E.C., J.B., P.B., L.K-B., C.M., C.T., S.H., W.C.S., M.R.C., M.E.W., F.T.M.

Investigation: A.Y., E.L., J.A.M., C.B., L.P., H.O. J.K., J.Z., D.S., G.P., E.C., J.B., P.B., L.K-B., C.M., C.T., S.H., M.S., F.F., M.A.N., D.V., S.B., Y.A.Q., D.M.R., K.A.M., J.P., L.R., M.P.N., S.A., L.U.R., P.K., V.P., S.L.C., S.W.S., T.P., M.O., J.D., M.M., L.J.M.J., V.S.L., R.v.d.K., D.C., D.P., A-C.R., G.B., A.H., B.J.W., R.M.C.G., J.M.C., S.W. D.A-B., C.L.P., M.S.B., D.W.C., E.L., I.V.L.R., M.K., C.C.A., P.V., L.H., M.Y.C., B.S., D.D., E.L.F.H., M.C.Z., R.B., M.D., S.C., R.K., M.R., Z.S.N., M.M., J.V.L., B.R.C., K.J.C., K.E., S.F., D.A.B., L.E., M.W., D.K-V., G.L., E.A., E.C., L.S., T.A., J.C.M., W.C.S., M.R.C., M.E.W., F.T.M.

Resources: all co-authors

Data Curation: C.B.P., A.Y., C.B., L.P., F.F., M.A.N., D.V., S.B.

Writing- Original Draft: C.B.P., A.Y., E.L., J.A.M., C.B., L.P., W.C.S., M.R.C., M.E.W., F.T.M

Writing- Reviewing & Editing: all co-authors

Supervision: J.Z., D.S., G.P., F.F., M.A.N., S.L.C., S.W.S., R. v.d.K., D.P., G.B., B.J.W., S.W., C.L.P., D.W.C., M.K., P.V., B.S., E.L.F.H., S.C., B.R.C., K.Z., D.A.B., D.K-V., E.A., T.A., J.C.M., W.C.S., M.R.C., M.E.W., F.T.M

Visualization: C.B.P., A.Y., E.L., J.A.M., C.B., L.P., H.O., J.K., J.Z., D.S., G.P., E.C., J.B., P.B., L.K-B., C.M., C.T., S.H., M.S., K.M.A., T.P., M.O., J.D., M.M., L.J.M.J., V.S.L., R.v.d.K., D.C., D.P., A-C.R., G.B., A.H., B.J.W., R.M.C.G., J.M.C., S.W. D.A-B., C.L.P., M.S.B., D.W.C., E.L., I.V.L.R., M.K., C.C.A., P.V., L.H., M.Y.C., B.S., D.D., E.L.F.H., M.C.Z., R.B., M.D., S.C., R.K., M.R., Z.S.N., M.M., J.V.L., B.R.C., K.J.C., K.E., S.F., D.A.B., L.E., M.W., D.K-V., G.L., E.A., E.C., L.S., T.A., J.C.M., W.C.S., M.R.C., M.E.W., F.T.M.

Project Administration: C.B.P., A.Y., E.L., J.A.M., C.B., L.P., H.O., J.Z., D.S., G.P., M.S., F.F., M.A.N., P.N., J.P., S.L.C., R. v.d.K., D.P., G.B., B.J.W., S.W., C.L.P., D.W.C., M.K., P.V., B.S., E.L.F.H., S.C., B.R.C., K.Z., D.A.B., D.K-V., E.A., T.A., J.C.M., W.C.S., M.R.C., M.E.W., F.T.M.

Funding Acquisition: M.A.N., S.L.C., S.W.S., R. v.d.K., D.P., G.B., B.J.W., S.W., C.L.P., D.W.C., M.K., P.V., B.S., E.L.F.H., S.C., B.R.C., K.Z., D.A.B., D.K-V., E.A., T.A., J.C.M., M.R.C., M.E.W., F.T.M

## Materials availability

Cell lines and sublines used in this study are available upon request pending approval from the original suppliers of these lines. KOLF2.1J and its derivatives are available from The Jackson Laboratories at https://www.jax.org/jax-mice-and-services/ipsc. Plasmids generated in this study have been deposited to Addgene. Code we developed as part of this study is available via Github at https://github.com/NIH-CARD/INDI-README (doi: 10.5281/zenodo.7086734). Cell culture maintenance and Piggybac-TO-NGN2 transfection protocols are available on Protocols.IO at dx.doi.org/10.17504/protocols.io.n2bvjxm2nlk5/v1 and dx.doi.org/10.17504/protocols.io.q26g744b1gwz/v1, respectively.

## Methods

### Plasmids

The plasmids PB-TO-hNGN2 (Addgene plasmid #172115), PB-TO-hNIL (NGN2, ISL1, and LHX3, Addgene plasmid #172113), EFa1-Transposase (Addgene plasmid #172116) were used for transcription factor-based differentiation experiments. For the p53 reporter assay performed on all sub-lines, the PG13-mCherry p53 reporter plasmid was generated by replacing the luciferase sequence of PG13-Luc reporter plasmid (el-Deiry et al. 1993) (Addgene plasmid #16442), which contains 13 copies of a p53-binding site followed by the polyoma promoter, with the mCherry sequence by the In-Fusion HD Cloning Plus kit (Takara, 638910). For the transposon-based p53 reporter assay performed on KOLF2.1J, the pCMV-hyPBase plasmid was obtained from Wellcome Trust Sanger Institute (Yusa et al. 2011). The PB-PG13-mCherry-EF1a-PuroR-EGFP plasmid, generated by inserting the PG13-mCherry sequence into the PB-EF1a-PuroR-EGFP plasmid (H Oguro, unpublished), contains the PiggyBac 3’-inverted terminal repeat (PB 3’ITR), 5’-HS4 chicken β-globin (cHS4) insulator, PG13-mCherry reporter, woodchuck hepatitis virus post-transcriptional regulatory element (WPRE), bovine growth hormone polyadenylation signal (BGHpA), EF1α promoter, puromycin registrant gene, P2A peptide, EGFP, BGHpA, cHS4 insulator, and PB 5’ITR, in this order.

### Sub-cloning

Eight candidate iPSC lines were put through a uniform workflow for sub-line generation, expansion, and archiving, as described in detail in the Supplementary Information.

### Proliferation rates

iPSCs were maintained in Essential 8 medium (Thermo Fisher Scientific) on Matrigel (1:100, Corning)-coated plates and passaged at 70-80% confluence with Accutase (Thermo Fisher Scientific) to a single-cell suspension. Dissociated iPSCs were plated onto a Matrigel coated 48 well plates at 30,000 cells/well in Essential 8 medium and 10 µM Rock inhibitor Y-27632 (Selleck Chem, n=6 wells per line). After 24 hr, the media was changed to Essential 8 medium. Plates were scanned in an Incucyte® S3 Live-Cell Analysis System every 24 h and confluence was analysed with Incucyte software, Basic Analysis. After 48 hr, iPSCs were dissociated with Accutase and total cell numbers were counted (n=4 wells per line).

### Flow cytometry analysis of pluripotency markers

iPSCs were dissociated into single cells using TrypLE (Thermo) for 5-10 minutes, pelleted by centrifugation at 200x*g* for 5 minutes, and then fixed in 500 µL 4% paraformaldehyde for 10-15 minutes at room temperature. After fixation, cell pellets were washed with 1mL PBS, and incubated with 50 µl of permeabilization buffer (PBS plus 2% FBS and 0.2% Tween-20) for 10 minutes. During the permeabilization, 1 µl antibody was diluted in a 5 µl permeabilization buffer and added to 96-well for each staining reaction, then 50 µl permeabilized cells were added to each well and incubated for 1 hour at 4°C with mixing occasionally. After staining, cell pellets were washed and resuspended with 200 µl per-well PBS for flow cytometry analysis. The antibodies and isotype controls used were: TRA-1-60 Monoclonal Antibody (TRA-1-60), DyLight 488 conjugate (Life Technology MA1-023-D488X); Mouse IgM Isotype Control, DyLight 488 conjugate (Life Technology MA1-194-D488); CABS352A4 Milli-Mark™ Anti-Nanog-Alexa Fluor 488 Antibody, NT (EMD Millipore FCABS352A4); Rabbit IgG isotype control, AlexaFluor 488 conjugate, (Cell Signaling 4340S).

### p53 reporter assay

iPSCs were maintained in StemFlex medium (Thermo Fisher, A3349401) on Synthemax II-SC substrate (Corning). For the plasmid-based assay, at 70% confluence, the culture medium was replaced with fresh medium supplemented with RevitaCell (Thermo Fisher) five hours before nucleofection. Cells were dissociated with pre-warmed Accutase (STEMCELL Technologies) at 37°C for 7 minutes and 4×10^5^ cells were transferred to a well of a 96-well V-bottom plate (Corning) then centrifuged at 100x*g* for 3 minutes. Cell pellets were resuspended with 20 µL of P3 Primary Cell Buffer (Lonza) containing 5 µg of the PG13-mCherry reporter plasmid and transferred to a well of a 16-well Nucleocuvette strip, followed by nucleofection with the 4D-Nucleofector Unit (Lonza) using the CA-137 pulse code. After nucleofection, 1.5×10^5^ cells (for the no treatment group) or 2.5×10^5^ cells (for the doxorubicin treatment group) were seeded in the StemFlex medium supplemented with RevitaCell on a well of a 48-well plate coated with Matrigel hESC-Qualified Matrix (Corning). One day after nucleofection, the medium was changed to StemFlex without RevitaCell. Two days after nucleofection, the medium was changed to StemFlex with or without 20 nM doxorubicin (Bio-Techne, 2252). Three days after nucleofection, cells were dissociated with Accutase and mCherry expression in the singlet cell population was analyzed using a FACSymphony A5 flow cytometer (BD Biosciences).

For the transposon-based P53 reporter assay, 2×10^5^ cells were nucleofected with the PB-PG13-mCherry-EF1a-PuroR-EGFP and pCMV-hyPBase plasmids (0.75 μg each) with the 4D-Nucleofector Unit using the condition described above, and seeded onto a well of a Synthemax-coated 12-well plate. Transposon-integrated cells were selected by adding 2 μg/mL puromycin (MilliporeSigma, P8833) to the StemFlex medium for 3 days or longer. Selected cells were transferred to a Matrigel-coated 24-well plate followed by 50 nM doxorubicin (or vehicle) treatment for 24 hours. Cells were dissociated with Accutase, and mCherry expression in the EGFP positive singlet cell population was analyzed using the BD LSR II flow cytometer (BD Biosciences).*TP53*-deficient KOLF2 cells (W Skarnes, unpublished) were used as a negative control. Non-viable cells were excluded by staining with 4’,6-diamidino-2-phenylindole (Thermo Fisher, D1306). Flow-cytometric data were analyzed using FlowJo software (BD Biosciences).

### DNA and RNA preparation

DNA and RNA extraction was performed by the JAX Genome Technologies service, quantified by TapeStation (Agilent), and assigned a DIN or RIN value. The extracted DNA and RNA from each of the 8 sub-lines was submitted to Psomagen for Illumina short read whole genome sequencing. From an additional well, high molecular weight genomic DNA extraction was performed by the JAX Genome Technologies service, quantified by TapeStation (Agilent), and assigned a DIN value. The DNA from each of the 8 sub-lines was submitted to Psomagen for 10x Genomics long read whole genome sequencing. From an additional well, genomic DNA for each sub-line was also prepared using the DNeasy Blood and Tissue kit (Qiagen) and submitted to the JAX Genome Technologies service for Illumina short read whole genome sequencing.

### Whole genome sequencing and annotation of variants

iPSC lines and a National Institute of Standards and Technology (NIST) reference (HG-002) were sequenced with 30x coverage and paired-end through the Illumina short read sequencing and the 10x Genomics linked-read sequencing by Jax and/or Psomagen, Inc. For Illumina short read data, SNVs and indels were called using the HaplotypeCaller (link) following the Genome Analysis Toolkit (GATK) best practices and executed through the Google genomics alpha pipeline. FASTQ files were processed into unmapped BAM files using the paired-fastq-to-unmapped-bam workflow on the human GRCh38 build. Initial variant calling was performed using the PairedSingleSampleWf. The joint discovery was then executed using the JointGenotypingWf. Variants were filtered using the variant quality score recalibration (VQSR) with default filtering parameters. The structural variant calling was performed using the Manta algorithm (Version 1.6.0) and then standardised using the structural variant tool kit (SVTK). For the 10x Genomics linked-read data, the SNP and indel variants and the structural variants were called using the 10x Genomics LongRanger wgs (version 2.2) pipeline. Sequencing reads were aligned to the human GRCh38 build containing decoy contigs and subjected to variant calling and phasing. The GATK’s HaplotypeCaller mode was applied to call SNPs and Indels.

Variants were also annotated using ANNOVAR (Wang, Li, and Hakonarson 2010) including the ClinVar database (version clinvar_20200316) to identify potential known pathogenic variants. Additionally, all data were screened for loss of function variants (stop, frame-shift and splicing) in INDI project genes and specific variants of interest including *APOE* haplotype (rs429358 and rs7412), *MAPT* haplotype (rs1800547) and *TMEM106B* (rs3173615) genotype. Polygenic risk scores for AD and PD were calculated using PLINK (v1.9) with the weights of recent GWAS (Kunkle et al. 2019; Nalls et al. 2019). As a reference population for the polygenic risk score we used AD (Data-Field 131037), PD (Data-Field 131023) and controls (no known neurodegenerative disease, no parent with a known neurodegenerative disease and >=60 years old at recruitment) from the UK Biobank (application ID: 33601)(Bycroft et al. 2018).

### CRISPR/Cas9 genome editing

Editing was performed on each iPSC sub-line by high-throughput engineering of a missense mutation (S38C) in exon 1 of the *TIMP3* gene, using optimized conditions for homology-directed repair (HDR) (Skarnes, Pellegrino, and McDonough 2019). Cas9 sgRNA to *TIMP3* (CCAGGAGCGCTTACCGATGT/CGG) was chemically synthesized with 2’-O-methyl and 3’-phosphorothioate end modifications (Synthego CRISPRevolution sgRNA) and resuspended in TE buffer at a concentration of 4 µg/µl. RNP was formed by combining SpCas9 nuclease (HiFi V3, IDT) with sgRNA at a molar ratio of 1:4. A 100-nt single stranded oligo donor (ssODN) containing a G to C SNV was synthesized with HDR-optimized end modifications (Alt-R™ HDR Donor Oligo, IDT) and resuspended in DPBS-/- at a concentration of 200 pmol/µl. For high-throughput introduction of Cas9 RNP and ssODN into human iPS cells, 8 wells were transfected using Amaxa nucleofection with P3 Primary Cell 4D-Nucleofector 16-well Strips (Lonza). Each well contained a single-cell suspension of 1.6 × 10^5^ cells in 20 µl of Primary Cell P3 buffer with supplement (Lonza) containing 2 µg Cas9, 1.6µg sgRNA, and 40 pmol ssODN. Nucleofection was performed using Amaxa program CA-137. Immediately following electroporation, cells were distributed to wells of a Matrigel-coated 24-well plate containing StemFlex, RevitaCell, and 30 µM final Alt-R® HDR enhancer (IDT). Cells were incubated at 32°C for 3 days before transfer to 37°C. At 24h post-nucleofection, and every other day thereafter, the media was replaced with only StemFlex. Upon reaching near confluency, cells were single-cell-plated into Synthemax-coated 10cm dishes as described above. At Day 10, 24 colonies per cell sub-line were manually picked as described (Skarnes, Pellegrino, and McDonough 2019) and incubated in Matrigel-coated 96-well plates for 4-5 days before being frozen down.

Crude cell lysates for each clone were prepared as described (Skarnes, Pellegrino, and McDonough 2019) and used to amplify a 896 bp genomic region containing the CRISPR target site. Sanger sequencing of purified PCR products was carried out by the Genome Technologies service at The Jackson Laboratory in Bar Harbor, Maine. Sequence traces were aligned and analysed using SeqManPro (https://www.dnastar.com/software/lasergene/molecular-biology/) and Synthego ICE (https://ice.synthego.com/). Two additional wells of each clone were lysed, pooled, and genomic DNA was purified using the 96-well high-throughput DNeasy Blood and Tissue kit (Qiagen). Array-based genotyping was performed on the resulting genomic DNA.

### Genotyping array genotyping of subclones

To assess the genomic fidelity of IPSC sub-lines and subclones after editing, DNA was isolated and Illumina genotyping array was performed using the NeuroChip array and standard Illumina genotyping protocols (Blauwendraat et al. 2017). In total 185 subclones were successfully genotyped including at least 20 clones per included IPSC sub-line. Genomic fidelity was assessed using two strategies; 1) comparison between genotyping array data and WGS data and 2) Assessment of genome wide B-allele frequency and Log R ratio values. For the comparison between genotyping array and WGS data, all data was merged using PLINK (v1.9) and only overlapping variants were kept. Potential genetic differences were identified using the --merge-mode 7 option in plink which reports mismatching non-missing calls between two datasets. Variants discordant in more than 33% of the genotyped clones were excluded due to high likelihood of being genotyping errors. Mismatching non-missing calls were plotted using R (v3.6.1) per chromosome and visually inspected for large clusters of discordant array genotypes and WGS. Genotyping array data was also assessed for large events based on the B-allele frequency and Log R ratio values. The B-allele frequency and Log R ratio values were downloaded from Illumina GenomeStudio and processed and plotted using the GWASTools package in R (v3.6.1) (Gogarten et al. 2012). GenCall score variant filtering thresholds of >0.4 and >0.7 were used to filter out calls likely arising from genotyping errors.

### Comparative whole genome sequence analysis of KOLF2-C1 and KOLF2.1J

To retrieve the highly confident variants for KOLF2.1J, the three variant sets originally discovered from its default variant calling pipeline was subjected to filtering to exclude those that did not pass the thresholds of PASS, QUAL >= 30, DP >= 10, QP >= 2.0, and MQ >= 40. After filtering, the variant sets were intersected to generate the 3,278,414 common variants (SNPs/Indels) of high confidence, illustrated in Figure 5B. The common variants were subjected to annotation and effect prediction using VEP. About 88.9% variants were SNVs (Figure 5). Of the coding variants, 54.4% were found to be synonymous, 43.4% were missense, and the remainder were LOF, splice site, or other types of variant (Figure 5). KOLF2.1J highly confident SNPs/Indels were compared to KOLF-C1 SNPs/Indels to exclude the possibility that it gained deleterious variants after editing (Figure 5B). The rare and deleterious coding genes that were unique to KOLF2.1J were predicted using VEP (gnomAD_NFE_AF < 0.001 and CADD_PHRED > 30). Clinical relevance and dosage sensitivity of the predicted deleterious variants was annotated through ClinGen (Rehm et al. 2015). 25 protein-coding SNPs/Indels were found in KOLF2.1J but not in KOLF2-C1 (Table S2B). However, none had minor allele frequencies less than 0.001 or a CADD score greater than 30, suggesting that KOLF2.1J didn’t gain deleterious mutations after genomic editing and passaging. Four rare and deleterious variants were found in both KOLF2.1J and KOLF-C1 but were not classified as pathogenic (Table S2C).

### Comparison of KOLF2.1J to donor KOLF2 fibroblasts

In order to compare whether variants are new in the KOLF2.1J line vs the donor genome, we downloaded fibroblast exome sequencing from HipSci resource (https://www.hipsci.org/, ERZ486245). Reads covering the 40 variants of interest (Table S2D) were subset using samtools (v1.14) (Danecek et al. 2021). Reads were visually inspected using IGV 2.11.9 (J. T. Robinson et al. 2017) and compared to the generated KOLF2.1J WGS data generated for this manuscript. Variant presence was grouped into three categories: 1) Yes, meaning variant was present in fibroblast exome sequencing data at an allelic frequency near 50%, 2) No, meaning variant was not detected in fibroblast exome sequencing data and 3) Low allelic presence, meaning variant was present in <10% of total reads (Table S2D).

### Transcription Factor-NGN2 differentiation into cortical neurons

We expressed human NGN2 under a tetracycline-inducible promoter as previously described (Fernandopulle et al. 2018) using a PiggyBac system for delivery. iPSCs were transfected with PB-TO-hNGN2 vector in a 1:2 ratio (transposase:vector) using Lipofectamine Stem (Invitrogen), then selected after 24-48 hrs for 3 to 14 days with 8 µg/mL of puromycin (Sigma-Aldrich).

iPSCs with a stably integrated human NGN2 were single-cell dissociated using Accutase (Thermo Scientific) and plated at a concentration of 1.5×10^6^ per well of a Matrigel-coated (1:100, Corning) 6 well plate for 3 days with neuronal induction media (NIM: Knockout DMEM/F12, 1X N2 Supplement, 1X Non-Essential Amino Acids, 1X Glutamax (all from Thermo Fisher Scientific), 10 µM Rock inhibitor Y-27632 (SelleckChem) and 2 µg/mL Doxycycline (Sigma-Aldrich)). On day 3, 1.5×10^6^ cells were replated onto a poly-L-ornithine coated (PLO, Sigma-Aldrich) 6-well plate for 14 days using Brainphys (Stem Cell Technologies) 1X B27 Supplement (Thermo Fisher Scientific), 10 ng/mL BDNF (PeproTech), 10 ng/mL NT3 (PeproTech), 1 µg/mL Laminin (R&D) and 2 µg/mL Doxycycline (Sigma-Aldrich). For neuronal maintenance, half of the media was changed every 2-3 days.

### Transcription Factor-based differentiation into hNIL-expressing Lower Motor Neurons

Over-expression of NGN2-ISL1-LHX3 (hNIL) (Fernandopulle et al. 2018) was performed as described with a PiggyBac system for delivery. iPSCs were transfected with PB-TO-hNIL vector in a 1:2 ratio (transposase:vector) using Lipofectamine Stem (Invitrogen), then selected after 24-48 hours for 3 to 14 days with 8 µg/mL of puromycin (Sigma-Aldrich). The iPSCs with a stably integrated human NIL under a tetracycline-inducible promoter were exposed to 2 µg/mL doxycycline in neuronal induction medium (NIM). Briefly, on day 0 the iPSCs were single-cell dissociated using Accutase (Thermo Scientific) and plated at a concentration of 1.5×10^6^ per well of a Matrigel-coated (1:100, Corning) 10 cm dish in Essential 8 medium (Thermo Fisher Scientific) and 10 µM Rock inhibitor Y-27632 (Selleck Chem). On day 1, the media was changed with NIM: DMEM/F12, 1X N2, 1X Non-Essential Amino Acids, 1X Glutamax (all reagents were from Thermo Fisher Scientific), containing 10µM Rock inhibitor Y-27632, 2 µg/ml Doxycycline (Selleck Chem), and 0.2 µM Compound E (Stem Cell Technologies). On day 3, 1×10^6^ cells per were re-plated onto one well of a poly-L-ornithine (PLO, Sigma-Aldrich) and 15 µg/mL Laminin-coated (R&D) 6-well plate in NIM with 10µM Rock inhibitor Y-27632 (SelleckChem), 2 µg/mL Doxycycline (Sigma-Aldrich), 0.2 µM Compound E (Stem Cell Technologies), 1 µg/mL Laminin (R&D) and 40 µM BrdU (Sigma-Aldrich). On day 4, the media was changed to NIM with 1X B27 Supplement (Thermo Fisher Scientific), 1X Culture One Supplement (Thermo Fisher Scientific), 1 µg/mL Laminin (R&D), 20 ng/mL BDNF (PeproTech), 20 ng/mL GDNF (PeproTech) and 10 ng/mL NT3 (PeproTech). On day 7, half of the media was changed. On day 10, half of the media was changed with Neurobasal, 1X B27, 1X N2, 1X Culture One (all from Thermo Fisher Scientific), 40 ng/mL BDNF (PeproTech), 40 ng/mL GDNF (PeproTech), 20 ng/mL NT3 (PeproTech), 1 µg/mL Laminin (R&D) and 2 µg/mL Doxycycline (Sigma-Aldrich). For neuronal maintenance, half media was changed every 2-3 days.

### Directed differentiation to cortical and hypothalamic lineages

Prior to differentiation, hiPSC cell lines were maintained in mTeSR1 on geltrex (1:100, Thermo Fisher Scientific) and passaged with EDTA (0.5 µM, pH 8.0, Thermo Fisher, 15575-020) at 60-80% confluence. For each line, we confirmed that colonies had clearly defined borders and cultures lacked differentiated cells when viewed under a phase contrast microscope. Lines were passaged at least twice under these conditions before differentiation experiments were initiated, and lines were synchronized by adjusting split ratios so that the last passage was 3-4 days before plating for differentiation. For differentiation, hiPSC lines were dissociated to a single-cell suspension with TrypLE, counted, and plated onto coated plates. Cell lines were pooled for cortical differentiation, and grown separately for hypothalamic differentiation. For cortical and hypothalamic differentiation we followed published methods (Merkle, Maroof, et al. 2015; Kirwan, Jura, and Merkle 2017).

Briefly, dissociated hiPSCs were plated at 1×10^4^ cells/cm^2^ into 6-well plates in the presence of 10 µM ROCK inhibitor (Y-27632, Tocris Bioscience, Bristol, UK). Cortical and hypothalamic differentiation took place on a substrate of geltrex (1:100) in N2B27 media containing 500 ml Neurobasal-A (Thermo Fisher Scientific, cat. no 10888022), 500 ml DMEM/F12 with GlutaMAX (Thermo Fisher Scientific, cat. no 31331093), 10 ml Glutamax (Thermo Fisher Scientific, cat. no 35050038), 10 ml sodium bicarbonate (Thermo Fisher Scientific, cat. no 25080-094), 5 ml MEM Nonessential Amino Acids (Thermo Fisher Scientific, cat. no 11140035), 1 ml 200 mM ascorbic acid (Sigma-Aldrich, cat. no A4403) 10 ml 100x penicillin-streptomycin (Thermo Fisher Scientific, cat. no. 15140122), 20 ml 50x B27 supplement (Thermo Fisher Scientific, cat. no 17504044), 10 ml 100x N2 supplement (Thermo Fisher Scientific, cat. no 17502048). Patterning to forebrain progenitors in this medium was directed using 100 nM LDN-193189 (Stemgent, cat. no. 04-0074), 10 μM SB431542 (Sigma-Aldrich, cat. no. S4317), and 2 μM XAV939 (Stemgent, cat. no. 04-0046). The concentrations of these small molecules were adjusted over time as previously described (Kirwan, Jura, and Merkle 2017), and for hypothalamic differentiation the small molecules SAG (Fisher Scientific, cat. no. 56-666-01MG) and purmorphamine (Calbiochem, cat. no. 540220) were each added to a final concentration of 1 μM from day 2-8 of differentiation. To assess the efficiency of differentiation and the identity of progenitors, cells were plated in a separate 48-well plate at 4×10^5^ cells/well on day 11 and fixed for immunofluorescence assay on day 16.

### Cell dissociation for single-cell RNA sequencing

iNeurons and iLowerMotorneurons were washed once with PBS after 17 days of differentiation. The lines were then either differentiated in separate wells and pooled in a single tube at the end of differentiation, or the lines were pooled at the beginning of differentiation and were resuspended in 1x PBS-0.04% BSA (Jackson Immunoresearch) and washed 3 additional times with this solution after single-cell dissociation. Single-cell pellets were resuspended in 1x PBS-0.04% BSA, counted using an automated cell counter (Countess II) and the concentration was adjusted to 1×10^6^ cells/mL.

Cortical and hypothalamic neurons were washed once with 1X DPBS (Thermo Fisher, 14190-144) before adding TrypLE containing 20 units/ml of papain after 5 weeks of differentiation (34 days for cortical and 37 days for hypothalamic). Cultures were incubated at 37°C until the cells physically detached from each other when viewed under a phase contrast microscope and could be readily dissociated with a P1000 pipette. Enzyme mix was aspirated and cells were dissociated with a P1000 in DMEM:F12 (Thermo Fisher Scientific, 10565-018) supplemented with 10 µM Rock inhibitor, 33 μg/ml DNase I (Worthington, LK003170), and 45 uM Actinomycin D to block dissociation-induced transcription. The resulting cell suspension for each separate well was passed through a 40 µm cell strainer and brought to 10 ml in the dissociation solution centrifuged at 160x g for 5 minutes. For cortical differentiations, each well was uniquely barcoded using cholesterol-modified MULTIseq oligonucleotides to facilitate cell pooling during droplet-based single-cell cDNA library preparation based on a published protocol (McGinnis et al. 2019). After two additional washes, cells were resuspended in 1x DPBS containing 0.04% BSA (Sigma, A0281) and washed 3 additional times in 1x DPBS containing 0.04% BSA. Single-cell suspensions were counted using an automated cell counter (Countess II).

### Chromium 10x Genomics library and sequencing

For iPSC, iNeurons, and iLowerMotorneurons, single-cell RNA sequencing was performed using Chromium Single Cell 3’ Reagent kit V3.1 (PN-1000128) and 2.5×10^4^ cells per condition were loaded into the 10x Genomics chip G. For cortical and hypothalamic neurons, single-cell suspensions were processed by the Chromium Controller (10x Genomics) using Chromium Single Cell 3’ Reagent Kit v3 (PN-1000075) according to the manufacturer’s specifications. On average, 15,000 cells from each 10x reaction were directly loaded into one inlet of the 10x Genomics chip. Barcoded libraries were sequenced using the Illumina Novaseq 6000 (one lane per 10x chip position) with 75 bp paired-end reads to an average depth of approximately 5×10^4^ reads per cell.

### scRNA-seq data processing and quality control

Raw sequencing libraries were processed using 10x Genomics’ Cell Ranger platform (version 3.1). Reads were aligned and quantified to the 10x Genomics provided human reference genome (GRCh38, Ensembl 93). The samples were then grouped based on differentiation protocols and each group were processed independently in subsequent downstream analysis.

Droplets containing captured cells were called using the emptyDrops function from the DropletUtils R package (Lun et al. 2019), using varying UMI threshold per differentiation protocol groups and an FDR of 0.001. Low quality cells and outlier cells were then filtered based on the total unique molecular identifier (UMI) content, number of detected features/genes and fraction of mitochondrial content. Cells were discarded if their UMI content is more than ± 3 median absolute deviation (MAD) away from the median, or the detected features is more than ± 3 MAD away from the median, or the fraction of mitochondrial content is higher than 3 MAD from median. Gene expression levels were normalized using the logNormCounts function from the scran R package (Lun, McCarthy, and Marioni 2016), with size factors estimated using the computeSizeFactors function. Cells were assigned to cell cycle phases based on the expression of the G2/M and S phase markers (Tirosh et al. 2016) using the CellCycleScoring function from Seurat R package (Hao et al. 2021).

### Doublet detection

Doublet identification was done in two stages: at the individual sample level and across samples within the same differentiation protocol. First, doublets were detected at the individual sample level using the hybrid method (cxds_bcds_hybrid function with estNdbl parameter set to true) from the scds R package (Bais and Kostka 2020). This is followed by identification of ‘guilt-by-association’ doublets, where doublets were further identified if there is enrichment of scds’ hybrid-based doublets in the neighboring cells (number of neighbor = 3 for iPSC and 5 for other differentiations). Clustering was then performed on cells in each sample (see below), and this was repeated for each identified cluster to form smaller sub-clusters. Finally, cells were also classified as doublets if cells belong to sub-clusters containing more than 50% of Vireo-identified or MULTI-seq-identified doublets.

For doublet identification across samples within the same differentiation protocol, samples were first batch corrected (see below) into a single dataset per differentiation protocol, followed by two rounds of clustering to identify cell sub-clusters. Cells were classified as cross-sample doublets if they belonged to sub-clusters with an enriched fraction of per-sample doublets (>3 MAD away from the median). Cells which were classified as either per-sample doublets, cross-sample doublets, Vireo doublets or MULTIseq doublets were excluded from further downstream analysis.

### Batch correction and dimensionality reduction

For each differentiation protocol, samples were combined into a single dataset and corrected for batch effect using the fast MNN function from Batchelor R package (Haghverdi et al. 2018) on the first 50 principal components computed from highly variable genes (HVG). HVG were selected by fitting the mean-variance curve on the normalized gene expression across all samples within a differentiation group with modelGeneVar from scran R package and filtering for genes which have higher variance than the fitted trend. Mitochondrial genes and ribosomal genes for large and small ribosomal subunits were excluded from mean-variance curve fitting as these genes have both high variance and expression. For visualization, Uniform Manifold Approximation and Projection (UMAP) two-dimensional embedding (McInnes et al. 2018) were calculated from the corrected principal component with the following settings: spread = 1 and minimum distance = 0.4.

### Clustering and annotation

Cells were grouped into clusters for each differentiation protocol using the community detection-based Louvain clustering method. Briefly, shared nearest-neighbor graphs were constructed from the 50 corrected principal components, followed by clustering using the Louvain method (cluster_louvain function) from the igraph R package (Nepusz 2006). Each cluster was then manually annotated with cell type based on a list of curated markers (Table S5) and further assigned into one of four cell type groups for evaluating differentiation efficiency of each cell sub-line.

### Cell sub-line and replicate demultiplexing

Cell sub-line identity was inferred based on genotype information using 10x Genomics VarTrix and Vireo tools (Huang, McCarthy, and Stegle 2019). Variant count matrices for captured cells were produced by VarTrix using aligned reads from Cell Ranger output and variants called from whole genome sequencing data (see above). Cell sub-line identity for captured cells were then determined with Vireo using the variant count matrix and variant information. Only variants from 7 cell sub-lines were utilized for demultiplexing as both NN0003932 and NN0004297 (denoted as NN_combined) were derived from the same donor.

Replicate demultiplexing of MULTI-seq labeled samples was performed using the deMULTIplex R package. Briefly, MULTI-seq barcode reads from captured cells were aligned to the MULTI-seq barcodes used for labeling each sample, followed by read deduplication based on UMI and generation of MULTIseq barcode count matrix. Replicate classification was then performed on the barcode count matrix iteratively until there are no negative classified cells, followed by a negative-cell reclassification to recover incorrectly classified negative cells.

### Immunofluorescence and imaging

Cells were fixed in 4% paraformaldehyde for 10 min at room temperature and washed 3x in PBS. Afterwards, the cells were incubated in blocking solution composed of PBS, 0.2% Triton-X and 4% donkey serum for 1 hr at 4°C. Primary antibody, anti-FOXA2 (R&D Systems AF2400) and anti-Lmx1 (Merck Millipore, AB10533) were added 1:100 into blocking solution (PBS, 5 % donkey serum, 0.1 % Triton X10) and the cells were incubated for 2 hr at room temperature. Samples were washed in 3x blocking solution before the addition of 1:500 donkey anti-goat IgG secondary antibody, Alexa Fluor® 561 (Thermo Fisher Scientific) and 1:500 donkey anti-goat IgG (H+L) secondary antibody, Alexa Fluor® 647 (Thermo Fisher Scientific) to the cells for 1 hr at room temperature. Samples were washed 3x in blocking solution and 1x in PBS. 1:1000 Hoechst 33342 (Thermo Fisher Scientific) was added to the cells in PBS before imaging them.

---

**Figure S1.**
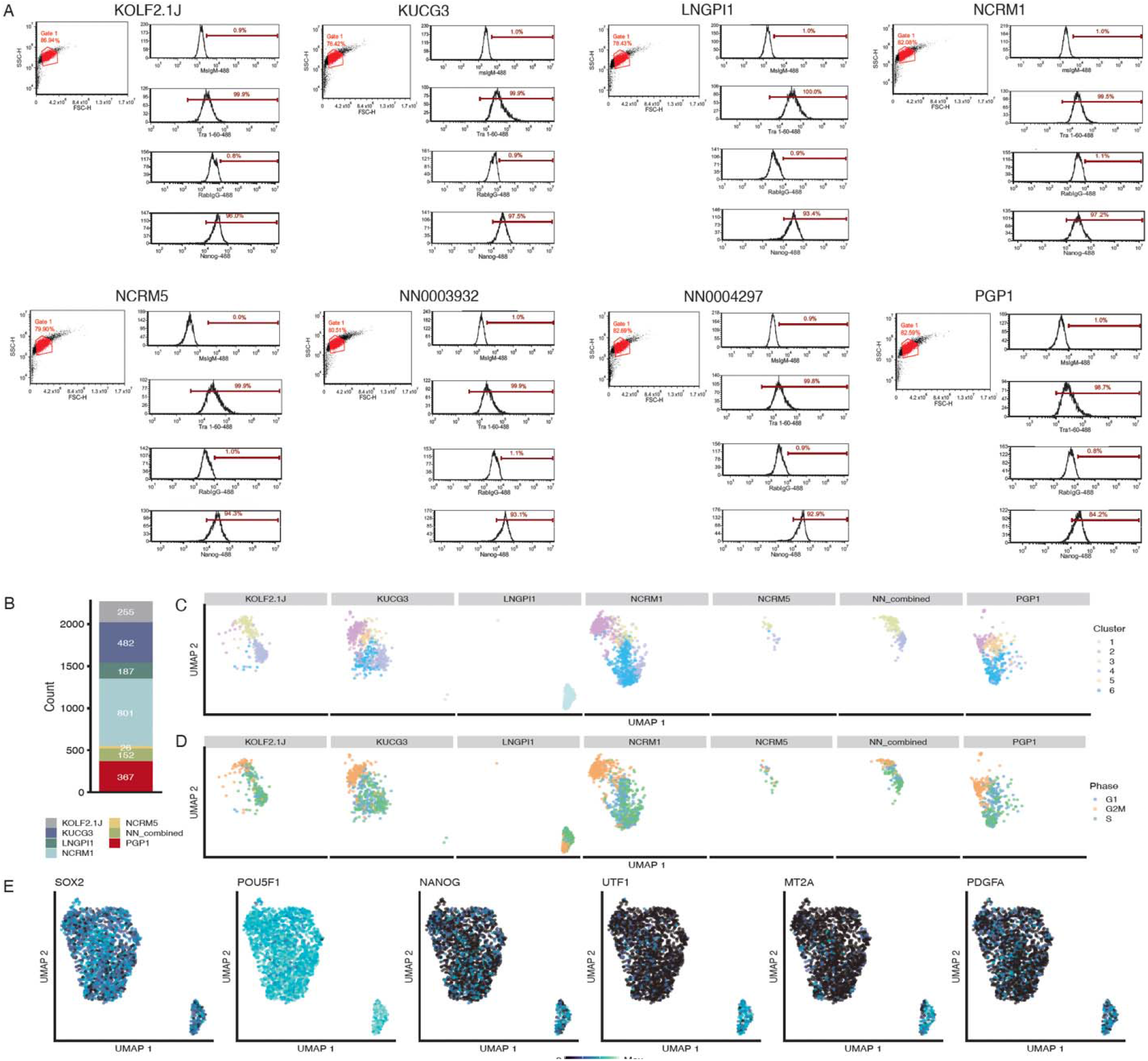
Flow cytometric and single-cell transcriptomic analysis of pluripotency markers. (A) For each line, live iPSCs were analyzed by flow cytometry after selection by a FSC/SSC gate to quantify the percentage of cells expressing TRA-1-60+ or NANOG. Results for each cell line were: KOLF2.1J (99.9% TRA-1-60+, 96% NANOG+), KUCG3 (99.9% TRA-1-60+, 97.5% NANOG+), LNGPI1 (100% TRA-1-60+, 93.4% NANOG+), NCRM1 (99.5% TRA-160+, 97.2% NANOG+), NCRM5 (99.9% TRA-1-60+, 94.3% NANOG+), NN0003932 (99.9% TRA-1-60+, 93.1% NANOG+), NN0004297 (99.8% TRA-1-60+, 92.9% NANOG+), PGP1 (98.7% TRA-1-60+, 84.2% NANOG+). (B) Bar plot displaying the proportion of each genetically distinct sub-line within the analyzed single-cell pool. (C-D) UMAP plots of all iPSC cells faceted by sub-line and colored by clusters identified (C), or predicted cell cycle phase (D) (E) UMAP plots of all iPSC cells faceted by sub-line and colored by expression of selected genes indicative of pluripotency (left three) or predictors of poor neuronal differentiation efficiency (right three). Relates to Figure 1.

**Figure S2.**
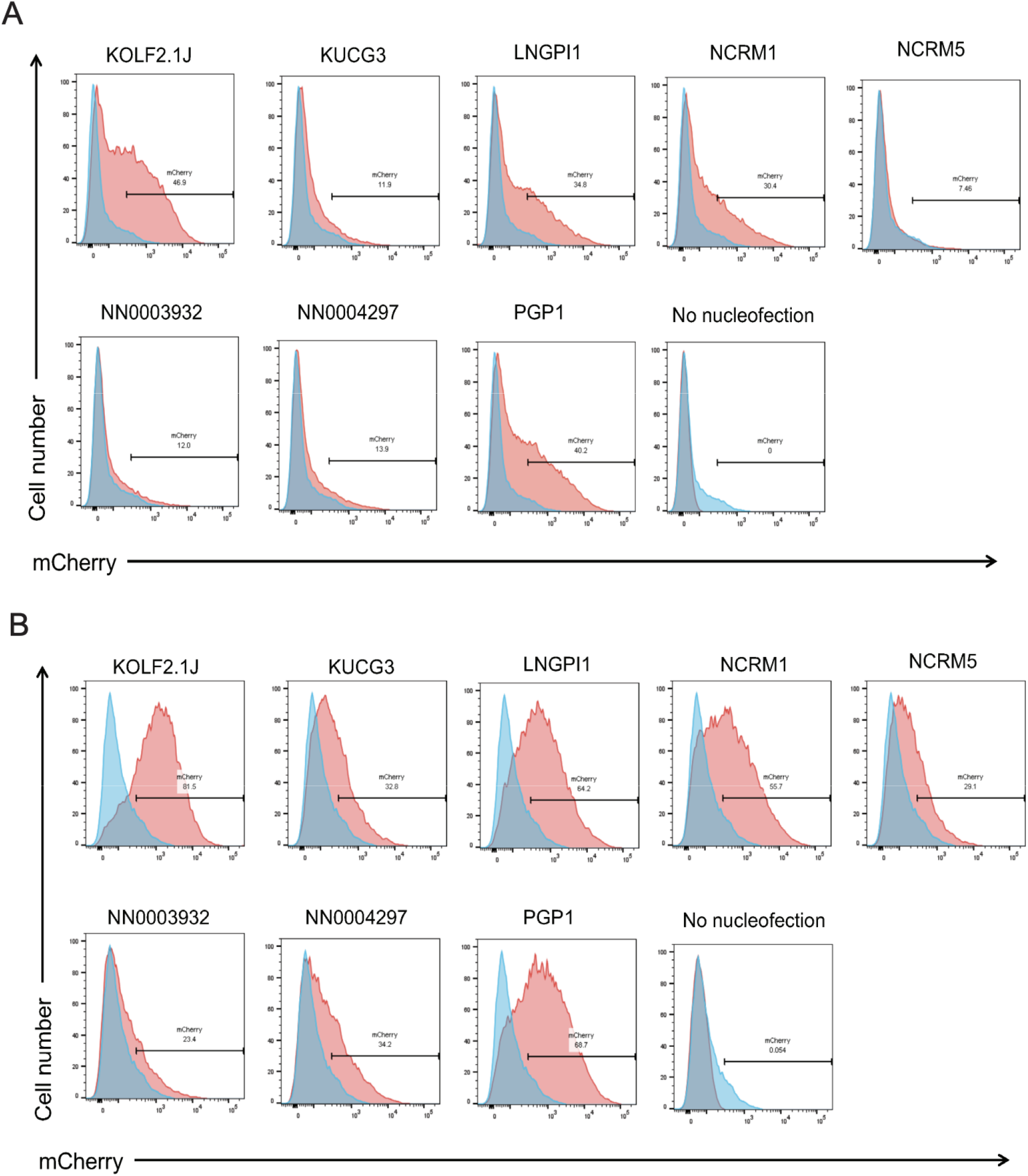
Fluorescent assay for baseline and DNA damage-induced p53 pathway activity. Episomal p53 reporter assay assessing p53 function in each candidate cell sub-line (red) or *TP53* knockout cells (blue) in the presence of vehicle control (A) or the DNA damaging agent doxorubicin (B). Relates to Figure 1

**Figure S3.**
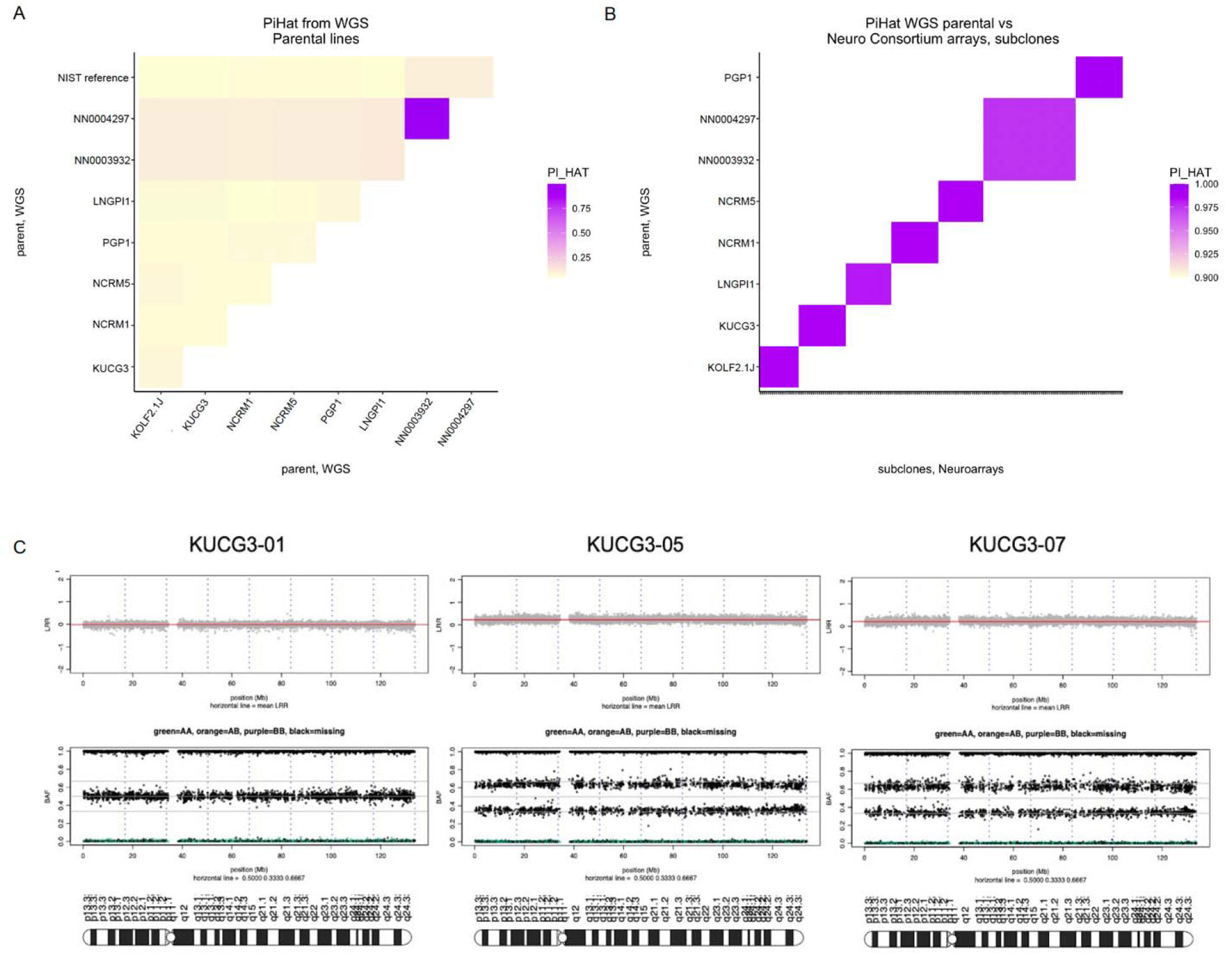
Genomic fidelity of eight candidate cell lines. (A) Pairwise comparisons of pi-hat (color scale) between iPSC lines from WGS data show that only the two lines derived from the same donor are directly related. (B) Pairwise comparison of pi-hat (color scale) from WGS for all lines on the y axis against subclones from DNA microarray-based genotyping on the x-axis (compressed) indicates high genomic fidelity between parental lines and their gene-edited derivatives. (C) After editing at the *TIMP3* locus, two clones of sub-line KUCG3 showed a duplication of chromosome 12 with an unaffected clone (KUCG-01) shown for comparison. Upper plots show Log R ratio (LRR) where the mean LRR (red line) is 0 for the normal clone and >1 for the abnormal clones, and middle panels show the B allele frequency (BAF) for bi-allelic probes along the arrays with evidence of duplicated alleles across the chromosome. Ideograms of chr12 are shown below each image for scale. (D). Ideogram of chromosome 22 with the *TIMP3* gene in 22q12.3 indicated by a red bar, using the UCSC Genome Browser. (E-F) NeuroArray genotyping revealed Chr22 CN-LOH from chr22q12.3-ter in clones derived from the NRCM1 sub-line (E), and the PGP1 sub-line (F). Unaffected clones from each sub-line are shown for comparison. Upper plots show Log R ratio (LRR) where the mean LRR (red line) is 0 across chromosome 22, and middle plots show the B allele frequency (BAF) for bi-allelic probes along the arrays. Ideograms of chr22 are shown below each image for scale. Relates to Figure 3.

**Figure S4.**
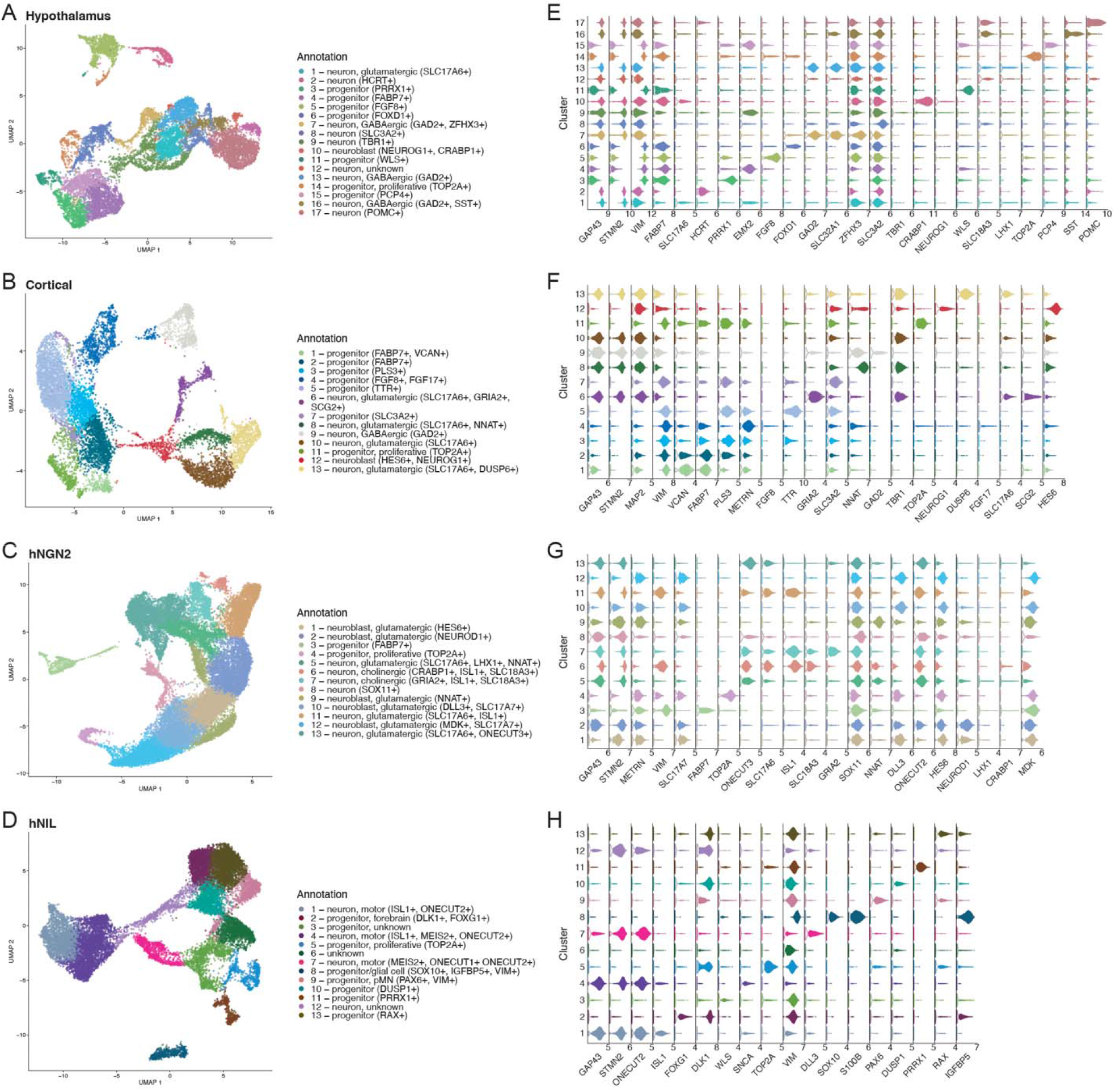
Cluster identity and marker expression across differentiation protocols. (A-D). UMAP plots for each of the four differentiation protocols evaluated in this study; hypothalamus (A), cortical (B), hNGN2 (C) and hNIL (D), colored by cluster identity. (E-H). Beeswarm plots showing expression of informative differentiation-specific curated markers for each of the four differentiation protocols; hypothalamus (E), cortical (F), hNGN2 (G) and hNIL (H). Relates to Figure 4.

**Figure S5.**
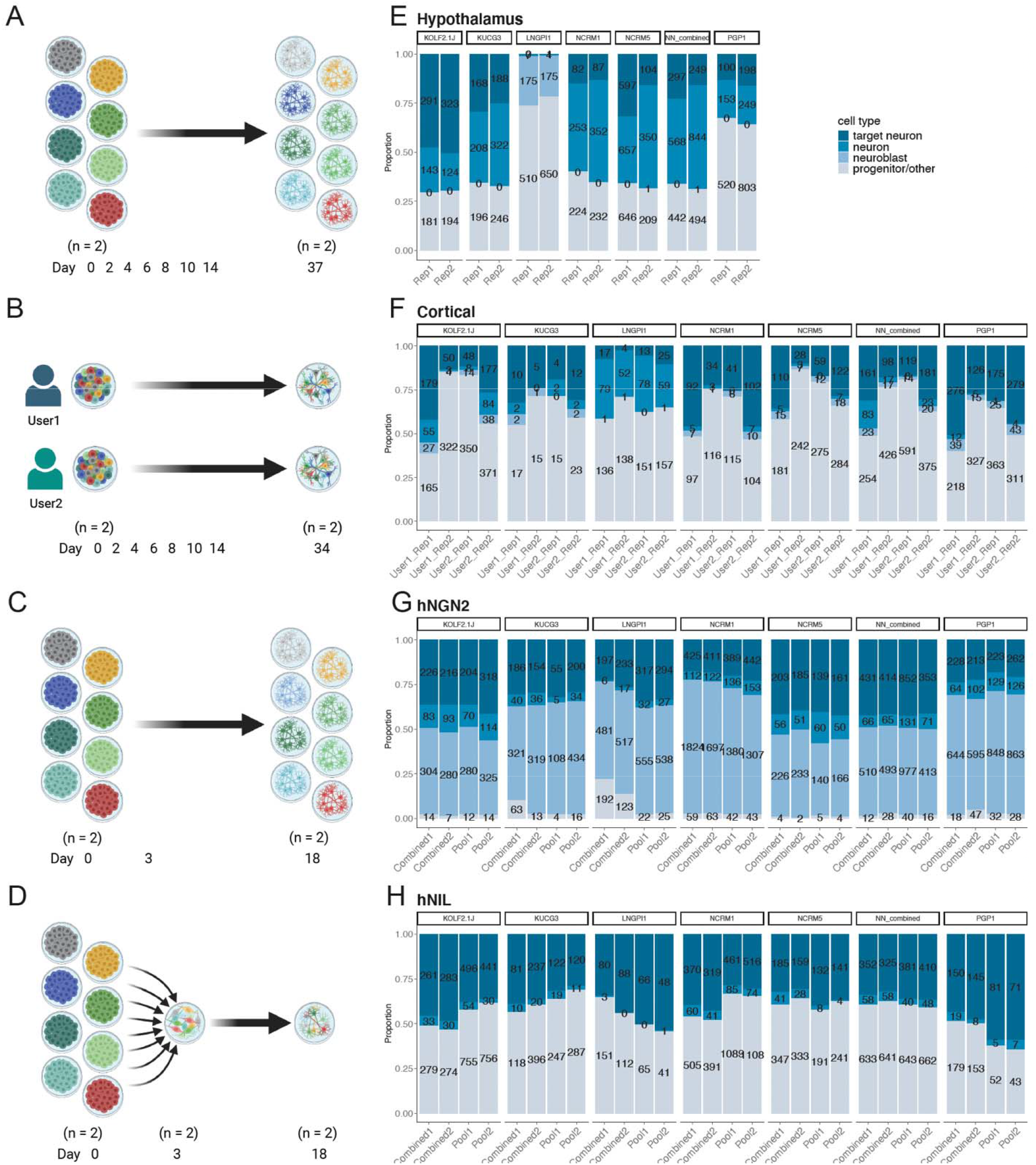
Differentiation performance of candidate cell sub-lines across replicates. (A-D). Schematic of experimental replicate structure for the four differentiation protocols evaluated in this study. For hypothalamic differentiation, the differentiations were performed individually for each cell sub-line with each sub-line having two replicates. (A); while for cortical differentiation, the differentiations were performed with all the sub-lines pooled together by two different users, with each user having two replicates (B). In the hNGN2 and hNIL differentiations, cells sub-lines were differentiated both individually for each cell sub-line (C, indicated as Combined in G and H) and with all sub-lines pooled together after initial transcription factor induction at day 3 (D, indicated as Pool in G and H), with each differentiation method having two replicates. (E-H). Bar plots showing proportion of cells assigned to each cell type per replicate, faceted by cell sub-line, for each of the four differentiation protocols – hypothalamus (E), cortical (F), hNGN2 (G) and hNIL (H). Schematics created with Biorender.com. Relates to Figure 4.

**Figure S6.**
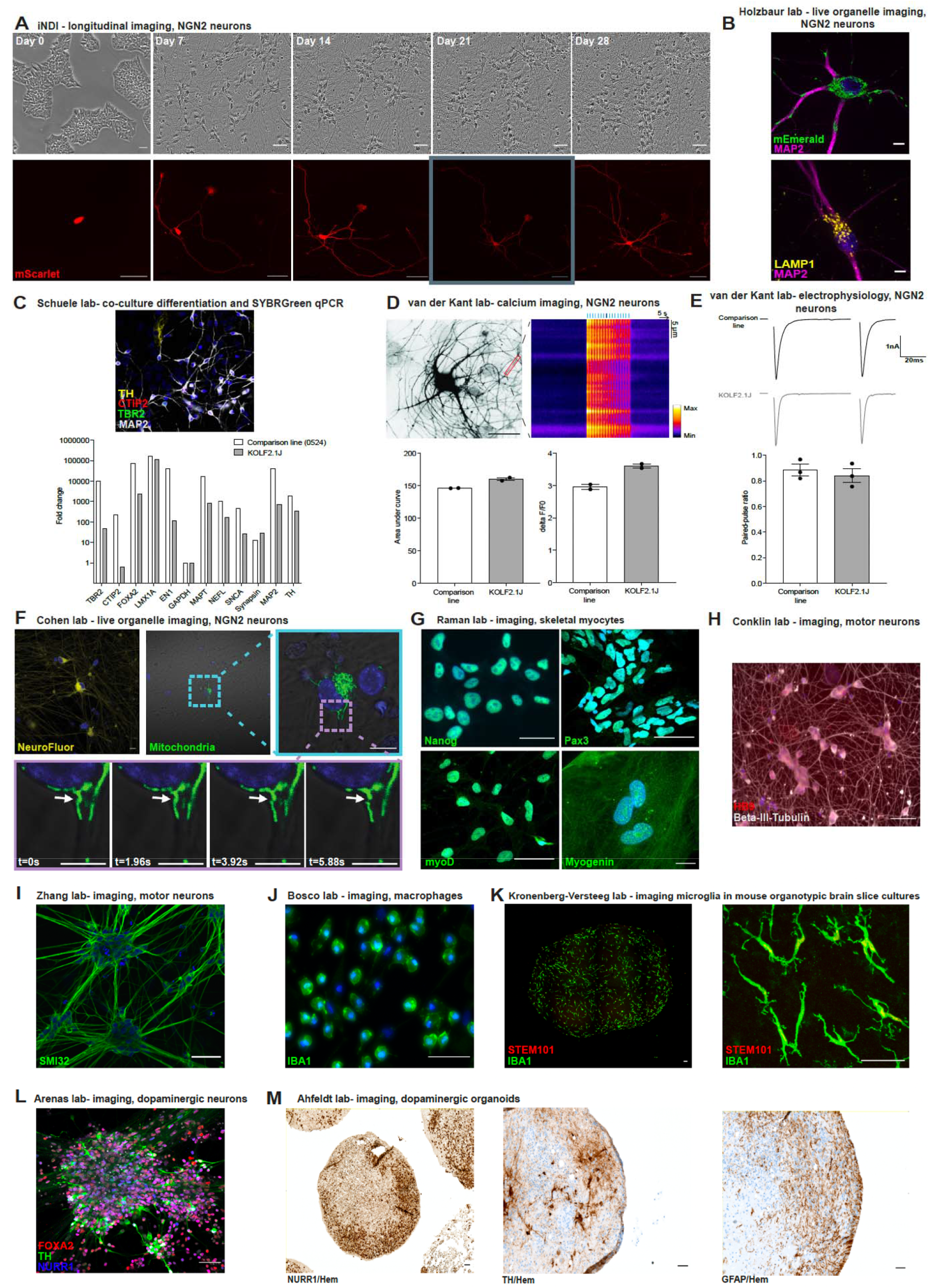
Differentiation and functional evaluation of KOLF2.1J by diverse methods. Unless indicated in the subpanel, scale bars represent 50 mm and nuclear markers are in blue. Relates to Figure 6. (A) Differentiation of KOLF2.1J into NGN2-expressing cortical neurons. Bright field images of KOLF2.1J-hNGN2 throughout differentiation (top row). Time-course of a neuron transduced with cytosolic mScarlet to identify neurites (bottom row). (B) Live organelle imaging of KOLF2.1J differentiated into NGN2-expressing cortical neurons. Cells were stained for MAP2 (purple) and organelles were visualized by an expressed plasmid for mitochondria (mEmerald-mito, green, upper) immunostaining for LAMP1 for lysosomes (yellow, lower). Scale bar indicates 5 µm. (C) Mixed neuronal culture of cortical, striatal, and dopaminergic KOLF2.1J-derived neurons, co-cultured with primary human astrocytes. Cells immuno-positive for CTIP2 (red), TBR2 (green), MAP2 (gray), and tyrosine hydroxylase (TH; yellow) were observed (top). SYBR Green qPCR expression array of iPSC-derived dopaminergic neurons from KOLF2.1 and the 0524 comparison line (bottom graph). (D) Representative example of a KOLF2.1J-derived neuron during calcium imaging (upper left). Kymograph of traces of intracellular calcium (Fluo5-AM) levels upon repetitive electrical stimulation (blue bars) in KOLF2.1J (upper right) indicating robust calcium influx upon repetitive stimulation similar to the BioniC13 comparison line. (E) Typical example traces of paired-pulse recordings for the BioniC13 and the KOLF2.1J line (upper). Paired-pulse ratio at 10 Hz measured as the ratio of the second eEPSC over the first eEPSC amplitude (lower) does not differ between BioniC13 and KOLF2.1J. N=3 differentiations. (F) Time-lapse images of mitochondrial movement in KOLF2.1J cells differentiated into NGN2-expressing cortical neurons. Images were taken every 1.96 seconds. Scale bar indicates 5 µm. (G) KOLF2.1J cells stained for the pluripotency marker Nanog (top left) were differentiated into skeletal myocytes as indicated by staining for PAX3 in pre-myogenic progenitors at d15 (top right), and myoblasts as indicated by staining for myoD at d20 (bottom left), and myogenin at d30 (bottom right). (H) KOLF2.1J iPSCs were differentiated into motor neurons using an inducible transgenic system. Beta-III-tubulin (TUJ1, grey; cell body/axons) and HB9 (red; nuclear) expression indicates cells that successfully differentiated by d10. (I) KOLF2.1J neurons differentiated to motor neurons and stained positively for SMI32 (green). (J) KOLF2.1J-derived macrophages at d7 of differentiation express the myeloid marker ionized calcium-binding adapter molecule 1 (IBA1; green). (K) KOLF2.1J-derived microglial precursors at d21 post-engraftment onto mouse organotypic brain slice cultures at low (left) or high (right) magnification. Cultures were immunostained for IBA1 (green) and STEM101 (red, against human nuclear protein Ku80). (L) Some KOLF2.1J-derived midbrain dopaminergic neurons were triple-positive for TH (green), FOXA2 (red), and NURR1 (blue) at d28. (M) A d35 midbrain organoid section indicating NURR1-positive nuclei (brown, left). d100 midbrain organoid sections are depicted in center and right images. Dopaminergic (DA) neurons are indicated by tyrosine hydroxylase-positive staining (TH; center), and astrocytes are indicated by GFAP-positive staining (right) in brown. Counterstain is hematoxylin (Hem).

