## Supplemental Information for "A reference induced pluripotent stem cell line for large-scale collaborative studies"

**Supplemental Methods**

*Sub-cloning*

All iPS cell lines were maintained feeder-free in complete StemFlex media (Thermo) without the addition of antibiotics. StemFlex was replaced every other day and cells were incubated at 37°C/5%CO_2_. Upon addition of the provided 10x StemFlex supplement, complete StemFlex media was stored at 4°C for less than one month, and only the volume needed for a given experiment was removed and pre-warmed to room temperature before use. Following thawing and single-cell plating, RevitaCell (Thermo) at a 1x concentration was included in the media for one day to improve survival of iPS cells in a single-cell state.

To generate clonal sublines, iPSC lines were first thawed and plated in Matrigel (Corning)-coated 24-well plates (for low cell number vials) or Synthemax (Corning)-coated 6-well plates (for high cell number vials). Cells were further expanded into Synthemax-coated 6-well plates. The cells were allowed to replicate over several days until they reached near confluency. Spent media was collected and tested by qPCR for Mycoplasma by the UCONN Stem Cell Core. Cells were detached as a single-cell suspension by gentle scraping after a 10 minute incubation at 37°C in Accutase (Stemcell Technologies). To generate well separated iPS cell colonies, 1000 cells were plated onto a Synthemax-coated 10cm dish and incubated for 9-10 days. This resulted in a variable total number of iPS cell colonies per cell line (Table S1A). For each cell line, 8 colonies were manually picked into wells of a Matrigel-coated 24-well plate and grown to near confluence. During this incubation, cell morphology was observed daily under the microscope. Of these 8 clones, 3 clones per cell line with minimal or no observable cell differentiation were further expanded into Synthemax-coated wells of a 6-well plate. This provided sufficient cells to both cryopreserve each subline, and prepare cells for G-band karyotyping. After a 7 minute 37°C Accutase treatment, cells were gently scraped in Knockout serum replacement (Thermo) containing 10% DMSO, aliquoted into cryovials, and stored at -80°C for one day before being moved to vapor-phase liquid nitrogen storage. The remaining cells were washed in DPBS-/- and cell pellets were stored at -80°C.

*Expansion of selected sublines*

Each of the 8 selected iPS cell lines were expanded into 2 x 96-well Matrix plates (Thermo). Briefly, cells were thawed into a single well each of a Synthemax-coated 6-well plate. Once confluent, the cells were clump passaged using ReLeSR into a Synthemax-coated 10cm plate. These were subsequently expanded by clump passaging into 6 x 10cm plates. Cells were treated with Accutase, gently scraped in cold Knockout serum replacement with 10% DMSO, pooled, and distributed to 192 individual minivials of 2 open-capped 96-well Matrix plates. Minivials were topped with 100 µl of sterile paraffin oil (Sigma), frozen at -80°C for one day, and stored in vapor-phase liquid nitrogen.

The expanded Matrix plate stocks for all 8 lines were tested as follows. One minivial was removed from liquid nitrogen and the cells were thawed and transferred, leaving the oil behind, into a well of a 96-well V-bottom plate. The V-bottom plate was spun at 300xg for 5 min, and the supernatant was removed. The cell pellet was gently resuspended in complete StemFlex with RevitaCell and incubated in a single well of a Synthemax-coated 6-well plate. After incubation to near confluency, the cells were clump passaged at a 1:10 dilution with ReLeSR into multiple wells of a new Synthemax-coated 6-well plate for mock editing, nucleic acid preparation, *Mycoplasma* testing, and Directional Genomic Hybridization, as described below.

*Genetic correction of the KOLF2-C1 cell line*

CRISPR-Cas9 editing of KOLF2-C1 hiPSCs (p9) was performed to correct a 19bp pathogenic deletion in one copy of the ARID2 gene (Hildebrandt et al. 2019). Cells were grown in a 5% CO2, 37C incubator in StemFlex media (ThermoFisher) on Synthemax-treated wells (Synthemax II; Corning) and dissociated to single cells with Accutase (Stemcell Technologies). In a volume of 0.1 mL P3 Primary Cell P3 buffer (Lonza), 8 x 10^5^ KOLF2-C1 cells were nucleofected using the Lonza Amaxa 4D nucleofector (program CA137) with 20 mg Cas9 protein (HiFi v3; IDT), 16 mg single guide RNA targeted to the deleted ARID2 allele (5’-AAAAGATCACTTGCTAATGCCGG…-3’; chemically-modified; Synthego), and 200 pmol of a 120-mer oligonucleotide repair template (desalted Ultramer; IDT) corresponding to wild-type sequence (5’-ACGTATGCACTCTCCTATCAAATGAAAGCAAGCACGTCATGCAACTTGAAAAAGATCCTAAAATCATCACTTTACTACTTGCTAATGCCGGGGTGTTTGACGACAGTAAGTTTTAAGCTG-3’; deleted sequence underlined). Cas9 ribonucleoprotein complexes were assembled *in vitro* for 30 min prior to nucleofection. The nucleofected cells were seeded onto one well of a Synthemax-treated 6-well plate in StemFlex media containing 1X RevitaCell (ThermoFisher) and 30 mM Alt-R HDR Enhancer (IDT) and cultured at 32℃ (cold shock) for three days. After 24 hrs, the media was changed to remove HDR Enhancer and RevitaCell. At 72 hrs, the media was replenished and the cells were cultured to confluency at 37℃. Following the cloning of single-cell derived colonies, confluent wells of replicate 96-well plates were archived and lysed for genotyping. The target region was amplified by PCR (950 bp amplicon; forward primer, 5’-TTGGCAATGATGGCCAAATGGTATG-3’; reverse primer, 5’-AAAACCCACAACTAGCAAACCCTAC-3’) using LongAmp polymerase (NEB) and subjected to Sanger sequencing (forward primer, 5’-GTCAAAGTTATGGGCTGTCC-3’; reverse primer, 5’-GTTGACAAACAAAAAGTACTTTCTCC-3’). From a screen of 96 clones, four ARID2-corrected clones were identified (A2, A11, E3, H5) and expanded for karyotyping. All clones showed a normal karyotype, and clone A2 was selected for further characterization (designated KOLF2.1J).

*G-band karyotyping*

To prepare cells for G band karyotyping, sublines were clump passaged using ReLeSR (STEMCELL Technologies) 1:2 into new wells of a 6-well plate and incubated overnight. Sublines from parental cells KOLF2.1J, NCRM1, NCRM5, and PGP1 were prepared for karyotyping by the Cytogenetics Laboratory at The Jackson Laboratory for Genomic Medicine. Specific sublines KUCG3-C1, LNGPI1-C1, NN0003932-C3, and NN0004297-C1 were processed by the Cellular Engineering laboratory at The Jackson Laboratory for Genomic Medicine according to the fixation protocol recommended by KromaTiD, and shipped to KromaTiD for karyotyping.

Briefly, colcemid (Thermo) was added to a final concentration of 0.1µg/ml and cells were incubated for 4 hrs. Cells were then treated with Accutase for 10 min, detached by gentle scraping in DPBS-/-, and collected in a 15ml polystyrene conical tube. The single-cell suspension was spun for 10 min at 1000rpm at RT. The supernatant was mostly removed, leaving 1ml remaining on the cell pellet. The cell pellet was resuspended by vigorous flicking and 6ml of RT 75mM KCl was added to induce osmotic swelling. The cell mixture was incubated at RT and inverted slowly every 5 min. After 30 min, 1.5 ml of freshly made fixative (3:1 methanol:acetic acid) was added, mixed by slow inversion, and the swollen cells were spun at 1000rpm for 10 min at RT. The supernatant was mostly removed, leaving 1 ml remaining on the cell pellet. The cell pellet was resuspended by vigorous flicking and 5ml of fixative was added in a dropwise fashion while the cells were gently vortexed. Cells were incubated in fixative for 20 min at RT, then spun down at 1000rpm for 10 min at RT, before transfer to a cryovial for shipment. For samples analyzed by the JAX Cytogenetics Laboratory, 15 cells (metaphases) were counted and sexed, and 5 cells were analyzed for each sample. For samples analyzed by KromaTiD, 20 cells (metaphases) were analyzed for each sample.

*Directional Genomic Hybridization (dGH™)*

To assess the genomic stability of edited samples, KromaTiD’s directional genomic hybridization (dGH™) was performed using an assay consisting of dGH chromatid paints for human chromosomes 1, 2 and 3 in a single color. For 8 samples of human iPSCs, live, actively dividing cultures were prepared for dGH by adding 5.0 mM 5-bromo-deoxyuridine (BrdU) and 1.0 mM 5-bromo-deoxycytidine (BrdC) to the cell culture media for the duration of one division cycle, as described elsewhere (Ray et al. 2013). At 4 h prior to harvest, colcemid was added to a final concentration of 0.1 µg/ml to each sample. The samples were harvested and fixed in 3:1 methanol/acetic acid, and metaphase spreads were prepared for each sample using standard cytogenetic techniques (Howe, Umrigar, and Tsien 2014). To remove the newly replicated strand in each metaphase chromosome, prepared slides of metaphase spreads singly substituted with BrdU and BrdC were submersed in Hoechst 33258 (Millipore Sigma) for 15 min, selectively photolysed using a SpectroLinker XL 1500 UV Crosslinker equipped with 365nm UV bulbs for 35 min, followed by exonucleolytic degradation of the nicked DNA with Exonuclease III (New England Biolabs) for 35 min. A hybridization mixture consisting of probe and hybridization buffer was applied to the slides, cover-slipped, and sealed with rubber cement. Slides were denatured *in situ* at 68°C for 3 min, incubated overnight at 37°C, and then washed 5X in 2× SSC at 43°C. Slides were mounted in Vectashield with DAPI, and images of metaphase spreads were acquired on an Applied Spectral Imaging Harmony system using a Zeiss Axio Imager.Z2m microscope equipped with a Basler acA2440-35um camera, and a Lumencor SOLA SE-FISH LED light source, using a 100x objective. A total of 200 metaphase spreads from each sample were scored for genomic structural rearrangements including translocations, inversions, sister chromatid exchanges, insertions, deletions and other complex structural rearrangements involving the painted chromosomes, as well for dicentric or acentric chromosomes present in the metaphase spread. One sample had insufficient material to score 200 spreads. For this sample, 181 spreads were scored. dGH sample preparation was performed on a single cell line to establish the timing for analog addition and harvest, with the belief that all 8 cell lines had similar growth characteristics. However, the cell cycle timing proved to be different enough between the 8 cell lines to affect dGH. This resulted in a percentage of scorable cells that was quite low for some samples, requiring use of many prepared slides to achieve the desired scored sample size. Note that for KUCG3, the entire sample was consumed before 200 scorable cells could be observed, and a total of 181 spreads were scored. The samples ranged from 9 scored metaphases per slide (LNGPI1) to 67 scored metaphases per slide (NCRM1).

Chromosomes 1, 2, and 3 combined comprise 23% of the human genome, and results can be used to extrapolate event rates across the whole genome. Results were reported as the rate of total events per sample, with a breakdown of event rates by event type.

**Differentiation of KOLF2.1J to different cellular phenotypes**

*Three germ layer differentiation (Bu lab)*

Cell Culture

KOLF2.1J iPSCs were thawed and plated onto a Matrigel-coated dish in mTSER1 (Stemcell Technologies) containing 10 mM rock inhibitor Y27632 (Sigma-Aldrich). Medium change with mTSeR1 without a rock inhibitor was performed daily after the first 24 h. Pluripotency of KOLF2.1J iPSCs was evaluated by three germ layer differentiation using STEMdiff Trilineage Differentiation kit (Stemcell Technologies) following the manufacturer’s instructions. Cells were passaged with Accutase (Stemcell Technologies) when approximately 70% confluent and plated onto Matrigel-coated 24-well plates with mTeSR1 containing rock inhibitor at a density of 10,000 cells/well. After 24h, lineage-specific differentiation medium was exchanged daily for either 5 days (endoderm and mesoderm) or 7 days (ectoderm).

Immunostaining

Differentiation was confirmed by immunostaining specific markers of each germ layer (Ectoderm: Nestin; Mesoderm: Brachyury; Endoderm: SOX17). Cells were fixed in 4% PFA in PBS for 10 min, permeabilized with 0.2% Triton X-100 in PBS for 10 min and blocked for 60 min in blocking buffer (1% bovine serum albumin in PBS), at room temperature. Cells were then incubated overnight at 4ºC with primary antibodies diluted in blocking buffer: mouse anti-Nestin (1:300, Abcam); goat anti-Brachyury (1:300, R&D); mouse anti-SOX17 (1:300, Abcam); followed by incubation with Alexa fluor anti-goat or anti-mouse IgG secondary antibody diluted in blocking buffer, at room temperature for 1 h in the dark (1:400, ThermoFisher). Images were acquired using a Keyence digital microscope.

Cell Quantification

Trilineage differentiation efficiency was quantified to establish a comparison with a commercial APOE3 iPSC line from ALSTEM. Images (n=2-5) from separate (2 or 3) endoderm, mesoderm, and ectoderm differentiation experiments were quantified using Fiji to determine the percentage of Sox17-positive cells (endoderm), Brachyury-positive cells (mesoderm) and Nestin-positive cells (ectoderm; Figure 6A).

*Cortical neurons (iNDI)*

Cell Culture

KOLF2.1J iPSCs were differentiated into hNGN2-expressing cortical neurons using an established protocol (Fernandopulle et al. 2018), as described above. For the neurite outgrowth experiments, cytosolic mScarlet (MK-EF1a-mScarlet) was transduced in KOLF2.1J with stably integrated human NGN2 using a lentivirus expressing cytosolic mScarlet to identify neurite (Tian et al. 2019). The course of the neuron differentiation was recorded by scanning using an Incucyte® S3 Live-Cell Analysis System with a 20X objective. Imaging was performed every 24 h at 37°C for 28 days. Phase images were acquired for every time point (Figure S6A).

*Cortical neurons (Wainger lab)*

Cell culture

To assess neuronal differentiation efficiency, KOLF2.1J was compared to two control iPSC lines in the NINDS Human Cell and Data Repository (ND50003, ND50004) and a third control line (11a (Boulting et al. 2011)). iPSCs were plated in T75 flasks (Fisher Scientific) coated with Vitronectin XF (Stemcell Technologies) according to manufacturer instructions. iPSCs were cultured in mTeSR plus (Stemcell Technologies) and media was changed every 2-3 days.

To generate iPSCs with a stable tet-inducible NGN2 construct, 2.75µg PiggyBac transposase and 2.75µg tet-NGN2 (both from Michael Ward) were nucleofected using a Nucleofector I (Lonza, protocol A-23) and the Human Stem Cell Nucleofector Kit 1 (Lonza). To improve survival after nucleofection, mTeSR plus was supplemented with CET (Chen et al. 2021) (50nM Chroman 1, MedChem Express; 5uM Emricasan, Selleckchem; 0.7uM Trans-ISRIB, Tocris). The next day, CET was removed and replaced with puromycin (InvivoGen, 10ug/mL) to select for tet-NGN2 integration. Cells were continually supplemented with puromycin until all cells were positive for BFP2, indicating all remaining cells had integrated the tet-NGN2 construct.

To differentiate iPSCs into i^3^ neurons, 12 million iPSCs were diluted in induction media (DMEM/F12, Life Technologies; N2 supplement, Gibco; NEAA, Corning; Glutamax, Thermo Fisher Scientific; Penicillin/Streptomycin, Life Technologies; 2µg/mL Doxycycline, Millipore Sigma) supplemented with CET and plated into a T175 flask (Fisher Scientific) coated with Vitronectin XF. The next day, media was exchanged for fresh induction media without CET. The following day differentiated i^3^ neurons were treated with accutase (Thermo Fisher Scientific), counted, and frozen in batches of 1 million cells/vial (500 µL freezing media: 40% FBS, Hyclone; 10% DMSO, Millipore Sigma; 50% induction media; CET).

Differentiated i^3^ neurons were plated at 40,000 cells/well in induction media with CET in 96-well plates [Cellvis coated with PDL (Millipore Sigma) and Laminin (Life Technologies) according to manufacturer recommendations]. The next day, induction media was exchanged with cortical neuron media (Neurobasal, Life Technologies; B27 supplement, Gibco; Glutamax, Thermo Fisher Scientific; Penicillin/Streptomycin, Life Technologies; 50mM NaCl, Millipore Sigma; 10ng/mL NT-3, Peprotech; 10ng/mL BDNF, Life Technologies) supplemented with 10uM U/FdU (Millipore Sigma). Two days later, media was exchanged for fresh cortical neuron media without U/FdU. Thereafter, half the media was exchanged every 2-3 days.

Immunostaining

Cells were fixed 7 days after thawing using 4% formaldehyde (Life Technologies) for 15 min. Cells were then blocked and permeabilized for 1 h in PBT (1xPBS, Gibco; 0.5% Triton-X, Millipore Sigma; 1% BSA, Millipore Sigma) and then incubated overnight in primary antibody solution: PBT with 1:1000 anti-Brn2 (Cell Signaling), 1:500 anti-Tuj1 (Aves), and 1:1000 Hoechst (Thermo Fisher Scientific). Cells were then washed 2x with PBS and incubated for 1 h in a secondary antibody solution: PBT with 1:1000 donkey anti-rabbit 488 (Thermo Fisher Scientific), and 1:1000 donkey anti-chicken 647 (Jackson Immunoresearch). Samples were then washed 2x with PBS and imaged on a Zeiss LSM900. Images were quantified using custom MATLAB scripts (Figure 6B).

*Excitatory glutamatergic neuron differentiation (Holzbaur lab)*

Cell culture

KOLF2.1J differentiation to NGN2-expressing neurons was performed using an established protocol (Fernandopulle et al. 2018). In short, doxycycline-inducible human NGN2 was stably expressed in wild-type KOLF2.1J iPSCs using a Piggybac vector. Following puromycin selection, iPSCs were differentiated into excitatory glutamatergic neurons with 2 µg/mL doxycycline. Following 72 h of incubation with doxycycline, neurons were dissociated with Accutase (Sigma) and cryo-preserved. On day of use, pre-differentiated neurons were thawed and plated at a density of 300,000 neurons per imaging dish (MatTek) coated with poly-L-ornithine. iPSC-derived neurons were cultured in BrainPhys Neuronal Medium (StemCell) supplemented with 2% B27 (GIBCO), 10 ng/mL BDNF (PeproTech), 10 ng/mL NT-3 (PeproTech), and 1 µg/mL mouse laminin (Corning).

To visualize mitochondria, cells were transfected with a PGK-mEmerald-mito plasmid. To engineer the plasmid, a parent plasmid was obtained from Addgene expressing 4xmito-mEmerald behind a CMV promoter. This insert was put behind a PGK promoter made by PCR using pLenti-PGK-LifeAct-GFP-W (Addgene) as the template, with the vector backbone from a pEGFP vector (Clontech).

iPSC-derived neurons were transfected at DIV18 (72 h prior to fixation) with 4 µL Lipofectamine Stem Transfection Reagent (ThermoFisher) and 1.25 µg plasmid DNA per dish.

Immunostaining

At DIV21, iPSC-derived neurons were fixed with 4% PFA supplemented with 4% sucrose (w/v) for 10 min at 37°C. Neurons were then permeabilized with 0.2% Triton-X in PBS for 15 min at room temperature. After washing three times with PBS, neurons were blocked for 1 h at room temperature with 5% goat serum and 1% BSA in PBS. Neurons were incubated in primary antibodies anti-MAP2 (1:200, EMD Millipore) and anti-LAMP1 AlexaFluor 488-conjugated (1:20, R&D Systems) diluted in blocking solution overnight at 4°C, washed three times with PBS, and then incubated in secondary antibody goat anti-Mouse IgG AlexaFluor 594 (1:2000, Thermo Fisher) diluted in blocking solution for 1 h at room temperature. Nuclear counterstaining was performed with Hoechst dye (ThermoFisher). Fixed iPSC-derived neurons were imaged on a PerkinElmer UltraView Vox Spinning Disk Confocal system with a Nikon Eclipse Ti inverted microscope, a Hamamatsu EMCCD C9100-50 camera controlled by Volocity software, and an Apochromat 100x 1.49 NA oil immersion objective. Z-stacks were collected at 0.15 nm/step (Figure S6B).

*Cortical, medium spiny neuron, and dopaminergic neuron co-cultures (Schüle lab)*

Cell Culture

To generate mixed cultures of cortical, medium spiny (MSN), and dopaminergic neurons, KOLF2.1J iPSCs were first individually neuralized to pre-patterned forebrain, lateral ganglionic eminence, and floor-plate neural progenitor cells (NPCs) by chemical induction (Calatayud et al., n.d.; Shi, Kirwan, and Livesey 2012; Arber et al. 2015; Kriks et al. 2011). KOLF2.1J iPSCs were plated at a concentration of 300,000 cells/cm^2^ in 2 ml of a 1:1 mixture of Stemflex and KSR media (15% knockout serum replacement (KSR), 1x penicillin-streptomycin, 1x GlutaMAX, 1x NEAA, 0.1mM b-mercaptoethanol in Knockout DMEM/F12) containing 1 uM Thiazovivin (THZ) in a Matrigel (1:50 in KnockOut DMEM) -coated well of a 6 well plate. On day 0, media was replaced with 3 ml of (100% KRS media), and cultures were supplemented with small molecules according to the region-specific patterning protocol (see below: (1) forebrain, (2) lateral ganglionic eminence, (3) floorplate NPCs). Starting day 4, KSR Media was gradually shifted to N2B27 media [1x Penicillin Streptomycin, 1x GlutaMAX, 1x NEAA, 0.5x B27 without vitamin A, 0.5x N2 in Neurobasal:DMEM/F12 (1:1)] until day 8 on which 100% N2B27 Media was used. On day 13, splitting (1:1) was performed with Accutase, and cells were replated into a new Matrigel (1:80)-coated well. On day 15, cells were split up onto two Matrigel-coated wells (splitting 1:2). On day 20, three NPC types were detached with Accutase, counted, and combined (1:1:1) in N2B27 media containing 0.5% fetal bovine serum (FBS) and 1 mM THZ.

1. Forebrain NPCs pre-patterned to derive into cortical neurons were generated from iPSCs according to an adapted protocol (Calatayud et al., n.d.) of (Shi, Kirwan, and Livesey 2012). To increase the outcome of PAX6-positive cells, we applied dual SMAD-inhibition by LDN/SB431542 instead of noggin/SB431542 (Surmacz et al. 2012). We further applied wingless/integrated (Wnt) pathway inhibition by exposure to IWP-2 (Moya et al. 2014); see below table).

| **Day of Neuralization** | **Supplements** | **Base Media** | **Media changes** |
| --- | --- | --- | --- |
| Day -1 | THZ (1 µM) | Stemflex (50%) \ KSR (50%) |  |
| Day 0 - 3 | LDN (500 nm), SB431542 (10 µM), IWP-2 (1 µM) | KSR (100%) | Every 2^nd^ day |
| Day 4 - 5 | LDN (500 nm), SB431542 (10 µM) | KSR (66%) \ N2B27 (33%) | Every 2^nd^ day |
| Day 6 - 7 | LDN (500 nm) | KSR (33%) \ N2B27 (66%) | Every day |
| Day 11 - 12 | LDN (500 nm) | N2B27 (100%) | Every day |
| Day 11 - 13 | **THZ (1 µM) 24h before and after splitting | N2B27 (100%) | Every day |
| Day 14 - 20 | THZ (1 µM) | N2B27 (100%) | Every 2^nd^ day |

2. Lateral ganglionic eminence NPCs pre-patterned to derive into GABAergic medium spiny neurons (MSNs) were generated from iPSCs according to a modified protocol (Calatayud et al., n.d.) by (Arber et al. 2015). Exposure to SB431542 was reduced to 6 days (Day 0 to Day 6), while we started to supplement Activin A on day 8 till day 20. We further adapted the LDN concentration to 500 nM and included IWP-2 (see below table).

| **Day of Neuralization** | **Supplements** | **Base Media** | **Media changes** |
| --- | --- | --- | --- |
| Day -1 | THZ (1 µM) | Stemflex (50%) \ KSR (50%) |  |
| Day 0 - 3 | LDN (500 nm), SB431542 (10 µM), IWP-2 (1 µM) | KSR (100%) | Every 2^nd^ day |
| Day 4 - 5 | LDN (500 nm), SB431542 (10 µM) | KSR (66%) \ N2B27 (33%) | Every 2^nd^ day |
| Day 6 - 7 | LDN (500 nm) | KSR (33%) \ N2B27 (66%) | Every day |
| Day 8 - 13 | LDN (500 nm), ActivinA (25ng/ml)  **THZ (1 µM) 24h before and after splitting | N2B27 (100%) | Every day |
| Day 14 - 20 | ActivinA (25ng/ml), THZ (1 µM) | N2B27 (100%) | Every 2^nd^ day |

3. For floor plate-induction of NPCs, we used a modified dual SMAD-inhibition protocol by (Kriks et al. 2011). The concentration of LDN was adapted to 500 nM. We further optimized the time point of FGF8 exposure starting subsequently to SAG/purmorphamine (purm) exposure from day 7 (see below table).

| **Day of Neuralization** | **Supplements** | **Base Media** | **Media changes** |
| --- | --- | --- | --- |
| Day -1 | THZ (1 µM) | Stemflex (50%) \ KSR (50%) |  |
| Day 0 | LDN (500 nm), SB431542 (10 µM) | KSR (100%) |  |
| Day 1 - 2 | LDN (500 nm), SB431542 (10 µM), SAG (2 µM), Purm (2 µM) | KSR (100%) | Every 2^nd^ day |
| Day 3 | LDN (500 nm), SB431542 (10 µM), SAG (2 µM), Purm (2 µM), CHIR (3 µM) | KSR (100%) |  |
| Day 4 - 5 | LDN (500 nm), SB431542 (10 µM), SAG (2 µM), Purm (2 µM), CHIR (3 µM) | KSR (66%) \ N2B27 (33%) | Every 2^nd^ day |
| Day 6 | LDN (500 nm), SAG (2 µM), Purm (2 µM) , CHIR (3 µM) | KSR (33%) \ N2B27 (66%) |  |
| Day 7 | LDN (500 nm), CHIR (3 µM), FGF8 (100ng/ml) | KSR (33%) \ N2B27 (66%) |  |
| Day 8 - 15 | LDN (500 nm), CHIR (3 µM), FGF8 (100ng/ml)  **THZ (1 µM) 24h before and after splitting | N2B27 (100%) | Every 2^nd^ day |
| Day 16 - 20 | THZ (1 µM) | N2B27 (100%) | Every 2^nd^ day |

Codex multiplex imaging

For CODEX® the NPCs had to be plated and differentiated on specific glass coverslips (22x22 mm Electron Microscopy Sciences). The previous day, glass coverslips were placed in 6 wells and coated with poly-D-Lysine (100µg/ml, 2h at RT)/laminin (10µg/ml, 2h at 37 °C) and then pre-seeded with primary human Ara-C inactivated astrocytes (620 cells/cm^2^; Sigma-Aldrich) in astrocyte media. Before plating the NPCs, the astrocyte media was taken out of the well. Then the NPCs were gently pipetted at a concentration of 100,000 cells/cm^2^ in 300 µL of N2B27 media containing THZ (1 µM) onto the glass coverslips only. The NPCs were allowed to attach for 10 min on RT before another 2 ml of N2B27 media containing THZ (1 µM) was carefully added to the well. The following day the media was replaced with 3 ml of terminal differentiation media^2^ (1x penicillin-streptomycin, 1x GlutaMAX, 1x NEAA, 1x B27 without vitamin A, 20 ng/µL BDNF, 10 ng/µL GDNF, 0.5 mM dibutyryl cAMP, 0.2 mM ascorbic acid, 10 µM DAPT in Neurobasal A) containing 0.5% FBS. NPCs were differentiated into neurons for 30 days. For the first two weeks of differentiation half of the media was changed every 3 days, subsequently, media changes were conducted every 7 days. To prevent neurons from detaching, 1 µg/ml of mouse laminin was added to the media every second media change.

Codex multiplex imaging was performed for 10 different antibodies. Fixation, preparation, and staining of the sample, as well as image processing, was conducted as described in (Heinrich et al. 2022; Zafar et al. 2022)

SybrGreen Assay of dopaminergic neurons derived from KOLF2.1J and 0524 control lines

Using a previously established 96-well SYBR Green qPCR expression array (Srinivasaraghavan, Zafar, and Schuele 2022), iPSC-derived dopaminergic neurons from KOLF2.1 and 0524-1 (Khan et al. 2020) were characterized. iPSCs were dissociated with ReLesR and ~2,500,000 cells were pelleted for 5 min by centrifugation (5,000 g at 4°C). The pellet was washed with 1 ml of PBS and again centrifuged at 5,000 g at 4°C. RNA was then extracted using the Purelink™️ RNA Mini Kit. To increase RNA yield, iPSC-derived dopaminergic neurons (~500,000) were dissociated and lysed with Purelink™️ lysis buffer directly in a 6 well. Subsequently, the cell lysate was transferred to the homogenization tube and RNA extraction was conducted according to the Purelink™️ protocol. The RNA concentration was quantified by NanoDrop and 1 μg of RNA was treated with DNase I for 15 min at RT to eliminate contamination with genomic DNA. After inactivating DNase I for 3 min at 65°C for 3 min, HighCapacity cDNA Reverse Transcription kit was used for cDNA reverse transcription. Negative control was generated by adding nuclease-free water instead of RNA. For the reverse transcriptase control RNA was added but not the reverse transcriptase. Each cDNA sample was diluted 1:5 with nuclease-free H_2_O for a final concentration of 10 ng/μL and stored at -20°C.

The day before the SYBR Green assay, both forward and reverse primers of our genes of interest were pre-plated in triplicates at a final concentration of 300 nM each into a 384-well of an optical PCR plate. Additional wells with GAPDH primers were included for the negative and reverse transcriptase control. After plating the primers, the 384 well-plate was centrifuged at 2,000 rpm for 2 min. To dry out the primers, the plate was then kept in a box at RT for at least 12 h. Then the reaction mix (2.5 μL of PowerUp SYBR Green Master Mix, 1.5 μL of nuclease-free water, and 1 μL of 10ng/ml sample cDNA) was pipetted to the wells that were pre-coated with primers, and the plate was sealed with optical adhesive film. Following another centrifugation step at 2,000 rpm for 2 min, the SYBR Green reaction was run using a QuantStudio 6 Flex.

The cycling conditions indicated in the below were used to amplify cDNA.

| **Steps** | **Temperature** | **Duration** | **Cycles** |
| --- | --- | --- | --- |
| UDG activation | 50 °C | 2 min |  |
| Dual-Lock™ Taq DNA polymerase | 95 °C | 2 min |  |
| Denaturation | 95 °C | 15 s | 40 |
| Annealing/ extend | 60 °C | 1 min |  |

The conditions in the below table were used to quantify the melt curve of the PCR product.

| **Step** | **Ramp rate** | **Temperature** | **Time** |
| --- | --- | --- | --- |
| Denaturation | 1.6 °C/s | 95 °C | 15 s |
| Annealing | 1.6 °C/ s | 60 °C | 1 min |
| Dissociation | 0.15 °C/ s | 95 °C | 15 s |

The cycle threshold (Ct) values were used to calculate the fold change (2^-ΔΔCt^) as a measure of relative gene expression. First, the mean Ct value was calculated for each gene. The mean Ct values were used to determine ΔCt by ΔCt = mean Cttarget gene - mean Cthousekeeping gene (GAPDH). Then ΔΔCt was calculated using ΔΔCt = ΔCttreated cell (neuron) – ΔCtuntreated cell (iPSC). Finally, the fold change was obtained by 2^-ΔΔCt^ (Figure S6C).

*Cortical neuron differentiation (Verhage, van der Kant laboratories)*

Cell culture

Cells were differentiated into cortical neurons through forced overexpression of NGN2. iPSCs were infected with high-titer lentiviral particles encoding pTetO-Ngn2-Puro (Addgene) and FUΔGW-rtTa (Addgene) in E8 medium containing 5µM Rock-inhibitor (RI; TetuBio). The next day, the cells were transferred to N2 medium (DMEM/F12 medium supplemented with 200mM Glutamax, 20% Dextrose, 1% N2 supplement B (all Life Tech) and 0.1% Pen/Strep (Invitrogen)) containing 2µg/ml doxycycline hyclate (Sigma Aldrich) as well as the SMAD inhibitors LDN-193189 (100nM; Stemgent) and SB431542 (10µM; Tocris) and the WNT inhibitor XAV939 (2µM; Stemgent) to start NGN2 expression and neuronal differentiation. The next day, 100% of the medium was exchanged for N2 medium containing doxycycline, inhibitors and 2µg/ml puromycin (Merck-Millipore). After another day, the medium was changed to N2 containing doxycycline and 10 µM FUDR (Sigma Aldrich). On the final day, the cells were dissociated and replated on glia islands in neurobasal medium (supplemented with 200mM Glutamax, 20% Dextrose, NEAA, B27, 0.1% P/S, 0.5% Fetal bovine serum; all Life Tech) containing doxycycline and growth factors BDNF, CNTF and GDNF (all 10ng/ml; Stem Cell Tech). 2k neurons were plated on 18mm coverslips covered in glia islands. To create the islands, rat glia were plated on etched coverslips stamped with microdots of 0.1mg/ml poly-D-lysine (Sigma Aldrich), 0.7mg/ml rat tail collagen (BD Biosciences) and 10mM acetic acid as described previously (Meijer et al., 2019). Neurons were kept at 37°C and 5% CO_2_. 50% of medium was refreshed once a week. Experiments were performed on days 42 to 46 after the start of induction as in (Meijer et al. 2019).

Bioni010-C-13 iPSCs were acquired from the European Bank for Induced Pluripotent Stem Cells (EBiSC,<https://ebisc.org/>). This iPSC line was CRISPR-engineered to carry the NGN2 induction cassette in the AAVS1 safe-harbor locus (Schmid et al., 2021). Therefore, the line was induced only through the addition of doxycycline and no addition of antibiotics was necessary. In all other aspects it was treated similarly to the KOLF2.1J cell line.

Calcium imaging

6-week old KOLF2.1J neurons were incubated for 10 min with 2 µM Fluo-5F-AM (Molecular Probes; stock in DMSO) at 37°C. Coverslips were transferred to an imaging chamber and perfused with Tyrode’s solution (2 mM CaCL2, 2.5 mM KCl, 119 mM NaCl, 2 mM MgCl2, 30 mM glucose, 25 mM HEPES; ph 7.4). Imaging was acquired on a custom-built microscope (AxioObserver.Z1, Zeiss) with 40x oil objective (NA1.3). Neurons were imaged for 30 s as baseline and then stimulated with 16 trains of 50 action potentials at 50 Hz with a 0.5 s interval. Electrode field stimulation was applied using a stimulus generator (A-385, World Precision Instruments) controlled by a Master-8 (AMPI) to deliver 1 ms pulses of 30 mA. Experiments were performed at room temperature (20-24C) For calcium influx quantification, 20 neurite-located ROIs (6 x 6 pixels) and background ROIs were measured per neuron in ImageJ (Figure S6D).

Electrophysiology methods

Autaptic neurons were subjected to whole-cell voltage-clamp recordings (Vm = -70 mV). Experiments were performed at room temperature with borosilicate glass pipettes (Science products GmbH, 2.5-4.5 MOhm) filled with (in mM): 136 KCl, 17.8 HEPES, 1 EGTA, 0.6 MgCl2*6H2O, 4 ATP-Mg, 0.3 GTP-Na, 12 phosphocreatine dipotassium salt and 50 U/ml phosphocreatine kinase (pH = 7.3, ~300 mOsmol). External solution (aCSF) contained the following (in mM): 10 HEPES, 10 Glucose, 140 NaCl, 2.4 KCl, 4 MgCl2 and 2 CaCl2 (pH = 7.30, ~300 mOsmol). aCSF was made from stock with HEPES and glucose added freshly, filter-sterilized and stored at 4°C until use. Patch-clamp recordings were performed with a MultiClamp 700B amplifier and Digidata 1550B or an Axopatch 200B amplifier and Digidata 1440A, controlled by Clampex 10.6 software (Molecular Devices). For gap-free recordings of spontaneous miniature EPSCs, the sampling rate was set to 20 kHz and low-pass Bessel filter was set to 5-6 kHz. For episodic stimulations, the sampling rate was set to 10 kHz and low-pass Bessel filter to 2 kHz. Resistance was compensated by 70-80% (bandwidth 7.52 Hz). Action potentials were elicited by a 1 ms depolarization to 30 mV. Recordings were excluded if series resistance was higher than 15 MOhm, if the leak current was larger than 300 pA and if the evoked EPSC was too asynchronous as assessed by a custom-made MATLAB script. Offline analysis was performed with MATLAB R2019a (Mathworks) using custom-written software routines (viewEPSC, downloaded from vhuson user on Github on 6^th^ Jan 2020). Data were tested for normality and homoscedasticity and significance was tested using the Wilcoxon rank sum test with p values below 0.05 considered as significant (Figures 6C; S6E).

*Cortical neuron differentiation (Cohen lab)*

Generation of PB-TO-hNGN2-hiPSCs

An hiPSCs-PB-TO-hNGN2 stable cell line was generated as previously described (Fernandopulle et al. 2018)**.** Briefly, KOLF2.1J wildtype cells were transfected with piggyBac plasmid carrying rrTA and Ngn2-Puro cassette (plasmids gifted from Michael Ward). Transfected cells were selected for stable integration using puromycin treatment (0.5 µg/mL) and propagated as a non-clonal pool. The expression of stem marker genes was validated by immunofluorescence and western blots for Nanog (Abcam), Sox2 (Abcam), Oct4 (Abcam), and SSEA4 (Thermo Fisher).

Differentiation into cortical neurons

hiPSCs-PB-TO- hNGN2 were differentiated into cortical neurons as previously described (Fernandopulle et al. 2018) with slight modification. In brief, hiPSCs were seeded in colonies (10-20 cells) at high confluency (50-60%) on Vitronectin coated plates. One day after seeding, the medium was changed to Induction media: DMEM/F12 with HEPES (Gibco); N2 supplement 100X (Gibco); non-essential amino acids 100X (Gibco), and supplemented with Doxycycline at a final concentration of 1µM (Sigma). The medium was changed every day. After 2.5 days, the pre-induced hiPSCs, were passaged in Accutase (StemPro™ Accutase™ Cell Dissociation Reagent, Gibco) and seeded single cell (300,000) on Poly-L-Ornithine (Sigma; 10x stock: 50 mg in 50 ml Borate Buffer) and laminin 10 µg/ml (Gibco) coated plates (Thermo Scientific™ Nunc™ Lab-Tek™ II Chambered Coverglass, 2-well). The day of seeding, the medium was changed to Cortical Neuron Culture Medium (CM): BrainPhys neuronal medium without Phenol-Red (STEMCELL Technologies); B27 supplement, 50X (Gibco); BDNF (10 µg/ml) in PBS containing 0.1% IgG and protease-free BSA (PeproTech); NT-3 (10 µg/ml) in PBS containing 0.1% IgG and protease-free BSA (PeproTech); laminin final con. 1 µg/ml (Gibco). CM medium was initially supplemented with ROCK inhibitor (RevitaCell supplement 100X, Gibco). The day after seeding, the medium was replaced with CM medium without ROCK inhibitor. The i^3^Neurons were kept for 27 days prior to imaging, with half of the medium replaced at least once per week with freshly prepared CM. Cells were immunostained for different neuronal markers: rabbit anti-MAP2 (Abcam); rabbit anti-NeuN (CellSignalling); rabbit anti-βIII-Tubulin (Abcam); rabbit anti-PSD95 (Abcam); mouse anti-VGLUT1 (Sigma); and rabbit anti-TBR1 (Abcam).

Transfection of i^3^NeuronsAfter 29 days of induction (day 27), the i^3^Neurons were transfected as previously described (Dalby et al. 2004) with some modifications. The transfection mix was prepared in Neurobasal (Gibco) with 6 µl of Lipofectamine2000 (Invitrogen), and 6 µg of total DNA divided as follows: 2 µg of pEIF1a::Transposase (gifted by Dr. M. Ward), and 4 µg of pEIF1a::Cox8(1-26)::eGFP (Twist Technologies). The medium of i^3^Neurons was changed to Neurobasal (without Glutamine) and pre-incubated at least 30 min. before adding the transfection mix. Next, half the medium was removed, and the transfection mix was added dropwise. After 2 hrs of incubation, the medium was completely replaced with CM.

Imaging

24 hrs after transfection the neurons were tested for vitality by NeuroFluo (final concentration 0.20µM, STEMCELL Technologies) and NuclBlue (Invitrogen) staining. Images were acquired with a Zeiss 800 Laser Scanning Confocal Microscope with a 63x Objective (Plan-Apochromat 63x/1.40 Oil DIC M27). The NeuroFluo/NuclBlue images and the brightfield images of mitochondria/NuclBlue were taken with a zoom of 0.5x, while the time lapse images were taken with a zoom of 2.2x, every 1.96 seconds for 100 cycles. The cropped timelapse corresponds to cycle 53 and is referred to as the starting time (t=0s) for the fission event observed (Figure S6F).

*Cortical and microglia differentiations (Paquet lab)*

Cortical neuron differentiation

iPSC-derived cortical neurons were generated as previously described (Paquet et al., 2016) with minor modifications. Briefly, 1 million iPSCs were plated on day 0 on 12-well tissue culture plates coated with Geltrex in neural induction medium (NI, 0.5x Neurobasal, 0.5x DMEM/F12, 0.1 mg/ml Pen/Strep, 0.5x B27, 0.5x N2, 2 mM Glutamax, 0.1 mM NEAA, 5 µg/ml Insulin, 0.1 mM 2-Mercaptho-Ethanol, 10 µM SB431542, 0.25 µM LDN-193189) and maintained for 8 days. On day 8 cells were dissociated using Accutase (Life Technologies) and resuspended in NI medium at 30 million cells/mL. Cells were plated on dried poly-L-ornithine and laminin-coated (POL; Sigma-Aldrich, Life Technologies) 6-well plates in 200-300 µL spots. Cells were left to adhere for ~45 min and NI medium was added. On day 10 NI was replaced with a neural maintenance medium (NM, which is NI without SB431542 and LDN-193189). Upon the appearance of neural rosettes, 20 ng/mL FGF2 was added for 2 days. When neurons started to form (~day 21), rosettes were isolated manually after treatment with STEMdiff Neural Rosette Selection Reagent (STEMCELL Technologies) for 1 h. Rosettes were washed and plated on POL 6-well plates and grown for 7 days. Cortical neurons were harvested with Accutase and plated on POL-coated coverslips at 400k per 24-well well and maintained in Neurobasal medium (NB, 1x Neurobasal, 1x B27, 2 mM Glutamax, 0.1 mg/mL Pen/Strep, all Life Technologies) until analysis was performed (Figure 6D).

Microglia differentiation

KOLF2.1J iPSCs were differentiated as described previously (McQuade et al. 2018; Reifschneider et al. 2021; Figure 6I).

Immunostaining and quantifications

Cells were fixed by 4% paraformaldehyde (PFA) for 20 min at RT, washed twice for 5 min with PBST (PBS, 0.1% Triton X-100), incubated in blocking solution (PBST, 3% donkey serum, 0,02% NaN3) for 1 hr, and incubated with primary antibodies in blocking solution (neurons: Rabbit anti-Tuj1, 1:500, Covance; chicken anti-MAP2, 1:2000, Abcam; rat anti-CTIP2, 1:200, Abcam. Microglia: Rabbit anti-PU.1, 1:300, Cell Signal; goat anti-TREM2, 1:400, Bio-Techne; rabbit anti-Iba1, 1:500, Invitrogen) at 4˚C overnight. On the next day, cells were washed 4 times for 10min in PBST, and incubated with Alexa488, Alexa555 or Alexa647-conjugated secondary antibodies (Invitrogen) and 4’,6-diamidino-2-phenylindole (DAPI) for 1 h at RT. Finally, cells were washed 4 times for 10min in PBST and mounted on slides for imaging. Images were visualized in a Zeiss Observer widefield microscope. Quantification of purity was performed manually for all microglial stainings, as well as MAP2 and TuJ1 for neurons, and using a custom macro in Fiji/ImageJ (National Institutes of Health) for the remaining stainings and DAPI. Briefly, the inbuild ImageJ Macro environment was used to write a script that would batch process triple stained ICC images. The script loaded the images from a certain directory, split the image into individual channels and then counted the number of positive cells of each stain after thresholding the image. Size and circularity exclusions were applied to restrict the counting to individual cells rather than clumps. To count DAPI positive cells, the inbuild threshold ‘Yen’ was used and particles with a size between 30-250µm and a circularity between 0.1-1 were counted. Cells were defined by the same size and circularity, but the inbuild threshold ‘Li’ was used to differentiate positive signal from background. Quantification of yield was performed by manually counting cells using a Neubauer chamber at the same time when neurons or hematopoietic microglia precursors are usually frozen for storage.

*Cortical neurons (Wray lab)*

Cell Culture

KOLF2.1J iPSCs were differentiated in parallel to previously described control iPSCs (Arber et al. 2020) into cortical glutamatergic neurons according to the protocol described in (Shi, Kirwan, and Livesey 2012)**.** This protocol generates glutamatergic neurons representative of the six cortical layers in a 100-day period. Briefly, iPSC cells were grown in StemFlex media (Thermofisher) on a Geltrex (Thermofisher)-coated plate in a humidified 5% CO_2_. For differentiation, cells were plated at 100% confluency (day 0 in culture) on geltrex-coated plates and grown in a neuronal induced medium which was replaced daily for ~12 days. Neuronal induction media was N2B27 media containing 10 μM SB431542 (Tocris) and 1 μM dorsomorphin (Tocris). N2B27 media consisted of a 1:1 mixture of Dulbecco's modified eagle medium F12 (DMEM-F12) and Neurobasal supplemented with 0.5× N2, 0.5× B27, 0.5× non-essential amino acids, 1 mM L-glutamine, 25U pen/strep, 10 μM β-mercaptoethanol and 25U insulin. Following 12 days of neuronal induction media, a uniform neuroepithelial layer could be observed. This was passaged using dispase (Life Technologies) and replated in large clumps onto laminin coated wells at a ratio of 1:3 (Sigma) to allow the formation of neuronal rosettes. These were fed every 3 days with N2B27 media and passged with dispase onto fresh laminin-coated wells as required. Once substantial neurogenesis was observed (25-35 days in culture), cells were dissociated to a single-cell suspension using Accutase (Innovative Cell Technologies) and plated onto fresh laminin coated wells. Neurons were plated for the final time at 35 days in culture at a density of 50,000 cells per cm^2^. Neurons were fed every 72h with N2/B27 media until 100 days in vitro, when corticogenesis is complete, and characterized by immunofluorescence for neuronal markers.

Immunostaining

At 100 DIV, cells were fixed in 4 % (wt/vol) paraformaldehyde (PFA) for 10 min before permeabilization with 0.3 % (vol/vol) Triton-X-100 in PBS for 10 min and blocking with 5 % BSA in PBS for 30 min at RT. After blocking, cells were incubated with rat anti-Ctip2 (1:300; Abcam) and mouse anti-Tuj1 (1:1000; Promega) primary antibodies diluted in the blocking solution overnight at 4 °C in a humid chamber. The following day, primary antibodies were removed, and cells were washed with PBS three times for 5 min. After the last wash, the cells were incubated with Alexa Fluor 488 anti-rat and Alexa Fluor 647 anti-mouse (1:1000) secondary antibodies diluted in a blocking buffer for 1h at RT protected from light. Secondary antibodies were then removed, and cells were washed twice with PBS and then counterstained with diamidino-2- phenylindole (DAPI) (Thermofisher) at 0.1 µg/ml for 10min at RT and washed three times with PBS. Cells were mounted on microscope glass slides (Thermofisher) with ProLong™ Diamond Antifade Mountant (Thermofisher). For high content imaging, plates were stored in PBS. Stained cells were imaged using a ZEISS LSM 800 microscope and analysed using Fiji software or for quantification, cells were imaged on the Opera Phenix High Content Screening System (Figure 6E).

*Skeletal myocyte differentiation (Raman lab)*

Skeletal Myocyte differentiation

Skeletal myocyte differentiation (Figure S6G) was performed according to a previously published study (Chal et al. 2016).

Immunostaining

iPSCs, premyogenic progenitors, and myoblasts were plated on 12-mm coverslips in 24-well plates. Myocytes were differentiated on the coverslips for ten days. Cells were washed with PBS and fixed using 4% paraformaldehyde for 15 min at room temperature. Cells were blocked and permeabilized in PBS containing 5% serum and 0.2% Triton X-100 for 1 h at room temperature. The coverslips were then incubated in primary antibodies rabbit anti-Nanog, mouse anti-Pax3, rabbit anti-MyoD, and mouse anti-myogenin diluted in blocking buffer in a humidified chamber overnight at 4C. Post primary antibody incubation, the coverslips were washed three times with PBS followed by 1 h incubation with appropriate Alexa Fluor-conjugated secondary antibodies in the dark in blocking buffer. Cells were washed with PBS, and nuclei were stained with Hoechst dye (1:5000) and mounted onto slides. The images were collected by using a Nikon A1R with a 60× Plan Apo 1.4-numerical-aperture (NA) objective lens or Zeiss LSM 800 with a 40x/1.3 NA Plan Apochromat oil immersion. Images were analyzed by using FIJI (<https://imagej.net/Fiji>).

*Cortical neuron differentiation (Parish lab)*

Cell Culture

The KOLF2.1J IPSC line was differentiated into cortical neurons using a dual SMAD neural induction protocol, as previously described (Gantner et al. 2021). Briefly, KOLF2-1 iPSCs were seeded onto Laminin-521-coated (5 µg/mL) plates at a density of 0.3 x 10^6^ cells/cm^2^ in mTeSRPlus medium (StemCell Technologies) with 10 µM Rock inhibitor Y-27632 (Tocris Bioscience). 24 hrs later, cells were switched to dual-SMAD inhibition for 11 days in ‘cortex medium’ comprised of 1:1 DMEM/F12 and Neurobasal with 0.5x B27, 0.5x N2, 0.5x ITSA, 1x GlutaMAX, 0.5x Penicillin Streptomycin and 50uM 2-Mercaptoethanol (Life Technologies) and supplemented with 100 nM LDN193189 (Stemgent) and 10 mM SB431542 (R&D Systems). On Day 11 (D11), progenitors were passaged in Accutase (Innovative Cell Technologies) for 5 min at 37°C and transferred to ‘cortex medium’ containing 20 ng/mL fibroblast growth factor 2 (FGF2; R&D Systems) for 8 days. On D19, cultures were again passaged and replated in cortex medium. On D32, cells were passaged and seeded on poly-L-ornithine (0.005% v/v, Sigma-Aldrich, USA) and Laminin-521-coated plates. One week following final plating, neurons were switched to ‘maturation medium’ consisting of 1:1 DMEM/F12 and Neurobasal, 1x B27, 1x N2, 1x ITSA, 1x NEAA, 1x GlutaMAX and 0.5x Penicillin Streptomycin, supplemented with brain-derived neurotrophic factor (BDNF, 40ng/ml; R&D Systems), glial cell-line derived neurotrophic factor (GDNF, 40ng/ml), N^6^,2′-O-Dibutyryladenosine 3′,5′-cyclic monophosphate (dcAMP, 0.05mM; Tocris Bioscience), ascorbic acid (200nM; Sigma-Aldrich) and laminin (1µg/ml; Sigma-Aldrich).

Immunostaining

Immunostaining of cultures and brain sections were performed as previously described (Gantner et al. 2021). In brief, cultures were fixed in 4% (w/v) paraformaldehyde for 10 min, washed in PBS and incubated overnight in PBS-Azide (0.02% w/v) solution containing 10% (w/v) normal donkey serum (NDS), 0.3% Triton-X and primary antibodies rabbit anti-TBR1 (1:1000, Abcam), rat anti-CTIP2 (1:500 Abcam), and goat anti-BRN2 (1:200, Santa Cruz). The following day, cells were washed, blocked with 10% NDS and incubated in secondary antibodies (AlexaFluor-488, -555 or -647; Jackson ImmunoResearch) for 1.5 hrs. Nuclei were counterstained with DAPI. Images were captured on a Zeiss Axio ObserverZ.1 inverted epifluorescence microscope at 20x and the total number of DAPI, TBR1, CTIP2, and BRN2 immunolabeled cells quantified (Figure 6F).

*Motor neuron differentiation (Conklin lab)*

Differentiation

KOLF2.1J iPSCs were differentiated to motor neurons by inducible expression of 3 transcription factors in the hNIL transgenic system (Fernandopulle et al. 2018). Briefly, 350k KOLF2.1J iPSCs were electroporated (Lonza 4D nucleofector, setting DS-138; Amaxa P3 Primary Cell 96-well Nucleofector kit) with plasmids containing the i3 LMN hNIL inducible construct (pUCM-CLYBL-hNIL, Addgene) and TALENs targeting the CLYBL safe harbor locus (pZT-C13-R1, Addgene; pZT-C13-L1, Addgene). Transfected cells were plated at clonal density and selected with 100µg/mL geneticin starting day 3 after transfection for 10 days, at which point all surviving colonies were observed to express mCherry. Surviving iPSC colonies were pooled and expanded for 5 passages before beginning the differentiation. Differentiation of motor neurons was conducted as described in (2). For day 3 replating, 20k cells were passaged into 96-well plastic cell culture dishes. On day 10, cells were fixed with 200 µL 4% PFA added directly to cell culture media (final concentration 2%) and fixed at room temperature for 20 min.

Immunostaining

Cells were permeabilized with 0.1% TritonX in PBS (PBST) for 10 min and blocked with 5% BSA in PBST for 1 h. Cells were incubated with primary antibodies in PBST (rabbit Anti-β-Tubulin III, Sigma, 1:500 or mouse Anti-HB9, DSHB, 1:200) at room temperature for 1 h, washed 3x with PBST for 5min, secondary antibodies (Alexa Fluor-594 Goat anti-Rabbit IgG, Invitrogen, 1:500 or Alexa Fluor-488 Goat anti-Mouse IgG, Abcam, 1:500) in PBST at room temperature for 45min, then washed 3x with PBST for 5min with DAPI in the first wash. Cells were imaged on a Keyence BZ-X710 (Figure S6H).

*Motor neuron differentiation (Zhang lab)*

Cell Culture

Motor neuron differentiation was performed as described previously (Coyne et al. 2020)

Immunostaining

On day 18, cells were fixed in 4% paraformaldehyde for 20 min, incubated in PBX (PBS and 0.1% Triton X-100) for 10 min. Cells were incubated in mouse anti-SMI32 (Biolegend) in PBST (PBS and 0.1% Tween-20) with 3% normal donkey serum at 4 °C overnight. Cells were then washed three times in PBST for a total period of 1 h and subsequently incubated in the secondary antibody solution, PBST with 3% normal donkey serum and 1:1000 Donkey anti-mouse Alexa 488, for 3 hrs at room temperature. Cells were washed three times in PBST for a total period of 1 h and then mounted in Prolong Gold with DAPI. Three images were selected per line for quantification (Figure S6I).

*Motor neuron differentiation (Cleveland lab)*

iPSCs were differentiated to motor neurons as described previously (Martinez et al. 2016). iPSCs were grown in Matrigel-coated plates until 70-90% confluency in mTeSR-Plus medium (~2 days from passaging). Neural induction (day 1) was done by medium change to N2B27 medium (DMEM/F12 + GlutaMAX supplemented with 1% PenStrep, 1:200 N2 supplement, 1:100 B27 Supplement, 150μM Ascorbic Acid) supplemented with 10μM SB431542, 1μM Dorsomorphin and 3μM CHIR99021. Daily media changes were done until day 7, when cells were passaged at 1:6 and medium was changed to N2B27 supplemented with 10μM SB431542, 1μM Dorsomorphin, 200nM Smoothened Agonist (SAG) and 1.5μM Retinoic Acid (RA). Daily medium change were done daily with doubling volumes to adjust for cell density. At day 18, cells were replated using Accutase at a density of 7.5x10^6^ cells in Matrigel-coated 10cm dishes, and fed with N2B27 medium supplemented with 200nM Smoothened Agonist (SAG) and 1.5μM Retinoic Acid (RA). On day 22, cells were passaged onto their final plates or microfluidic chambers coated with 10μg/ml poly-D-lysine, 10μg/ml poly-L-ornithine, and subsequently 20μg/ml laminin, and fed with N2B27 medium supplemented with growth factors (2ng/ml BDNF, CTNF and GDNF), and 2μM DAPT. From day 25 onward, cells were fed every 2-3 days with N2B27 medium containing growth factors.

Immunostaining

Motor Neurons were fixed with 4% PFA for 10 min at room temperature. Permeabilization and blocking were done with 0.01% (w/v) Tween/PBS containing 1.5% BSA for 1 h. Cells were incubated overnight at 4^o^C in primary antibodies mouse anti-islet1/2 (1:1000, DSHB) and rabbit anti-NF-H (1:500, Millipore) were diluted in 0.01% Triton X-100/1x PBS. Cells were then washed 3 times with 1x PBS, and incubated with secondary antibodies (1:500) for 30 min-1h. After two subsequent washes with 1x PBS, cells were incubated with 1:5000 DAPI for 5 min, and stored in 1x PBS until imaging.

Live imaging using SiR-Tubulin

Motor neurons are incubated with 250nM SiR-Tubulin (Cytoskeleton, Inc.) diluted in culture medium overnight, and imaged the next day. (Figure 6G).

*Macrophage differentiation (Bosco lab)*

Cell culture

iPSCs were maintained and differentiated as described previously with minor modifications (Gutbier et al., 2020). In summary, iPSCs were maintained in mTeSR1 Plus media (Stem Cell Technology) in plates coated with 10µg/mL of Cell Adhere Laminin 521 (Stem Cell Technology) diluted in Dulbecco's phosphate-buffered saline (DPBS) containing calcium and magnesium (Gibco). Embryoid body (EB) generation was performed by dissociating the iPSCs colonies and plating 10^4^ cells/well into low adherence 96-well plates (Corning) using mTeSR1 Plus media containing 50 ng/mL human bone morphogenetic protein 4 (BMP-4, Fisher Scientific), 50 ng/mL human vascular endothelial growth factor (VEGF, PeproTech), 20 ng/mL human stem cell factor (SCF, PeproTech), and 10uM ROCK inhibitor (Fisher Scientific). Plated cells were centrifuged at 800 rpm for 3 min and half-medium change was performed on day 2. For myeloid maturation, EBs were gently dislodged from the wells and plated on 6 well plates coated with growth factor reduced matrigel (Corning) diluted in KnockOut DMEM/F-12 (Gibco). EBs were maintained in X-VIVO 15 media (Lonza) supplemented with 1% penicillin/streptomycin (Gibco), 1X GlutaMAX (Gibco), 55 µM 2-mercaptoethanol (Gibco), 100 ng/mL human macrophage colony-stimulating factor (M-CSF, PeproTech), and 25 ng/mL human interleukin- 3 (IL-3, PeproTech). Macrophage progenitors were collected weekly during medium change and terminally differentiated into unpolarized iPSC-derived macrophages by culture for seven days in X-VIVO 15 supplemented with 1% penicillin/streptomycin, 1X GlutaMAX, and 100 ng/mL M-CSF at a density of 100,000 cells per cm^2^. Half-medium change was performed every 3 days.

Immunostaining

iPSC-derived macrophages plated on coverslips were fixed with 4% paraformaldehyde (Fisher Scientific) and processed for immunofluorescence analysis as detailed previously (Schmidt et al., 2021). Cells were incubated in a rabbit anti-IBA1 (1:1000, Wako Chemicals USA) primary antibody.

Immunofluorescence images were collected with a Leica DMI 6000B inverted fluorescent microscope using a 40X air objective and Leica DFC365 FX camera with AF6000 Leica Software v3.1.0 (Leica Microsystems). Stacked images (z=0.2µm) were maximum projected. Brightness and contrast were equally adjusted post-acquisition to improve visualization of fluorescent signals. Brightfield images were acquired on live cells using an EVOS ci inverted microscope (AMG) with 4x and 10x objectives. (Figure S6J).

*Astrocyte differentiation (Kampmann lab)*

NPC cell culture

KOLF2.1J-derived astrocyte differentiation was conducted according to a previous study (Tcw et al. 2017). KOLF2.1J and WTC11 iPSCs were dissociated with accutase and seeded into Aggrewell 800 plates (Stemcell Technologies) following manufacturer's specifications at ~2 million cells/well in DMEM/F12 Media (Gibco) with 1X B27 Supplement minus vit A (Gibco) and 1X N2 (Gibco) in 10 nM Rock Inhibitor (Tocris) to generate embryoid bodies (EBs) (Day 1). The next day, the media was exchanged with the same media with Dual SMADi 0.1 μM LDN193189 (Tocris) and 10 μM SB431542 (Tocris), but without a rock inhibitor. Media containing Dual SMADi was exchanged every other day. On day 7, the cultures were plated onto matrigel coated plates. Adherent cultures were then fed with the same media every other day for the next week. On day 14, neural rosettes were released and replated onto matrigel coated plates in NPC media (1X N2, 1X B27, 20 ng/mL FGF2 in 1% BSA in DMEM/F12) and expanded. On day 21, NPCs were stained with PE mouse anti-CD133/1 (Miltenyi Biotec) and PerCP-Cy5.5 mouse anti-CD27 (BD Pharmingen) and CD133+/CD271- populations were sorted with BD FACSAria Fusion. Mouse IgG1κ was used for isotype control (Miltenyi Biotec).

Astrocyte cell culture

CD133+/CD271- NPCs were expanded in NPC media and then plated on matrigel coated plates with Astrocyte Media (ScienCell Research Laboratories). Astrocytes were cultured in astrocyte media over a period of 2 weeks for astrocyte differentiation on 96 well plates (Corning).

Immunostaining

Astrocytes were fixed with 4% paraformaldehyde in PBS for 10 min at room temperature. They were then washed 3 times with DPBS + 0.3% Triton-X 100 (PBSTx) and blocked with 10% Normal Goat Serum + 1% BSA in PBSTx for 1 h at room temperature. Astrocytes were incubated in the following primary antibodies: Mouse anti-S100β (1:500, Sigma Aldrich) and rabbit anti-NFIA (1:500, Sigma) at 4°C overnight. Following washes in PBSTx, astrocytes were incubated in secondary antibodies goat anti-mouse Alexa Fluor 488 (1:2000, ThermoFisher Scientific) and goat anti-rabbit 568 (1:2000, ThermoFisher Scientific) for 45 min at room temperature. Astrocytes were then washed again in PBSTx, stained with Hoechst, and then washed and stored for imaging in PBS. Plates were imaged on the IN Cell Analyzer 6000 (GE Healthcare), using a 60X 0.7 NA objective, 2x2 binning, with 9 fields per well (Figure 6H).

*Microglia differentiation (Kronenberg-Versteeg lab)*

Mouse organotypic hippocampal brain slice cultures

Hippocampal slice cultures were prepared from pups at postnatal day 4–6 (P4–6) according to previously published protocols (Novotny R et al, J Neurosci, 2016; Mayer D et al, J Virol, 2005). In brief, after decapitation, brains of pups were removed, hippocampi dissected and cut perpendicular to the longitudinal axis into 350 μm sections with a tissue chopper. Hippocampal sections were kept in ice-cold preparation buffer (minimum essential medium (MEM, Gibco) supplemented with 2 mM GlutaMAX™ (Gibco) at pH 7.3) until placed onto a humidified porous polyethylene (PTFE) membrane insert (Merck Millipore) in a 6-well plate with 1.2 ml culture medium (20% heat-inactivated horse serum (Gibco) in 1x MEM complemented with GlutaMax™ (1 mM), ascorbic acid (0.00125%, Sigma-Aldrich), insulin (1 μg/ml, Thermo Fisher), CaCl_2_ (1 mM, Sigma-Aldrich), MgSO_4_ (2 mM, Sigma-Aldrich) and D-glucose (13 mM, Roth) adjusted to pH 7.3) per well. HSCs were kept at 37 °C in humidified CO_2_-enriched atmosphere with media changed three times per week.

To allow engraftment of iMacs into OHSCs, endogenous murine microglia were depleted with a mouse-specific ɑ-CSF1R antibody (BioLegend, 5 mg/ml). On day 3-5 post preparation iPSC-derived microglia precursor cells were harvested and drop-grafted onto OHSCs in 1 µl (10,000 cells/µl) of medium per OHSC. Subsequently, iMacs were allowed to integrate into OHSCs and slices were fixed after 2-4 weeks in culture with 4% paraformaldehyde (PFA) in PBS at pH 7.4 for 2h. After fixation, slice cultures were rinsed 3 times with 0.1M PBS for 10 min and stored at 4 °C until further processing.

Human iPSC-derived microglia-like cell differentiation using a 2D protocol

hiPSC were differentiated as previously described (Takata et al, Immunity, 2017). In brief, 10,000 cells/cm^2^ were plated onto Geltrex (Thermo Fisher) coated 6 well plates. Starting the day after splitting, iPSC were specified to mesoderm with BMP-4 (Miltenyi), VEGF (Miltenyi) and CHIR99021 (Miltenyi) during the first four days of differentiation. Hemangioblast formation was induced by adding FGF-2 (Miltenyi) and maintained with VEGF and FGF-2 from days 4-6. Primitive hematopoiesis was promoted from day 6-10 through Wnt inhibition (Dkk1) and hematopoietic cells were matured from day 12-16 by continued incubation with SCF (R&D Systems), FGF-2, IL-3 (Miltenyi) and IL-6 (Miltenyi).

The first 8 days of the differentiation protocol the cells were cultured in a hypoxic environment (5% CO_2_ and 5% O_2_) before being moved to a normoxic incubator (5% CO_2_) after Differentiation Day 8. Cells were cultured in Stempro Medium (Thermo Fisher) with the addition of the following cytokines: D0 (5 ng/ml BMP4, 50 ng/ml VEGF, and 2 mM CHIR99021), Day 2 (5 ng/ml BMP4, 50 ng/ml VEGF and 5 ng/ml FGF2), Day 4 (15ng/ml VEGF and 5 ng/ml FGF2), Differentiation Day 6 to 10 (10 ng/ml VEGF, 10 ng/ml FGF2, 50 ng/ml SCF, 30 ng/ml DKK-1 (Miltenyi), 10 ng/ml IL-6, and 20 ng/ml IL-3), Differentiation Day 12 and 14 (10 ng/ml FGF2, 50 ng/ml SCF, 10 ng/ml IL-6, and 20 ng/ml IL-3). From day 16 the cells were fed with Stempro supplemented with 50 ng/ml CSF-1 (Miltenyi) with full media changes every 3 days. At D24, .

Immunostaining

Organotypic brain slice cultures were blocked with 5% normal donkey serum (2h) and 0.3% Triton X-100 (Sigma-Aldrich) in PBS. For microglial detection goat anti-Iba1 (1:250, Novus Biologicals) and mouse anti-STEM101 (1:250, TaKaRa) were used in 2% NDS, 0.3% Triton X-100 at 4 °C overnight. Following Alexa-fluorophore-conjugated secondary antibodies were applied in a concentration of 1:250 for 2h at RT: donkey-anti-goat Alexa-647, donkey-anti-mouse Alexa-488, (all Jackson ImmunoResearch).

Slices were analyzed with a Zeiss LSM 880 NLO microscope equipped with a 20x water-immersion objective (W Plan-Apochromat x20/1.0, Carl Zeiss, Jena) (Figure S6K).

*Dopaminergic neuron differentiation (Verstreken lab)*

On day -1, 400k KOLF2-1J or SFC065 (kindly provided by the laboratory of C. Klein) hiPSC/cm2 were seeded in Matrigel-coated 6-well plate wells in StemFlex medium supplemented with 10 µM RI. On day 0, medium was switched to to Neurobasal/0.5xB27 supplement without vitamin A, 0.5xN2, GlutaMAX, Penstrep, non-essential amino acids supplemented with LDN193189 (500 nM, Sigma), SB431542 (10 μM, Tocris), SHH-C24II (200 ng/ml, Miltenyi Biotec), Purmorphamine (0.7 μM, Sigma) and 0.7 µM CHIR99021 (Stemcell Technologies). CHIR99021 concentration was raised to 3 µM from day 4 to day 11, moment at which it was withdrawn from the medium. LDN193189 (500 nM, Sigma), SB431542 (10 μM, Tocris), SHH-C24II (200 ng/ml, Miltenyi Biotec), Purmorphamine (0.7 μM, Sigma) were withdrawn from the medium at day 7 and FGF8b (100 ng/mL, R&D systems) was introduced from day 9 until day 16. At day 11, medium was shifted to Neurobasal-A medium (Life Technologies; 10888-022) supplemented with 1x B27 without Vit. A (Life Technologies; 12587010), 1x GlutaMAX (Life Technologies; 35050-038), 1x PenStrep (Life Technologies; 15140-122), 10 ng/mL BDNF (R&D systems; 248-BDB-050/CF), 10 ng/mL GDNF (R&D systems; 212-GD-010), 200 µM ascorbic acid, 0.5 mM dbcAMP (Sigma-Aldrich), 10 µM DAPT (Tocris; 2634). On day 17, ventral midbrain neural progenitors were cryopreserved.

Immunostaining

Cells were fixed at day 17 of differentiation for 15 min in 4% formaldehyde. Cells were blocked for 1h at room temperature with 3% normal goat serum + 0.3% Triton X-100 (Sigma) in DPBS supplemented with Ca^2+^ and Mg^2+^ (Life Technologies). Cells were incubated overnight at 4°C in primary antibodies mouse anti-EN1 (1:200, DSHB), rabbit anti-LMX1A/B (1:2000, Millipore) and mouse anti-FOXA2 (1:500, Santa Cruz). Secondary antibodies (Alexa series purchased from Life Technologies) were incubated for 1h at room temperature in a blocking solution. Coverslips were mounted in Mowiol (Sigma) and imaged on an upright Nikon A1R confocal microscope equipped with a DIC N2 20X lens NA 0.75. Z-stacks were acquired with a pinhole of 1 Airy unit, a Galvano scanner with line averaging of 2, image size of 1024 x 1024 pixels and step intervals of 0.5 µm. Nuclear markers were quantified using QuPath (Bankhead P. et al., 2017; Figure 6J).

*Dopaminergic neuron differentiation (Arenas lab)*

Cell Culture

Dopaminergic neurons were differentiated by seeding KOLF2.1J cells at a density of 200,000 cells/cm^2^ and differentiating until day 28 as described (Kim et al. 2021), with the exception of SHH C25II, which was substituted by 2 µM purmorphamine. Briefly, wells were coated with Geltrex (Life Technologies) and cells were cultivated in Neurobasal/N2/B27 (Life Technologies) medium with 2mM L-glutamine (Invitrogen), 250 nM LDN193189 (Stemgent), 10 µM SB431542, 2 µM purmorphamine, 0.7µM CHIR99021, and 10 µM Y27632 (Tocris Bioscience) on day 0 of differentiation and were cultured to day 3 after the removal of Y27632 from day 1. 7.5µM CHIR9901 was supplemented from day 4 and LDN, SB and purmorphamine were withdrawn from day 7. On day 10, cells were shifted to Neurobasal/B27/L-Glu supplemented with 20 ng/mL brain-derived neurotrophic factor (BDNF, R&D systems), 20 ng/ml glial cell line-derived neurotrophic factor (GDNF, R&D systems), 1 ng/ml transforming growth factor type b3 (TGFb3, R&D systems), 0.2 mM ascorbic acid (Sigma), 0.2 mM dibutyryl cAMP (Sigma), and 3 µM CHIR99021. On day 11, cells were dissociated into single cells, and were replated onto plates coated with 15 mg/ml polyornithine,1 mg/ml laminin 111, and 2 mg/ml fibronectin at a density of 500,000 cells/cm^2^. The mDA differentiation media without CHIR99021 was applied for cell culture and was supplemented with 10 mM DAPT (Sigma) from day 12. On day 15, cells were replated using the same procedure and cultured until day 25 for characterizations.

Immunostaining

Cells were fixed by 4% paraformaldehyde (PFA) for 30 min at 4˚C, pre-incubated with 5% donkey serum in PBS with 0.1% Triton X-100 (PBST) for 1 hr, incubated with primary antibodies goat anti-FOXA2 (1:1000, R&D Systems), rabbit anti-LMX1 (1:2000, Millipore), mouse anti-Nurr1 (1:1000, Perseus Proteomics), or rabbit anti-TH (1:1000, Millipore) at 4˚C overnight, followed by Alexa488, Alexa555 or Alexa647-conjugated secondary antibodies (Invitrogen) for 1 hr and then 4’,6-diamidino-2-phenylindole (DAPI) for 15 min. Images were visualized in a Zeiss LSM 980 Airyscan microscope (Figure S6L).

*Midbrain organoid differentiation (Ahfeldt lab)*

Cell culture

For midbrain organoid differentiation, KOLF2.1J cells were cultured in Stemflex^TM^ medium (ThermoFisher) on Geltrex-coated under conditions of 37^o^C, 5% CO_2_ in a humidified incubator. Midbrain organoids were differentiated as detailed in (Sarrafha et al. 2021). Briefly, 40x10^6^ iPSCs were seeded in 125-ml disposable spinner flasks (Corning, VWR) in Stemflex + 10µM Rhok inhibitor Y-27632. Flasks were placed on a nine-position stir plate (Dura-Mag) at a speed of 65 rpm. Once spheres reached a size of 300-500µm differentiation was initiated by dual-SMAD inhibition with SB431542 (R&D Systems, 10 μM), LDN193189 (Stemgent, 100 nM), B27-Vit A, and N2 in DMEM-F12. Patterning media was supplemented with CHIR99021 (Stemgent, 3 μM), Purmorphamine (STEMCELL, 2 μM), and SAG (Abcam, 1 μM) for the midbrain-specific patterning (Kriks et al., 2011). Starting day 12, post patterning neural maturation media was composed by DMEM F12 supplemented with N2, B27-VitA, 20 ng/mL GDNF (R&D Systems), 20 ng/mL BDNF (R&D Systems), 0.2 mM ascorbic acid (Sigma), 0.1 mM dibutyryl cAMP (Biolong), 10 μM DAPT (Cayman Chemical). After day 35, spheres were transferred to ultralow attachment plates (Corning, VWR) for long-term cultures in DMEM F12 medium containing N2, B27-VitA, 10 ng/ML GDNF (R&D Systems), 10 ng/mL BDNF (R&D Systems), 0.2 mM ascorbic acid (Sigma).

Immunostaining

For organoid immunohistochemistry (IHC) analysis, at indicated time points midbrain organoids were washed in PBS and fixed with 4% PFA overnight. Fixed organoids were embedded in paraffin and serial sections (4-6 μm) were prepared using a Leica RM2255 microtome. Sections were placed on charged slides and baked overnight at 70°C and IHC was performed on Ventana Benchmark XT. Antigen retrieval with CC1 (citric acid buffer) was performed for 1 h, followed by incubation for 30 min in primary antibodies mouse anti-TH (1:500, EMD Millipore) rabbit anti-NURR1 (1:200, Millipore Sigma), and mouse anti-GFAP (1:10, Millipore). A multimer secondary antibody was used for all samples. IHC sections were imaged using an Aperio VERSA 8 digital slide scanner (Leica Biosystems, Wetzlar Germany; Figure S6M).

Calatayud, Carles, Esther Not Provided Muñoz-Pedrazo, Sandra Not Provided Fernández-Gallego, and Patrik Not Provided Verstreken. n.d. “Modular Generation of Cortical, Striatal and Ventral Midbrain Progenitor Cells v1.” *Protocols.io*. https://doi.org/[10.17504/protocols.io.btmsnk6e](http://dx.doi.org/10.17504/protocols.io.btmsnk6e).

Chal, Jérome, Ziad Al Tanoury, Marie Hestin, Bénédicte Gobert, Suvi Aivio, Aurore Hick, Thomas Cherrier, Alexander P. Nesmith, Kevin K. Parker, and Olivier Pourquié. 2016. “Generation of Human Muscle Fibers and Satellite-like Cells from Human Pluripotent Stem Cells in Vitro.” *Nature Protocols* 11 (10): 1833–50.

Chen, Yu, Carlos A. Tristan, Lu Chen, Vukasin M. Jovanovic, Claire Malley, Pei-Hsuan Chu, Seungmi Ryu, et al. 2021. “A Versatile Polypharmacology Platform Promotes Cytoprotection and Viability of Human Pluripotent and Differentiated Cells.” *Nature Methods* 18 (5): 528–41.

Coyne, Alyssa N., Benjamin L. Zaepfel, Lindsey Hayes, Boris Fitchman, Yuval Salzberg, En-Ching Luo, Kelly Bowen, et al. 2020. “G4C2 Repeat RNA Initiates a POM121-Mediated Reduction in Specific Nucleoporins in C9orf72 ALS/FTD.” *Neuron*. https://doi.org/[10.1016/j.neuron.2020.06.027](http://dx.doi.org/10.1016/j.neuron.2020.06.027).

Dalby, Brian, Sharon Cates, Adam Harris, Elise C. Ohki, Mary L. Tilkins, Paul J. Price, and Valentina C. Ciccarone. 2004. “Advanced Transfection with Lipofectamine 2000 Reagent: Primary Neurons, siRNA, and High-Throughput Applications.” *Methods*  33 (2): 95–103.

Faria Zafar, Alejandra Torres, Laurin Heinrich, Jasmine Singh, Cassandra Hempel, Oliver Braubach, Birgitt Schuele. 2022. “Protocol for CODEX Fixation Steps and Primary Antibody Staining for Induced Pluripotent Stem Cell-Derived Neurons.” https://doi.org/[10.17504/protocols.io.b3fhqjj6](http://dx.doi.org/10.17504/protocols.io.b3fhqjj6).

[Fernandopulle, Michael S., Ryan Prestil, Christopher Grunseich, Chao Wang, Li Gan, and Michael E. Ward. 2018. “Transcription Factor-Mediated Differentiation of Human iPSCs into Neurons.” *Current Protocols in Cell Biology / Editorial Board, Juan S. Bonifacino ... [et Al.]* 79 (1): e51.](http://paperpile.com/b/1MawSz/exQJ2)

Gantner, Carlos W., Cameron P. J. Hunt, Jonathan C. Niclis, Vanessa Penna, Stuart J. McDougall, Lachlan H. Thompson, and Clare L. Parish. 2021. “FGF-MAPK Signaling Regulates Human Deep-Layer Corticogenesis.” *Stem Cell Reports* 16 (5): 1262–75.

Heinrich, Laurin, Faria Zafar, C. Alejandra Morato Torres, Jasmine Singh, Anum Khan, Max Yang Chen, Cassandra Hempel, et al. 2022. “Multiplex Imaging of Human Induced Pluripotent Stem Cell-Derived Neurons with CO-Detection by indEXing (CODEX) Technology.” *Journal of Neuroscience Methods* 378 (August): 109653.

Hildebrandt, Matthew R., Miriam S. Reuter, Wei Wei, Naeimeh Tayebi, Jiajie Liu, Sazia Sharmin, Jaap Mulder, et al. 2019. “Precision Health Resource of Control iPSC Lines for Versatile Multilineage Differentiation.” *Stem Cell Reports* 13 (6): 1126–41.

Howe, Bradley, Ayesha Umrigar, and Fern Tsien. 2014. “Chromosome Preparation From Cultured Cells.” *Journal of Visualized Experiments*. https://doi.org/[10.3791/50203](http://dx.doi.org/10.3791/50203).

Khan, Themasap A., Omer Revah, Aaron Gordon, Se-Jin Yoon, Anna K. Krawisz, Carleton Goold, Yishan Sun, et al. 2020. “Neuronal Defects in a Human Cellular Model of 22q11.2 Deletion Syndrome.” *Nature Medicine* 26 (12): 1888–98.

Kim, Tae Wan, Jinghua Piao, So Yeon Koo, Sonja Kriks, Sun Young Chung, Doron Betel, Nicholas D. Socci, et al. 2021. “Biphasic Activation of WNT Signaling Facilitates the Derivation of Midbrain Dopamine Neurons from hESCs for Translational Use.” *Cell Stem Cell* 28 (2): 343–55.e5.

Kriks, Sonja, Jae-Won Shim, Jinghua Piao, Yosif M. Ganat, Dustin R. Wakeman, Zhong Xie, Luis Carrillo-Reid, et al. 2011. “Dopamine Neurons Derived from Human ES Cells Efficiently Engraft in Animal Models of Parkinson’s Disease.” *Nature* 480 (7378): 547–51.

Martinez, Fernando J., Gabriel A. Pratt, Eric L. Van Nostrand, Ranjan Batra, Stephanie C. Huelga, Katannya Kapeli, Peter Freese, et al. 2016. “Protein-RNA Networks Regulated by Normal and ALS-Associated Mutant HNRNPA2B1 in the Nervous System.” *Neuron* 92 (4): 780–95.

Meijer, Marieke, Kristina Rehbach, Jessie W. Brunner, Jessica A. Classen, Hanna C. A. Lammertse, Lola A. van Linge, Desiree Schut, et al. 2019. “A Single-Cell Model for Synaptic Transmission and Plasticity in Human iPSC-Derived Neurons.” *Cell Reports* 27 (7): 2199–2211.e6.

Moya, Noel, Josh Cutts, Terry Gaasterland, Karl Willert, and David A. Brafman. 2014. “Endogenous WNT Signaling Regulates hPSC-Derived Neural Progenitor Cell Heterogeneity and Specifies Their Regional Identity.” *Stem Cell Reports*. https://doi.org/[10.1016/j.stemcr.2014.10.004](http://dx.doi.org/10.1016/j.stemcr.2014.10.004).

Ray, F. Andrew, Erin Zimmerman, Bruce Robinson, Michael N. Cornforth, Joel S. Bedford, Edwin H. Goodwin, and Susan M. Bailey. 2013. “Directional Genomic Hybridization for Chromosomal Inversion Discovery and Detection.” *Chromosome Research: An International Journal on the Molecular, Supramolecular and Evolutionary Aspects of Chromosome Biology* 21 (2): 165–74.

Sarrafha, Lily, Gustavo M. Parfitt, Ricardo Reyes, Camille Goldman, Elena Coccia, Tatyana Kareva, and Tim Ahfeldt. 2021. “High-Throughput Generation of Midbrain Dopaminergic Neuron Organoids from Reporter Human Pluripotent Stem Cells.” *STAR Protocols* 2 (2): 100463.

Shi, Yichen, Peter Kirwan, and Frederick J. Livesey. 2012. “Directed Differentiation of Human Pluripotent Stem Cells to Cerebral Cortex Neurons and Neural Networks.” *Nature Protocols* 7 (10): 1836–46.

Srinivasaraghavan, Vasavi, Faria Zafar, and Birgitt Schuele. 2022. “Gene Expression Analysis in Stem Cell-Derived Cortical Neuronal Cultures Using Multi-Well SYBR Green Quantitative PCR Arrays.” *BIO-PROTOCOL*. https://doi.org/[10.21769/bioprotoc.4283](http://dx.doi.org/10.21769/bioprotoc.4283).

Surmacz, Beata, Heather Fox, Alex Gutteridge, Paul Fish, Sandra Lubitz, and Paul Whiting. 2012. “Directing Differentiation of Human Embryonic Stem Cells toward Anterior Neural Ectoderm Using Small Molecules.” *Stem Cells* 30 (9): 1875–84.

Tcw, Julia, Minghui Wang, Anna A. Pimenova, Kathryn R. Bowles, Brigham J. Hartley, Emre Lacin, Saima I. Machlovi, et al. 2017. “An Efficient Platform for Astrocyte Differentiation from Human Induced Pluripotent Stem Cells.” *Stem Cell Reports* 9 (2): 600–614.

Tian, Ruilin, Mariam A. Gachechiladze, Connor H. Ludwig, Matthew T. Laurie, Jason Y. Hong, Diane Nathaniel, Anika V. Prabhu, et al. 2019. “CRISPR Interference-Based Platform for Multimodal Genetic Screens in Human iPSC-Derived Neurons.” *Neuron* 104 (2): 239–55.e12.
